# Integrative Proteomic Analysis Implicates Inhibition of Intracellular Protein Trafficking in Therapy-Induced Migrastasis in Prostate Cancer

**DOI:** 10.64898/2026.07.02.736165

**Authors:** Weining Chen, Saadyeh Rashidi, Henry C.-H. Law, Fangfang Qiao, Johnny W. Zigmond, Katelyn L. O’Neill, Nicholas T. Woods, Chittibabu Guda, Raymond Bergan

## Abstract

**Background:** Dysregulated cell migration leading to metastasis remains the primary cause of cancer-related mortality. It has been challenging to understand how cells regulate migration. We have previously created the first selective inhibitor of cell migration, KBU2046. Here, we use it as a probe to identify regulatory processes.

**Methods:** Metastatic and primary human prostate cancer cells were treated for different times and at different concentrations with KBU2046. Immunofluorescent microscopy examined protein localization in cells. Label-free mass spectrometry (MS) was performed on total cell proteins, Tandem Mass Tag (TMT) labeling MS was used on membrane fractions, and temporal phosphoproteomic profiling. Results were analyzed with a suite of bioinformatic tools.

**Results:** KBU2046-induced migrastasis is associated with the accumulation of activated integrin β1 into focal adhesions. Whole-cell proteomics demonstrated suppression of processes that mediate intracellular protein trafficking and increases in mitochondrial energy-generation signatures. Evaluation of the membrane fraction identified increases in membrane repair and maintenance processes and decreases in those that drive motility. Temporal– and concentration-dependent phosphoproteomic profiling revealed that KBU2046 initiates a dynamic, cascading sequence of transient signaling waves rather than a static block.

**Conclusions:** KBU2046-induced migrastasis appears to operate through spatial decoupling rather than structural degradation. By restricting the intracellular trafficking machinery required for receptor recycling, KBU2046 limits focal adhesion turnover, providing a correlative framework to inhibit metastatic dissemination independent of direct cytotoxicity.

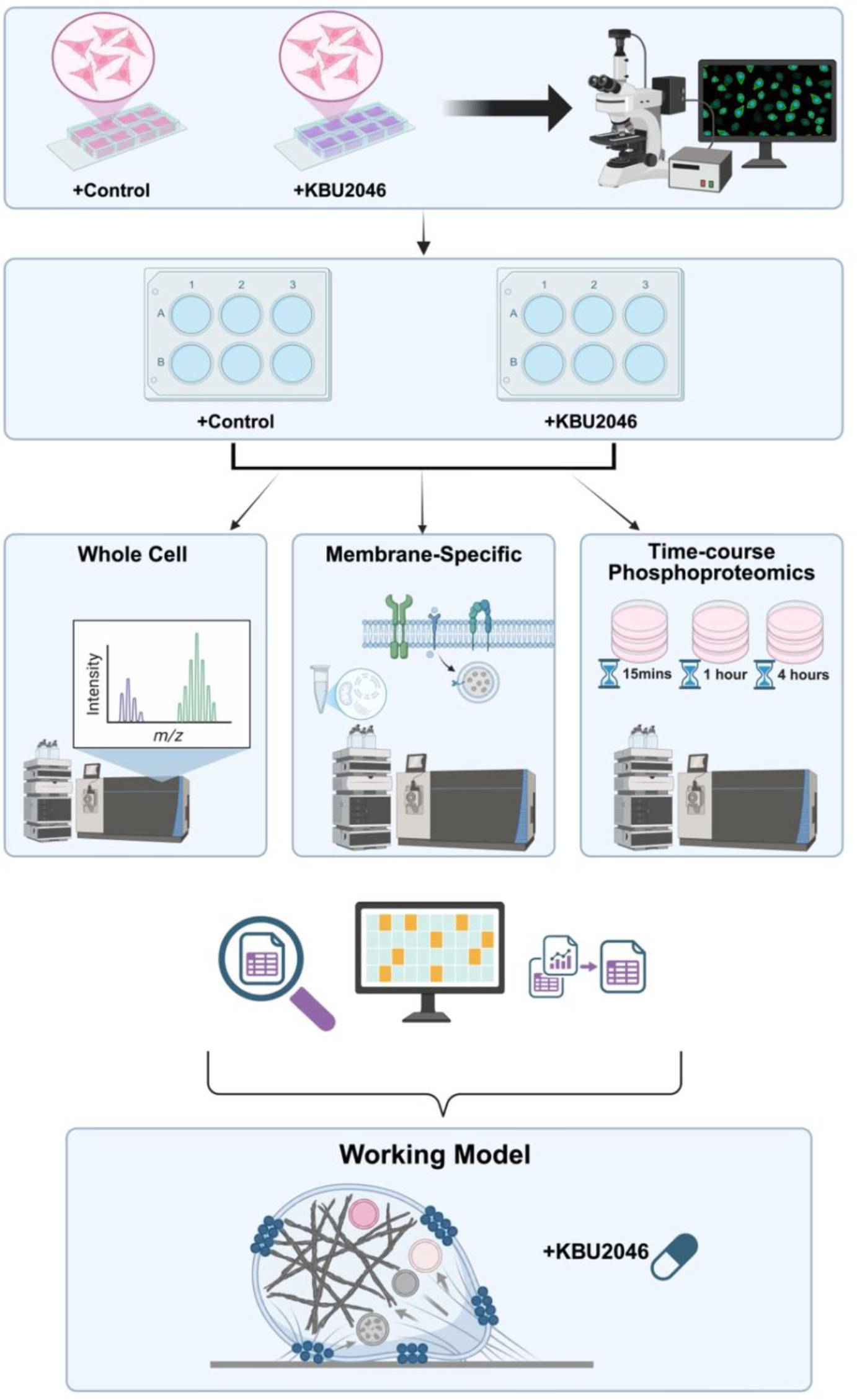
Graphic Abstract.

## 1. Introduction

Metastasis remains the primary cause of cancer-related mortality and presents a therapeutic challenge once the disease extends beyond the primary organ [1, 2]. Once distant metastasis has developed, systemic interventions constitute the mainstay for treatment of advanced malignancy. While there is an expanding array of chemotherapeutic, hormonal, radiopharmaceutical, and immunologic agents, their unifying effect on cancer cells relates to cytostasis and/or cytotoxicity.

Cancer metastasis constitutes a complex multistage process, with cellular motility playing a key role throughout progression [2–4]. Cell movement across a surface is termed migration. This fundamental process underlies cell invasion, which couples migration to protein degradation, allowing cells to move through otherwise intact organs. Migration also underlies metastasis, which requires migration and invasion for cells to transit multiple steps in the metastatic cascade. Selectively inhibiting cell migration, a concept termed migrastasis [5–7], offers a rational therapeutic strategy to disrupt dissemination. However, it has been a challenge to therapeutically target the induction of migrastasis. This is because while a large number of cellular processes and pathways alter migration, they lack specificity in their effects.

By coupling cheminformatics to tiered selection and deselection strategies designed to selectively perturb cellular processes, we have identified the first selective migrastatic agent, KBU2046 [8]. KBU2046 is a small molecule, and it induces migrastasis in several cancer types. Extensive studies have been performed on human prostate cancer (PCa), which demonstrate that KBU2046-mediated induction of migrastasis in turn inhibits cell invasion as well as the formation of metastasis, with the latter shown in several clinically relevant murine models. An associated comprehensive series of cellular and systemic studies demonstrates a lack of toxicity and has not identified other off-target effects [8].

However, the molecular mechanism by which these selective functional effects are induced remains unclear. In silico modeling indicates that KBU2046 binds selectively to the cleft formed by Heat Shock Protein 90 beta (HSP90β) and Cell Division Cycle 37 (CDC37), distinguishing it from agents that block the general ATPase domain of HSP90β. Inhibitors of HSP90β ATPase activity broadly disrupt many cell functions and induce cytotoxicity; KBU2046 has none of these effects [8]. Biochemical evaluations do demonstrate that KBU2046 selectively inhibits binding of Raf1 to the HSP90β/CDC37 heterocomplex, potentially inhibiting the activation of Raf1 and subsequent regulation of migration [8].

It is important to consider that these foundational molecular interactions are not confirmed, and even if they hold true, the downstream intracellular networks mediating the biological response remain undefined. Based on its unique and selective functional effects, KBU2046 represents a powerful biological probe that can be used to provide insight into how cells regulate migration and how migrastasis can be selectively induced therapeutically. By integrating whole-cell proteomic profiling to comprehensively inventory protein abundance, spatial membrane proteomics to quantify the dynamic surfaceome, and temporal and concentration-dependent phosphoproteomics to track the progression of signaling cascades, we characterized the fundamental cellular responses to therapy-induced migrastasis. Collectively, these multiplexed proteomic datasets establish a framework that elucidates how precision agents can decouple spatial cellular movement from basal biological viability.

## 2. Materials and Methods

### 2.1. Cell Culture

The human prostate cancer PC3 cell line was obtained from the American Type Culture Collection (ATCC, Manassas, VA). The 1532CPTX cancer cell line was generated from micro-dissected primary tumor from the patient. These HPV-transformed primary cell lines were a generous gift from S. Topalian (National Cancer Institute, Bethesda, MD), and have been characterized by us and others [8–11]. PC3 cells were cultured in RPMI 1640 (Thermo Scientific, Waltham, MA), 1532CPTX cells in keratinocyte-serum-free medium supplemented with recombinant epidermal growth factor and bovine pituitary extract (Gibco, Grand Island, NY), both along with Antibiotic-Antimycotic (cat #s: 15240112 and 15240062, respectively; Gibco), and in 10% and 3% fetal bovine serum (FBS; Gibco, Grand Island, NY) for PC3 and 1532CPTX cells, respectively, as previously described by us [8–11]. Cells were maintained under sub-confluent growth conditions, maintained in standard humidified conditions at 37 °C and 5% CO2, replenished from stocks every two months, routinely monitored for mycoplasma, and authenticated, as previously described by us [8, 12].

### 2.2 Antibodies and Reagents

The following primary and secondary antibodies were used: anti-Integrin β1 primary antibody, clone 12G10 (MAB2247; Sigma Aldrich, St. Louis, MO), and Anti-Mouse IgG (H+L) Highly Cross-Adsorbed Secondary Antibody (A32766TR; Invitrogen, Carlsbad, CA). Rhodamine Phalloidin (R415; Invitrogen, Carlsbad, CA) was used for filamentous actin visualization.

Chemicals, biochemical reagents, and select consumables included KBU2046, dimethyl sulfoxide (MilliporeSigma, Burlington, MA), 4% paraformaldehyde in phosphate buffered saline (PBS) (J61899.AK; Thermo Scientific, Waltham, MA), PBS (14190250; Gibco, Grand Island, NY), Triton X-100 (T8787 250ML; MilliporeSigma, Burlington, MA), bovine serum albumin (BSA) (BP1600 100; Fisher Scientific, Hampton, NH), glycine (MilliporeSigma, Burlington, MA), ProLong Gold Antifade Mountant with DAPI (P36931; Invitrogen, Carlsbad, CA), 8-well chamber slides (80841; Ibidi, Gräfelfing, Germany), removable coverslips (10811; Fisher Scientific, Hampton, NH), RIPA buffer (89900; Thermo Scientific, Waltham, MA), Mem PER Plus Membrane Protein Extraction Kit (89842; Thermo Scientific, Waltham, MA), Halt Protease and Phosphatase Inhibitor Single Use Cocktail (100X) (78442; Thermo Scientific, Waltham, MA), PMSF (MilliporeSigma, Burlington, MA), sodium orthovanadate (J60191.AD; Thermo Scientific Chemicals, Waltham, MA), sodium fluoride (S6776 100G; MilliporeSigma, Burlington, MA), beta glycerophosphate (G9422 10G; MilliporeSigma, Burlington, MA), BCA protein assay kit (A65453; Thermo Scientific, Waltham, MA), Tandem Mass Tag reagents (Thermo Fisher Scientific, Waltham, MA), and trypsin (Promega, Madison, WI).

### 2.3. Pharmacological Treatment

To characterize the phenotypic response, cells were treated with 10 µM KBU2046, 1 µM KBU2046, or a corresponding dimethyl sulfoxide (DMSO) vehicle control. For whole-cell proteomic profiling and membrane-enriched proteomics, cells were subjected to a prolonged 72-hour exposure, as denoted. For time-resolved phosphoproteomic profiling was performed by harvesting cells at 15-minute, 1-hour, and 4-hour time points post-treatment, as denoted.

### 2.4. Time-Course Immunofluorescence Microscopy

To define the temporal dynamics of the kinetic arrest phenotype, cells were seeded into 8-well chamber slides at a density of 1.5×10^4^ cells for the PC3 and 1×10^4^ cells for the 1532CPTX. Twenty-four hours after attachment, cells were treated with 10 µM KBU2046. Treatment durations were 2 hours for the 1532CPTX cells and 3 hours for the PC3 cells. At designated time points, cells were fixed in 4% paraformaldehyde in PBS for 10 minutes at room temperature. Following fixation, cells were washed three times in PBS and permeabilized with 0.1% Triton X-100 in PBS for 10 minutes. Nonspecific binding sites were blocked by incubating the chambers in a solution containing 50 mM glycine and 1% BSA in PBS for 1 hour at room temperature.

Cells were then incubated overnight at 4°C with the anti-Integrin β1 primary antibody diluted 1:100 in 0.1% BSA in PBS. Following primary incubation, cells were washed and incubated with the Anti-Mouse IgG (H+L) Highly Cross-Adsorbed Secondary Antibody diluted 1:1000 in 0.1% BSA in PBS for 1 hour at room temperature in the dark. The actin cytoskeletal architecture was visualized by co-incubating with Rhodamine Phalloidin at a 1:400 dilution. To simultaneously counterstain the nuclei and seal the preparations, the chambers were fitted with removable coverslips and mounted using ProLong Gold Antifade Mountant with DAPI. High-resolution image acquisition was performed using a Zeiss LSM 800 microscope equipped with an Airyscan detector (Carl Zeiss Microscopy, Jena, Germany), with support from the University of Nebraska Medical Center – UNMC Advanced Microscopy Core Facility (the core), RRID: SCR_022467. Z-stack reconstruction was used to assess the preservation of cell spreading and the morphological integrity of the actin cytoskeleton. Imaging was captured at 20X magnification for 1532CPTX cells and 63X magnification with a 2X zoom for PC3 cells to resolve fine receptor distribution patterns.

### 2.5. Proteomic Sample Preparation and Bioinformatics Overview

A multi-tiered proteomic strategy was employed to comprehensively map the cellular mechanism of action. This approach integrated whole-cell label-free proteomics to assess overall stoichiometric stability, membrane-enriched subcellular TMT-labeled proteomics to resolve spatial dynamics at the cell periphery, and quantitative temporal TMT-labeled phosphoproteomics to capture evolving regulatory signaling cascades. Extensive details regarding cellular extraction protocols, mass spectrometry data acquisition parameters, and comprehensive bioinformatic and statistical analysis pipelines are provided in the **Supplementary Materials and Methods.**

## 3. Results

### 3.1. Phenotypic Anchoring: KBU2046 Induces Spatial Alterations in Activated Integrin β1

Previous work by us demonstrated that KBU2046 selectively inhibits cell migration in human cancer cells across several types of cancer, including prostate cancer (PCa) [8]. However, the mechanisms mediating this effect remain undefined. Cellular migration involves the physical translocation of a cell across a surface, a requirement for invasion, and for the ultimate development of metastasis [13]. This ongoing process relies on the dynamic assembly of focal complexes at the leading edge of the cell to generate physical traction against the external substrate [14–18]. Integrins function as the primary transmembrane receptors mediating this physical adhesion and form the core components of these focal complexes, as well as the resultant mature focal adhesions [14, 15, 19]. We hypothesized that if KBU2046 inhibits the regulation of cell migration-associated machinery, then observable effects on integrins should be evident. We focused our investigation on Integrin β1, a major component of PCa cell focal adhesions, facilitator of migration, and critical mediator of metastasis to the bone, which serves as a primary site of PCa dissemination [20–22]. Interrogation of the activated conformation of integrin β1 allows assessment of the functional machinery responsible for adhesion, and we thus evaluated this by immunofluorescence staining using an antibody that recognizes activated integrin β1 [23, 24].

Studies first examined 1532CPTX cells, a previously characterized human PCa primary cell line that exhibits relatively well-defined architectural features [8–11]. Immunofluorescent staining for activated integrin β1 and actin demonstrates a prototypical staining pattern for an epithelial cell (**Figure 1A**). Cells exhibit a polarized pattern reflecting their movement in a given direction across a surface. Filamentous actin extensions are observed at the leading edge of the cell, demonstrating the formation of lamellipodia. Within lamellipodia, fine punctate collections of activated integrin β1 begin to form, indicative of nascent focal adhesions. Moving rearward in the cell, collections of activated integrin β1 increase in size and are concentrated in the center of the cell, reflecting mature focal adhesions that serve to anchor the cell. Finally, they disappear at the rear of the cell, reflecting dissolution of focal adhesions, thus allowing the cell’s lagging edge to detach and the cell to move forward. In contrast, upon KBU2046 treatment, 1532CPTX cells lose directional polarization and exhibit actin extensions in opposing directions. Strikingly, activated integrin β1 is now present in large, clustered structures, and these tend to be located on opposite sides of the cell. Together, these features give the appearance of a cell that is “stuck” in place.

**Figure 1.**
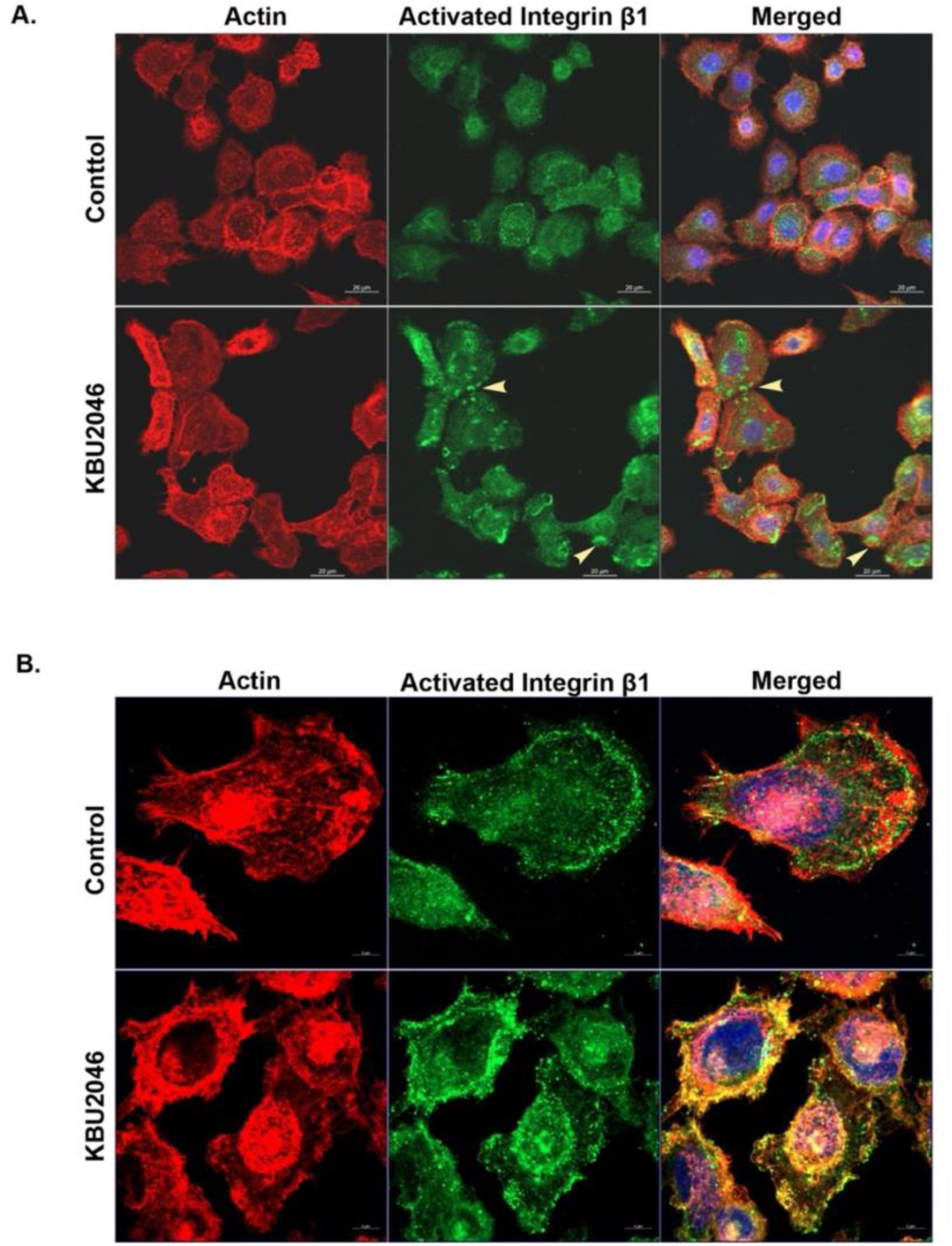
KBU2046 Induces Peripheral Sequestration of Activated Integrin β1. Immunofluorescence imaging of 1532CPTX (**A**) and PC3 (**B**) prostate cancer cells treated for 2 or 3 hrs, respectively, with 10 µM KBU2046 or vehicle (control). The arrow denotes the punctate collections of activated integrin β1. Representative images of cells stained for F-actin (Red) and activated integrin β1 (Green) are depicted from three independent biological replicates.

The morphological features of PC3 cells are much less refined than those of 1532CPTX cells, reflecting their aggressive and metastatic phenotype (**Figure 1B**). Within that context, PC3 cells exhibit a polarized pattern reflecting unidirectional movement, have a much more disorganized actin pattern, but exhibit punctate patterns of activated integrin β1 at the leading edge of the cell. Similarly, with KBU2046 treatment, PC3 cells lose directional polarization, the localization of actin is also less polarized, activated integrin β1 is also present in larger clusters, and these too tend to be located on opposite sides of the cell. Findings from both 1532CPTX and PC3 cells demonstrated that KBU2046 alters the localization of activated integrin β1 and induces associated changes in cell morphology. Both effects are consistent with changes in integrin dynamics. Prior studies demonstrated that KBU2046 was a selective migrastatic agent [8]; current findings corroborate those and extend them by demonstrating modulation of the location of a protein species known to be central to mediating migration.

### 3.2. Global Proteome: Quantitative label-free proteomic analysis of KBU2046-treated PC3 cells

Next, investigations aimed to use KBU2046 as a biological probe to interrogate potential regulatory mechanisms associated with the induction of migrastasis. Some have suggested that migrastasis may result from depletion of core migration components, while others raise the possibility that they are maintained, but there is disruption in function [5–7]. Loss-of-function explanations implicate the dissolution of adhesion complexes. While suggested explanations are all logical, there is little experimental data in the context of selective induction of migrastasis. For KBU2046-induced migrastasis, immunofluorescence data demonstrate peripheral changes in the localization of activated integrin β1. Further, this data does not indicate loss of protein expression, nor does it indicate loss of adhesion complexes. The latter is further supported by the fact that loss of adhesion complexes leads to cell detachment, but this is not a feature of KBU2046’s action. Together, these considerations support the hypothesis that KBU2046 is not globally depleting core migration components, that it is disrupting function, and the nature of that function is unique in that it has not yet been linked to induction of migrastasis.To examine this hypothesis, studies measured the effect of KBU2046 on protein profiles in PC3 cells by mass spectrometry (MS). Investigations focused on PC3 cells because prior studies have extensively characterized the treatment effects of KBU2046 on them, demonstrating that induction of migrastasis leads to downstream inhibition of cell invasion and metastasis; the latter was shown across multiple systemic models, with separate studies demonstrating lack of toxicity at cellular and systemic levels [8]. PC3 cells were treated with 10 µM KBU2046 for 72 hours, i.e., standard treatment conditions shown to selectively inhibit migration, or vehicle for control cells (n = 3 biological replicates per group), and a label-free global whole-cell proteomic screen by MS-based analytics compared protein abundance between groups (**Figure 2A**). This study identified 4,694 unique proteins and served as the initial dataset.

**Figure 2:**
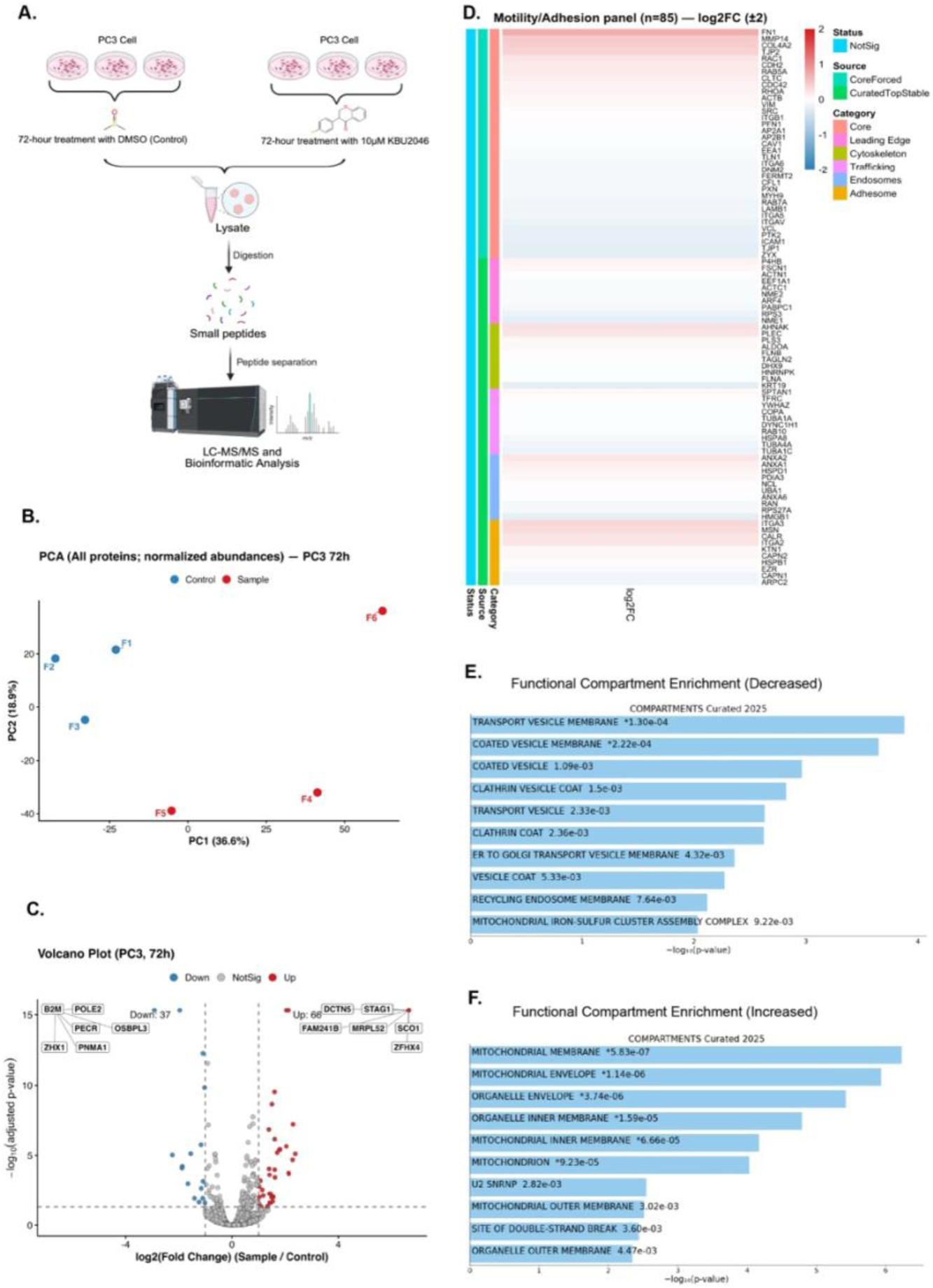
Global Proteomic Stability and Metabolic Reprioritization Under Prolonged KBU2046 Exposure. (**A**) **Whole-Cell Proteomics Workflow.** Schematic of the experimental design for the label-free global proteomics screen. (**B**) **Principal Component Analysis of the Whole Cell Proteome.** Principal Component Analysis was performed on the normalized, Log2(x+1)-transformed abundance values of all detected proteins (n=4,690) across control and KBU2046-treated conditions, with n =3 biological replicates/group. (**C**) **Volcano Plot of Global Differential Abundance.** Data are presented as a scatter plot of change in protein abundance levels, significance thresholds were set at |Log2 FC| ≥1.0 (vertical dashed lines) and adjusted p-value < 0.05 (horizontal dashed line), and colored points represent differentially abundant proteins as denoted. The apparent number of dots in the volcano plot is fewer than the actual number of significant proteins because multiple proteins share identical or highly similar Log2 FC and adjusted p-values, resulting in overlap on the plot. Consequently, several significant hits are superimposed, resulting in fewer visible points than the true number reported in the table. (**D**) **Motility Hardware Heatmap.** Targeted heat map displaying the Log2 FC of key motility, adhesion, and trafficking proteins selected for robust proteomic detection. The color scale is capped at ± 2.0 to standardize visualization. (**E-F**) **Functional Compartment Enrichment**. Bar plots showing the top-enriched spatial compartments identified by a COMPONENTS-curated analysis of proteins exhibiting altered abundance. Compartments are ranked by p-values (shown on each bar), with compartments exhibiting decreased (**E**) and increased (**F**) abundance listed separately.

Principal Component Analysis of the normalized protein abundance values assessed the global variance across the six experimental runs (**Figure 2B**). The plot demonstrates a variance shift between groups, with KBU2046-treated replicates showing more separation along PC1 (36.6%) and PC2 (18.9%) than the relatively clustered vehicle controls, suggesting greater proteomic heterogeneity following treatment. We next applied an adjusted p-value < 0.05 and a fold change (FC) threshold of 2.0, corresponding to |Log2 FC| ≥1.0, to select for changes in protein abundance to be further evaluated. The differential abundance landscape, visualized via a Volcano Plot (**Figure 2C**), shows that only 103 proteins, representing 2.2% of the quantified proteome, exhibited altered abundance. This subset included 66 unique proteins with increased abundance and 37 with decreased abundance. The limited number of modulated targets observed over an extended 72-hour treatment window supports a selective cellular response rather than a broad nonspecific perturbation.

To assess whether core motility– and adhesion-associated machinery were broadly preserved, the 4,694 MS-identified proteins were screened against a curated motility/adhesion protein panel. This panel was constructed by intersecting the dataset with defined gene sets from the Adhesome.org database [25] (232 core adhesome components) and specific MSigDB [26] functional compartments, including the Leading Edge (433 genes), Cytoskeleton (540 genes), Trafficking (630 genes), and Endosomal networks (630 genes). To ensure that the analysis accurately reflected the stability of the core machinery rather than detection artifacts, a robustness-first selection strategy was implemented (**Supplementary Materials and Methods**). Candidates were prioritized based on quantitative reliability, including consistent detection across biological replicates, high normalized abundance, and a low coefficient of variation in control samples (**Supplementary Materials and Methods**). Additionally, a core set of canonical markers, including specific integrins, actins, Ras-related GTPase proteins (Rabs), and clathrin subunits, was incorporated into the panel to enhance comprehensive coverage of the structural apparatus. From 4,694 screened proteins, this led to the identification of 85 curated motility– and adhesion-associated proteins whose relative abundance was visualized using a heatmap (**Figure 2D**). As illustrated by the predominantly neutral signal across the panel, the changes in abundance were remarkably subtle, confined to a narrow, biologically nominal range of –0.28 to +0.68. None of the proteins in this targeted panel satisfied both criteria for differential abundance, i.e., adjusted p-value < 0.05 and |Log2 FC| ≥ 1. Importantly, the preservation among structural targets supports that the abundance of proteins in core migratory remains stable with KBU2046 treatment.

The determination that core motility machinery remains stable in the face of migrastasis supports that there is a blockade in the machinery responsible for executing its function. While this core machinery is stable, the global proteome exhibited targeted changes within the subset of 103 differentially abundant proteins. To probe the role of these differentially abundant proteins in inducing migratory arrest, we evaluated the spatial requirements of cell movement. Evaluating proteomic shifts through a spatial lens is supported by the current biological context. We mapped differentially abundant protein signatures using the COMPARTMENTS Curated database [27], accessed through the Enrichr [28–30] platform. This database integrates comprehensive literature curation with high-throughput experimental evidence to provide specific subcellular resolution.

Interrogation of the protein fraction with decreased abundance using this spatially resolved database identified a targeted decrease in enrichment of the vesicular transport network (**Figure 2E**). Specifically, significant decreases in enrichment in proteins localized to the transport vesicle membrane (adjusted p-value: 0.023) and coated vesicle membrane (adjusted p-value: 0.023) were identified (**Supplementary Table S1 and Supplementary Materials** (**Bioinformatics Raw Data**)). This is relevant to the observed phenotype because vesicular transport, specifically clathrin-mediated endocytosis, serves as a primary mechanism for the internalization and recycling of surface integrins [31–34]. Cellular migration relies on the dynamic assembly and disassembly of focal contacts to generate physical traction [14–19]. The active turnover of these adhesion sites involves the internalization and recycling of surface integrins through endocytic vesicular pathways [35, 36]. The identified decreases in these vesicular transport pathways, along with the observed accumulation of activated integrin β1 clusters in our immunofluorescence studies, suggest that KBU2046 restricts this localized transport machinery.

Interestingly, the protein fraction with increased abundance reveals a coordinated shift in cellular bioenergetics (**Figure 2F**). The spatial mapping indicates a significant increase in enrichment within mitochondrial structures, including the mitochondrial membrane (adjusted p-value: 0.000136), mitochondrial envelope (adjusted p-value: 0.000136), and the organelle envelope (adjusted p-value: 0.000296). Significantly increased enrichment was also observed in the organelle inner membrane (adjusted p-value: 0.000947), mitochondrial inner membrane (adjusted p-value: 0.0032), and the mitochondrion overall (adjusted p-value: 0.0037) (**Supplementary Table S2 and Supplementary Materials** (**Bioinformatics Raw Data**)). Considering that cell migration is an energy-intensive process [37–40], it is unsurprising that changes in energy consumption would be associated with induced changes in migration, as the cell might try to counteract the inhibited movement.

The observation of phenotypic retention of the cell being “stuck” in place following treatment with KBU2046, despite maintaining a stable abundance of the core motility proteins, points toward a mechanism of functional restriction rather than structural degradation. The combination of preserved physical components and targeted attenuation of vesicular transport signatures establishes a distinct spatial profile. To assess whether this transport impairment affects the localization of the stable adhesion machinery, the investigators next analyzed the isolated physical interface of the cell through a targeted membrane-enriched proteomic analysis.

### 3.3. Analysis of changes in the membrane-confined proteome and their association with regulators of cell migration

To investigate changes in protein networks occurring at the cell boundary, Tandem Mass Tag (TMT) protein labeling, using a multiplex strategy, followed by MS proteomic analysis, was conducted on the membrane fraction of PC3 cells (**Figure 3A**). Cells were treated with 10 µM KBU2046 or vehicle (n. = 4 replicates) for 72 hours, and membrane fractions were isolated as described in the **Supplementary Materials and Methods**. Analysis of the membrane-associated compartment using a 1% False Discovery Rate threshold identified 5,715 unique proteins based on peptide spectral matching. Within this group, 5,585 proteins generated consistent reporter ion intensities across all multiplexed samples, providing the numerical abundance values required for relative quantification. It was this group of 5,585 proteins that was subjected to further analysis. Principal component analysis of the normalized abundance values showed that the variance was distributed across PC1 (32.86%) and PC2 (19.71%) (**Figure 3B**). Along the first principal component, the cells treated with KBU2046 separate from the vehicle controls, indicating that the primary variance at the cellular interface is driven by the treatment condition.

**Figure 3.**
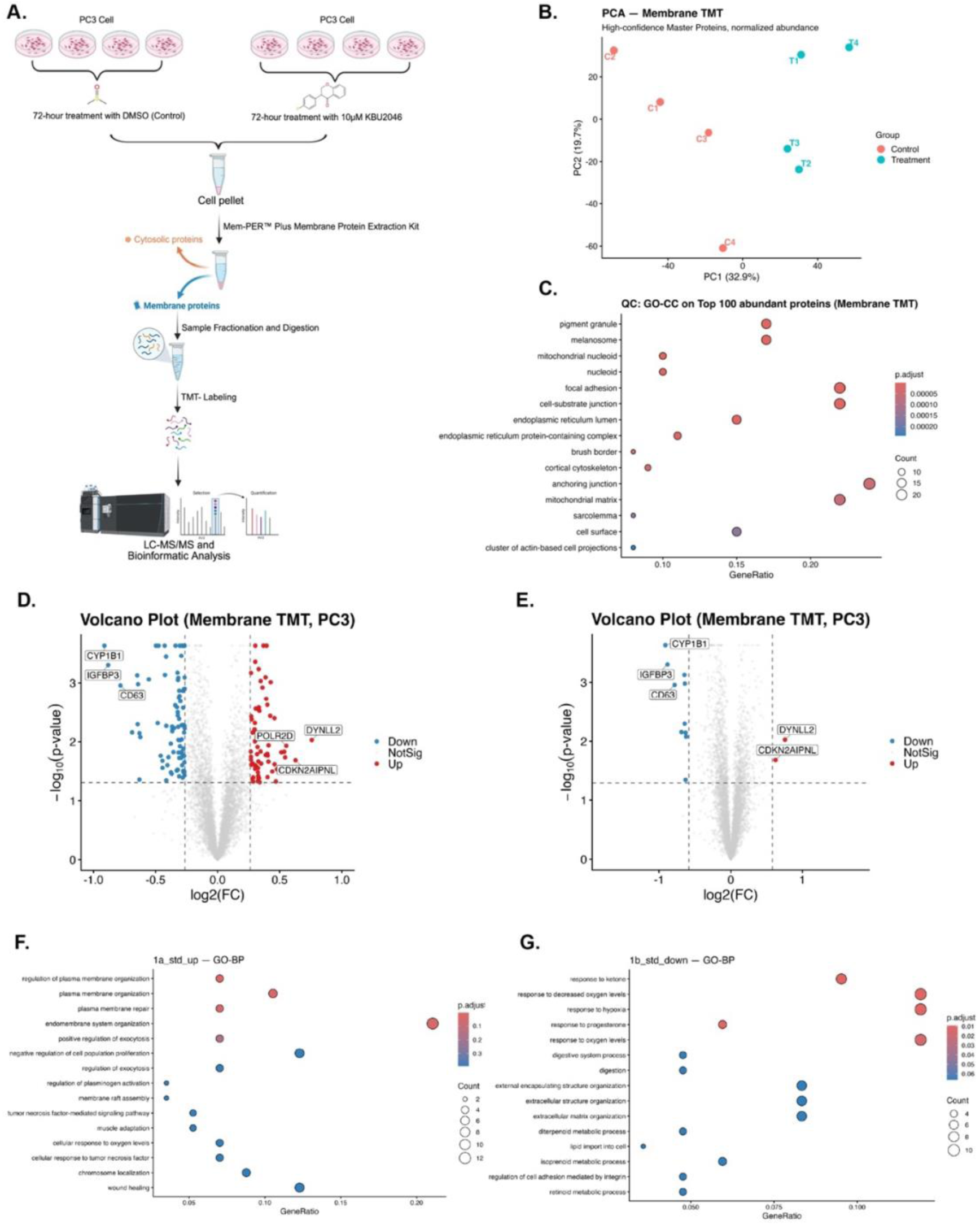
Membrane-enriched Proteomic Profiling Reveals Altered Abundance of Peripheral Trafficking-associated Proteins. (**A**) **Membrane-specific Proteomics Workflow schematic.** Schematic of experimental design. Data are derived from n = 4 independent biological replicates per treatment group. (**B-C**) **Subcellular fractionation purity and global variance.** (**B**) Principal component analysis of the normalized protein abundance values for KBU2046 and control cells. (**C**) Gene Ontology enrichment analysis of the 100 most abundant proteins detected in the whole membrane fraction. (**D-E**) **Differential Membrane Proteomics Volcano Plots.** (**D**) Standard Stringency (|Log2 FC| ≥ 0.26; 1.2-FC; p-value < 0.05). (**E**) High-Stringency (|Log2 FC| ≥ 0.58; 1.5-FC; p-value < 0.05). (**F-G**) **Gene Ontology Enrichment Analysis of Differentially Abundant Membrane Proteins.** Functional enrichment of Biological Process for membrane proteins exhibiting significantly increased (**F**) or decreased (**G**) abundance following KBU2046 treatment (|Log2 FC| ≥ 0.26, p-value <0.05). In these plots, the x-axis represents GeneRatio, the dot size indicates gene count, and the dot color reflects adjusted enrichment significance.

A tiered thresholding strategy was used to account for the ratio compression inherent in the multiplexed analysis of complex lipid matrices [41–43] and to evaluate specific changes. Using a 1.2-fold change cutoff allowed for broad pathway discovery, whereas a high-stringency 1.5-fold change cutoff prioritized robust molecular candidates. To assess the fidelity of our subcellular fractionation, Gene Ontology enrichment analysis of the top 100 most abundant proteins was performed (**Figure 3C**). This analysis validated the high purity of the isolated migratory machinery and revealed a proteomic profile rich in canonical structural and adhesion proteins. The data demonstrated significant enrichment for interface-associated compartments important to cellular migration, specifically highlighting those related to focal adhesion (GO:0005925; adjusted p-value: 4.40×10^-7^, gene count: 22, GeneRatio: 22/100) and cell-substrate junction (GO:0030055; adjusted p-value: 4.43×10^-7^, gene count: 22, GeneRatio: 22/100) networks. Of interest, within these migration-enriched terms the following core mediators were prominently identified (**Supplementary Materials (Bioinformatics Raw Data**)): integrin β1 (GeneID: ITGB1), vimentin (GeneID: VIM), plectin (GeneID: PLEC), filamin A (GeneID: FLNA), filamin B (GeneID: FLNB), myosin heavy chain 9 (GeneID: MYH9), actin beta (GeneID: ACTB), and actinin alpha 1 (GeneID: ACTN1). In addition to focal adhesion and cell-substrate junction gene ontology terms, the broader enrichment profile predominantly features additional terms related to cellular migration and cytoskeletal organization, such as the anchoring junction (GO:0070161; adjusted p-value: 4.32×10^-5^, protein count: 24), cortical cytoskeleton (GO:0030863; adjusted p-value: 4.32×10^-5^, protein count:9) and the actin cytoskeleton (GO:0015629; adjusted p-value: 3.31×10^-3^, protein count: 13). This indicates that even among categories distributed throughout the statistical ranking, the cell migration related pathways remain a predominant functional signature. The robust detection of these interconnected networks links the quality-control evaluation to the biological mechanism, as the prominent abundance of these specific structural markers confirms that the membrane-enriched preparation successfully captured the physical interface of the cell.

Volcano plots were used to visualize global shifts in the membrane-associated proteome (**Figure 3D and E**). The analysis detailed the differential abundance of specific membrane-associated proteins under a standard threshold (|Log2 FC| ≥ 0.26, 1.2-FC), identifying 66 targets with increased abundance and 87 targets with decreased abundance (**Figure 3D**). Applying a high Stringency Threshold (|Log2 FC| ≥ 0.58, 1.5-FC) prioritized a restricted core set of 2 targets with increased abundance and 10 targets with decreased abundance (**Figure 3E**). The asymmetry of this signature suggests that the primary mechanism of action involves the targeted reduction in abundance or internalization of specific membrane-associated proteins, rather than a compensatory increase in the abundance of new surface markers.

While mapping changes in protein abundance provides structural context, understanding the functional consequences of these shifts requires evaluating the associated biological processes. We therefore used Gene Ontology Biological Process enrichment analysis to examine these changes. Within the standard threshold dataset, the fraction with increased abundance revealed enrichment of pathways associated with active structural maintenance, including plasma membrane organization (GO:0007009; adjusted p-value: 0.033, gene count: 6, GeneRatio: 6/57) and plasma membrane repair (GO:0001778; adjusted p-value: 0.033, gene count: 4, GeneRatio: 4/57) (**Figure 3F**). These specific biological pathways encompass the molecular networks required to maintain boundary integrity and dynamically remodel the membrane architecture [44]. Interestingly, the fraction with decreased abundance showed decreased enrichment of the response to hypoxia (GO:0001666; adjusted p-value: 0.009, gene count: 10, GeneRatio: 10/84) and the response to progesterone (GO:0032570; adjusted p-value: 0.009, gene count: 5, GeneRatio:5/84) (**Figure 3G**). Given that hypoxia-associated signaling and hormonal responses often regulate pathways that facilitate cellular invasion [45, 46], the decreased enrichment of these processes offers a logical explanation for the structural modifications observed within these membrane networks. For the high-stringency proteins, 10 had decreased abundance, and 2 had increased abundance. These small input sizes precluded Gene Ontology enrichment algorithms from attaining reliable network inferences. Considering the above findings for the 66 proteins with increased abundance and 87 proteins with decreased abundance together, it outlines a framework wherein therapy-induced migrastasis is associated with an increase in membrane repair and maintenance, and a decrease in processes that drive motility.

While the proteomic analyses performed above together characterized protein abundance across the whole cell and then within the cell membrane, the dynamic progression of events governing these shifts remains undetermined. The downregulation of surface proteins in particular typically takes hours [47], whereas the primary pharmacological interaction occurs on a significantly faster timescale [48, 49]. To examine the initial signaling cascades driving this physical reorganization, we conducted a high-resolution time-course phosphoproteomics analysis.

### 3.4. Quantitative Temporal Phosphoproteomics: Temporal Deconvolution of the Mechanism

To probe the biochemical mechanism of KBU2046, a time– and concentration-dependent MS-based phosphoproteomic analysis of TMT-labeled proteins was performed. As schematically presented (**Figure 4A**), PC3 cells were treated with 1 or 10 μM KBU2046, or vehicle for controls, for 15 minutes, 1 hour, or 4 hours, giving 9 different conditions, each performed in replicates of n = 3. This targeted analysis identified deep proteomic coverage, quantifying 17,104 unique phosphopeptides across 3,785 unique phosphoproteins. Principal component analysis of the quantified phosphorylation sites indicated that the variance was distributed across PC1 (21.2%) and PC2 (19.3%) (**Figure 4B**). The sample distribution reflects the experimental separation across the different time intervals and treatment concentrations.

**Figure 4:**
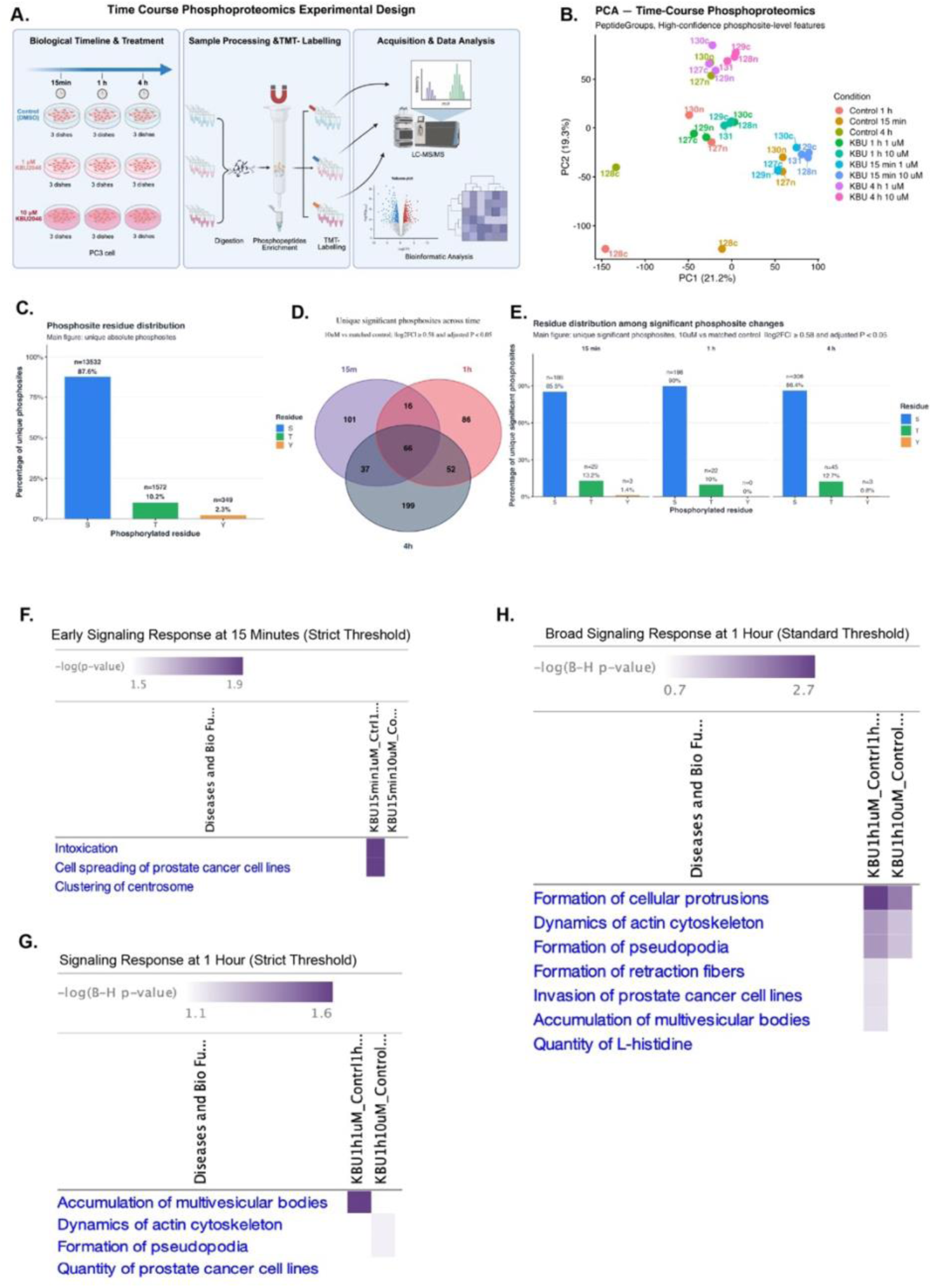
Temporal– and Concentration-Dependent Phosphoproteomic Profiling of KBU2046 treatment. (**A**) **Time-Course Phosphoproteomics Workflow**. Schematic of the experimental design. TMT labeling used a 9-plex labeling-per-time-point strategy, permitting combined LC-MS/MS analysis. Data are derived from n = 3 independent biological replicates per treatment condition. (**B-C**) **Global variance and residue distribution.** (**B**) Principal component analysis of the quantified phosphorylation sites. (**C**) Bar chart displaying the distribution of the unique phosphorylation sites for serine (S), threonine (T) and tyrosine (Y). ( **D-E**) **Quantitative Temporal Phosphoproteomics Deconvolutes the Mechanism of Action. Temporal Overlap and Active Signaling Integrity.** (**D**) Venn diagram analysis of unique significant phosphosites (|Log2 FC| ≥ 0.58, adjusted p-value < 0.05) in the 10 µM condition. (**E**) Residue distribution of the unique significant phosphosites across time intervals. (**F-H**) **Ingenuity Pathway Match Analysis of Diseases and Bio Functions**. Heatmaps display functional enrichment scores based on Benjamini-Hochberg (B-H) p-values, with darker purple indicating greater statistical significance. Findings from strict threshold selection criteria (|Log2 FC| ≥ 1.0) for 15-minute (**F**) and 1-hour (**G**) time points and standard threshold criteria (|Log2 FC| ≥ 0.5) for 1-hour time points (**H**) are shown.

Residue distribution analysis of the unique phosphosites detected that 87.6% were phosphoserine, 10.2% phosphothreonine, and 2.3% phosphotyrosine (**Figure 4C**). This distribution aligns with the canonical frequencies of eukaryotic phosphorylation sites, supporting the technical fidelity of the enrichment procedure [50, 51]. To ensure that these proportions were not artifacts of the analytical methods, a supplementary sensitivity analysis compared unique absolute phosphosites with expanded site occurrences, showing consistent results between the different counting approaches (**Supplementary Figure S1**). An intersection analysis of the significantly altered phosphosites detailed shifting signaling states across sequential time intervals. Within the 10 µM cohort, 16 unique phosphosites were shared exclusively between the 15-minute and 1-hour intervals, while a core set of 66 phosphosites remained conserved across all evaluated time points (**Figure 4D**). An evaluation of the residue distribution specifically among these actively responding protein subsets demonstrated that the hierarchical distribution of serine, threonine, and tyrosine is largely maintained across all three time points, despite minor temporal fluctuations (**Figure 4E**). Collectively, these data suggest that KBU2046 exerts dynamic temporal alterations of the global signaling networks.

To comprehensively capture the complexity of phosphoproteomic modulation across all time points and concentrations, a pairwise analysis was conducted, quantifying the drug effect, time evolution, and dose comparisons using both standard and stringent thresholds (**Tables 1 and 2**). Evaluation of KBU2046’s effect compared to vehicle controls indicates that it sustains continuous, yet dynamically changing signaling activity across all observed intervals at both threshold levels. In the analysis of time progression, a direct comparison between the 4-hour and 15-minute intervals demonstrates over 1,200 distinct changes at the standard threshold, along with 911 significant alterations under more rigorous filtering. One measure of the dynamic effect is seen in **Table 1** by considering that at 10 μM KBU2046 (i.e., the therapeutic dose), the number of phosphosites with altered abundance progressively increases from 281, 352 to 402 at the 15 min, 1-hour, and 4-hour time points, respectively. Of interest, phosphosites with increased abundance outnumber those with decreased abundance throughout, constituting 67%, 53% and 54%, respectively, demonstrating a tendency toward stabilization. However, at the 1 μM subtherapeutic dose, different patterns emerge. The number of phosphosites with altered abundance exhibits a trend opposite that observed with 10 μM, going from 508, 412 to 363 at the 15-minute, 1-hour and 4-hour time points, respectively. Further, while phosphosites with increased abundance predominate, the pattern is biphasic, going from 78%, 47% to 79%, respectively. Together, these findings indicate one pattern of continuous changes at the 10 μM therapeutic dose, but a different biphasic pattern at the 1 μM subtherapeutic dose. The latter raises the possibility of compensatory responses, which are supported by findings below.

**Table 1.** Phosphosites Altered by KBU2046: Standard Threshold Selection*.

| Comparison | Increased | % Increased | Decreased | % Decreased | Total |
| --- | --- | --- | --- | --- | --- |
| **Treatment Vs. Control** |  |  |  |  |  |
| 15 min (1 μM) vs Control | 394 | 78 | 114 | 22 | 508 |
| 15 min (10 μM) vs Control | 187 | 67 | 94 | 33 | 281 |
| 1 h (1 μM) vs Control | 192 | 47 | 220 | 53 | 412 |
| 1 h (10 μM) vs Control | 188 | 53 | 164 | 47 | 352 |
| 4 h (1 μM) vs Control | 286 | 79 | 77 | 21 | 363 |
| 4 h (10 μM) vs Control | 218 | 54 | 184 | 46 | 402 |
| **Time Evolution** |  |  |  |  |  |
| 4 h (1 μM) vs 15 min (1 μM) | 809 | 61 | 511 | 39 | 1320 |
| 4 h (10 μM) vs 15 min (10 μM) | 798 | 63 | 471 | 37 | 1269 |
| 4 h (1 μM) vs 1 h (1 μM) | 569 | 59 | 392 | 41 | 961 |
| 4 h (10 μM) vs 1 h (10 μM) | 619 | 55 | 507 | 45 | 1126 |
| **Dose Comparison** |  |  |  |  |  |
| 15 min: 10 μM vs 1 μM | 81 | 41 | 119 | 60 | 200 |
| 1 h: 10 μM vs 1 μM | 113 | 42 | 158 | 58 | 271 |
| 4 h: 10 μM vs 1 μM | 136 | 29 | 332 | 71 | 468 |
| *Pairwise comparisons using a threshold ($\vert \text{Log}_2 \text{FC}\vert \geq 0.26$; 1.2-FC) to capture a broad range of changes in unique phosphosites with altered abundance (adjusted p -value < 0.05) across three key analytical dimensions: KBU2046 vs. Control, Time Evolution, and Dose Comparison. The statistical analyses and abundance changes are derived from n = 3 independent biological replicates per condition. | | | | | |

**Table 2.** Phosphosites Altered by KBU2046: Stringent Threshold Selection*.

| Comparison | Increased | % Increased | Decreased | % Decreased | Total |
| --- | --- | --- | --- | --- | --- |
| **Treatment Vs. Control** |  |  |  |  |  |
| 15 min (1 μM) vs Control | 172 | 81 | 40 | 19 | 212 |
| 15 min (10 μM) vs Control | 144 | 67 | 71 | 33 | 215 |
| 1 h (1 μM) vs Control | 85 | 59 | 58 | 41 | 143 |
| 1 h (10 μM) vs Control | 114 | 55 | 95 | 45 | 209 |
| 4 h (1 μM) vs Control | 146 | 82 | 33 | 18 | 179 |
| 4 h (10 μM) vs Control | 186 | 53 | 164 | 47 | 350 |
| **Time Evolution** |  |  |  |  |  |
| 4 h (1 μM) vs 15 min (1 μM) | 603 | 60 | 395 | 40 | 998 |
| 4 h (10 μM) vs 15 min (10 μM) | 584 | 64 | 327 | 36 | 911 |
| 4 h (1 μM) vs 1 h (1 μM) | 368 | 58 | 268 | 42 | 636 |
| 4 h (10 μM) vs 1 h (10 μM) | 338 | 53 | 298 | 47 | 636 |
| **Dose Comparison** |  |  |  |  |  |
| 15 min: 10 μM vs 1 μM | 28 | 37 | 47 | 63 | 75 |
| 1 h: 10 μM vs 1 μM | 19 | 40 | 28 | 60 | 47 |
| 4 h: 10 μM vs 1 μM | 32 | 17 | 160 | 83 | 192 |
| *Pairwise comparisons using a threshold ($\vert \text{Log}_2 \text{FC}\vert \geq 0.58$; 1.5-FC) to capture a broad range of changes in unique phosphosites with altered abundance (adjusted p-value < 0.05) across three key analytical dimensions: KBU2046 vs. Control, Time Evolution, and Dose Comparison. The statistical analyses and abundance changes are derived from n = 3 independent biological replicates per condition. | | | | | |

To comprehensively map the signaling landscape, a tiered analysis strategy further evaluated the data. Match analysis using QIAGEN Ingenuity Pathway Analysis (IPA) software (QIAGEN Digital Insights) [52] was used to conduct a pathway-centric analysis. A standard threshold of (|Log2 FC| ≥ 0.5, 1.4-FC) was applied to provide the sensitivity to detect broad network shifts, while a stricter target-centric threshold (|Log2 FC| ≥ 1.0, 2.0-FC) was used to detect the primary signaling events happening early in the time course. Phosphosite-level abundance tables collapsed to one row per gene were analyzed using Core Analysis and Comparison Analysis to compare signaling divergence across dosage groups or time points, as described in **Supplementary Materials and Methods**.

The 15-minute phosphoproteome evaluation outlines the early phase of the cellular response to treatment. The data reveal functional differences that vary with dosage. Specifically, when the 1 μM subtherapeutic concentration is compared to control at the Log2 FC ≥ 1.0 threshold using the IPA Disease and Bio Functions Match analysis [52], there is significant enrichment of cell spreading of prostate cancer cell lines (Benjamini-Hochberg p-value: 4.17e-02) and of intoxication processes (Benjamini-Hochberg p-value: 4.17e-02) (**Figure 4F**). Under Gene Ontology taxonomy, intoxication processes denote changes in cell structure and/or those indicative of cell stress. In contrast, when this same analysis is applied to the 10 μM therapeutic concentration compared to the control, no such enrichment is identified. As discussed above, this provides another example of differential response patterns as a function of concentration, with additional ones provided below.

When the above analysis is performed at 1-hour interval, it demonstrates further evolution in signaling networks that are relevant to the regulation of cellular structure. At the 1 µM subtherapeutic level, with the strict threshold (|Log2 FC| ≥ 1.0), IPA Disease and Bio Functions Match analysis [52] revealed significant enrichment only for one function, the accumulation of multivesicular vesicles (Benjamini-Hochberg p-value: 2.72e-02) (**Figure 4G**). Here too, this same analysis applied to the 10 μM therapeutic concentration did not reveal any significant changes. It is important to recognize that this analysis of the 1-hour time point applied strict threshold criteria. The strict threshold criteria-based analytics were included for completeness in the 1-hour time point, as it was undertaken in the preceding 15-minute time point analysis. However, the strict threshold criteria were designed to rigorously identify early signaling events and thus to be most informative in the 15-minute analysis. In order to capture broader network alterations at the 1-hour time point, the standard threshold (|Log2 FC| ≥ 0.5) was applied. Under this criterion, the 1 and 10 μM concentrations revealed concordant significant processes that related to the formation of cellular protrusions (Benjamini-Hochberg p-values for 1 and 10 μM: 2.02e-03 and 8.58e-03), dynamics of actin cytoskeleton (Benjamini-Hochberg p-values for 1 and 10 μM: 1.67e-02 and 4.58e-02), and formation of pseudopodia (Benjamini-Hochberg p-values for 1 and 10 μM: 1.67e-02 and 4.58e-02) (**Figure 4H**). With respect to the 4-hour interval, no significant alterations were identified using the IPA Disease and Bio Functions Match analysis. This was the case for 1 and 10 μM concentrations, each performed under standard and strict selection criteria. Taken together, the above analyses identify significant effects on processes related to cell movement and do so in a selective manner. Given their identification only at subtherapeutic doses at 15-minute intervals, but with both doses at 1-hour intervals, which are then lost at 4-hour intervals, these findings support a dynamic signaling process. This notion is supported by findings presented above in **Table 1** and by findings below.

Following extensive structural alterations observed early after treatment, the 4-hour interval was examined to assess the cell’s response to the induced physical changes. To investigate the signaling cascades mediating the cellular response at the 4-hour time point, IPA Upstream Analysis was conducted across all evaluated time points. The 15-minute and 1-hour intervals did not yield statistically significant upstream regulators that met rigorous threshold criteria. However, the 4-hour interval revealed a distinct regulatory network, identifying significant alterations of MET signaling across both treatment concentrations. Specifically, the downstream targets of MET [53, 54] were significantly modulated in both the 1 µM subtherapeutic condition (Benjamini-Hochberg p-value: 9.78e-03) and the 10 µM therapeutic condition (Benjamini-Hochberg p-value: 2.17e-02) (**Figure 5A**). This is of high interest based on findings to date because MET signaling requires the receptor to be trafficked through endosomal compartments. This process is heavily dependent on active cytoskeletal dynamics and continuous vesicle recycling [55–58]. As shown above, the kinetic arrest of focal adhesions and the concurrent suppression of the vesicular transport machinery following KBU2046 treatment indicate that both of these processes are disrupted. Together, these findings support that disruption in the intracellular routing required for proper MET activation and the propagation of its signals is occurring.

**Figure 5.**
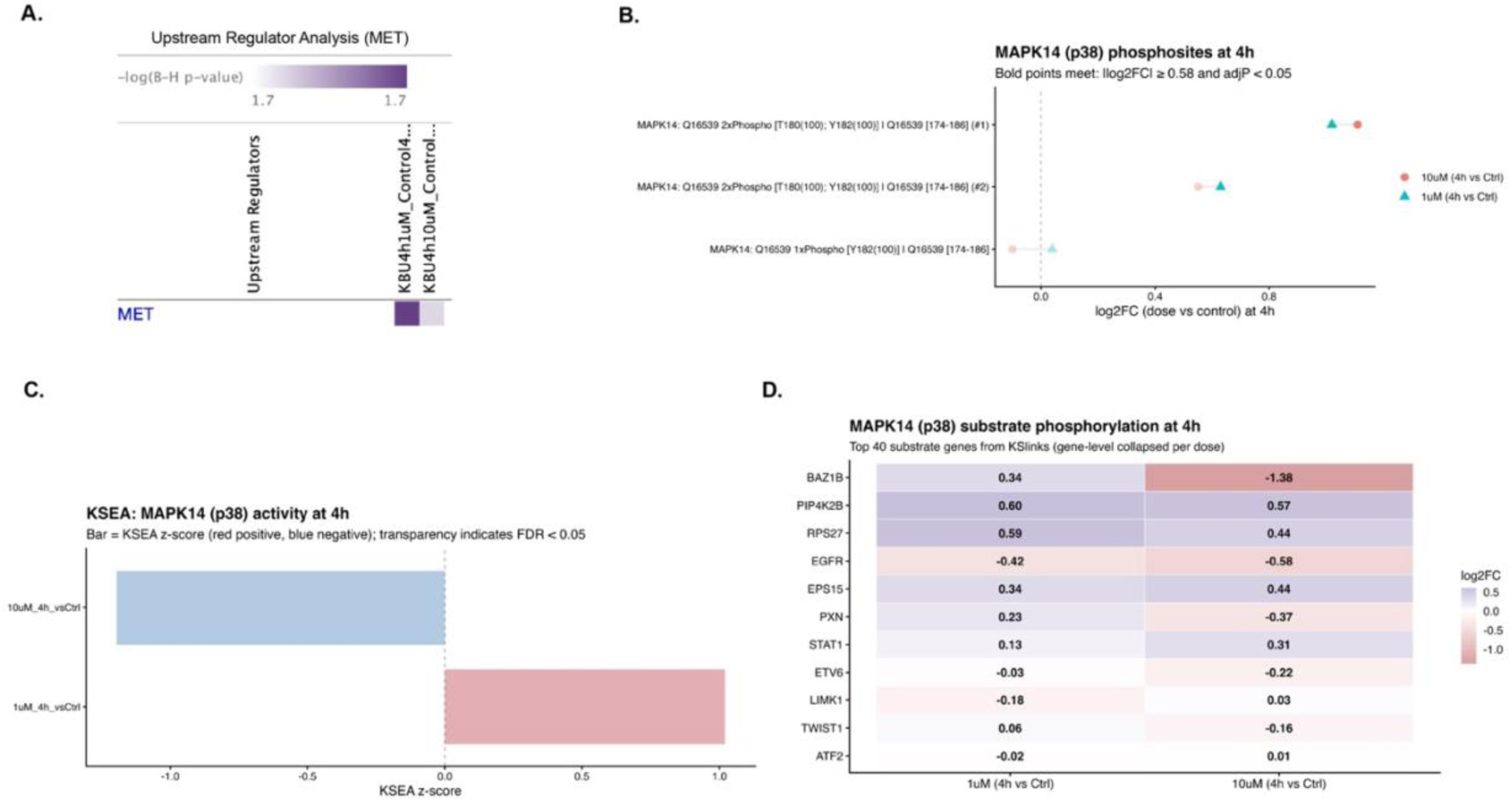
Kinase and Pathway Analysis of Later Stage Phosphoproteomic Alterations. (**A**) **Upstream Regulator Analysis (MET).** Upstream analysis using threshold (|Log2 FC| ≥1.0). (**B**) **MAPK14 Activating Phosphorylation**. Dumbbell plot showing MAPK14/p38 T180 and/or Y182 phosphosite-level changes at the 4-hour time point. Bold points indicate phosphosite entries meeting |log2 FC| ≥ 0.58 and adjusted p-value < 0.05. (**C**) **MAPK14 Kinase Substrate Enrichment Analysis Activity**. KSEA bar plot showing MAPK14/p38 substrate-network z-scores at 4 hours for 1 µM and 10 µM KBU2046 compared with control; transparency indicates whether FDR < 0.05. (**D**) **MAPK14 Substrates.** Heatmap showing phosphoproteomic Log2 FC changes in selected MAPK14-linked substrate genes at 4 hours.

This disruption of intracellular routing and associated integrin stasis results in a state of physical restriction [59]. The continuous cycling of surface integrins via clathrin-mediated endocytosis is necessary for focal adhesion disassembly [31]. Interrupting this intracellular routing may lead to prolonged structural anchoring to the extracellular matrix. We hypothesized that this combination of kinetic arrest and defective trafficking induced by KBU2046 creates a state of structural stress, which in turn activates specific mechanosensitive survival networks. To investigate how the cell manages this physical tension, the activation state of known stress integrators was evaluated.

Throughout the literature by us and others, MAPK14 is established as an integrator of response to these types of cellular events [60–64]. Prior studies demonstrate that mechanical stress signals are transduced through a specific signaling axis where Rho-family GTPases (such as Rac1 and Cdc42) activate MAPK14 (p38α), distinguishing this mechanosensitive pathway from alternative MAPK cascades activated by growth factors or cytokines [65]. In light of these findings, we interrogated our dataset to examine the specific activation status of MAPK14 and evaluate its role in managing the observed physical stress.

Functional activation of MAPK14 is structurally governed by a highly conserved threonine glycine tyrosine (Thr-Gly-Tyr) motif. In the resting state, this unphosphorylated loop sterically restricts the catalytic cleft. Dual phosphorylation of threonine (T) 180 and tyrosine (Y) 182 alters local electrostatic properties and induces a conformational rearrangement that unveils the substrate-binding pocket, resulting in a shift associated with downstream target engagement [66]. Site-specific phosphoproteomic data indicate significant dual phosphorylation of MAPK14 at these canonical T180 and Y182 residues for both 1 and 10 µM concentrations at 4 hours (**Figure 5B, Supplementary Table S3, and Supplementary Data** (**Bioinformatics Raw Data**)). The exact quantitative values and significance metrics for each mapped MAPK14 peptide entry visualized in **Figure 5B** are provided in **Supplementary Table S3**. In TMT-based quantitative proteomics, multiple observations of the same target sequence can exhibit minor quantitative variation in reporter ion intensities, often due to differing ion charge states or to ratio compression caused by background interference. While minor quantitative variation in the dose response trend exists between these spectral observations [41–43], these fluctuations fall within the expected technical boundaries of multiplexed quantification. The consistent positive magnitude of these modifications across independent spectral observations shows the activation loop remains hyperphosphorylated relative to the basal state. Together, these site-specific phosphoproteomic findings support increased activation-loop phosphorylation of MAPK14/p38 under KBU2046-associated cellular stress conditions.

Next, Kinase-Substrate Enrichment Analysis (KSEA) was performed, with details provided in the **Supplementary Materials and Methods**, to assess whether the phosphorylation status of MAPK14 was accompanied by a coordinated downstream MAPK14 substrate phosphorylation signature [67] (**Figure 5C**). Despite the increased MAPK14 activation-loop phosphorylation observed with treatment, global evaluation of the MAPK14 downstream substrate network did not show statistically significant coordinated effects (1 µM condition z-score: 1.02, adjusted p-value: 0.49; 10 µM condition z-score: –1.20, adjusted p-value: 0.45). These results suggest that increased MAPK14 phosphorylation was not accompanied by broad, coordinated activation of its annotated downstream substrate network. However, analysis of individual MAPK14-linked substrate phosphorylation events provided additional insight into possible substrate-specific effects (**Figure 5D**). To further assess substrate-specific effects, annotated MAPK14-linked substrate genes were individually evaluated for phosphoproteomic changes. From this, it was found that 10 μM KBU2046 was associated with a significant decrease in a BAZ1B S705/S708-containing phosphopeptide (Log2 FC: –1.38, adjusted p-value: 6.49e-15). No other significant changes were observed at the 10 μM concentration. To determine if this represented an active dephosphorylation event or a shift in baseline protein abundance, the BAZ1B protein abundance was interrogated. This revealed that total BAZ1B abundance levels were concurrently reduced under the 10 µM compared to control (Log2 FC: –1.09, adjusted p-value: 3.57e-15) (**Supplementary Figure S4 and Supplementary Data** (**Bioinformatics Raw Data**)). Thus, the reduction in phosphorylation is largely consistent with reduced total BAZ1B abundance rather than isolated site-specific dephosphorylation. Importantly, the 1 μM subtherapeutic dose compared to control did not induce any significant effect on BAZ1B phosphorylation, nor on its total protein abundance (Log2 FC: 0.17, adjusted p-value: 0.398) (**Supplementary Figure S4 and Supplementary Data** (**Bioinformatics Raw Data**)), nor on any other annotated MAPK14-linked substrate. BAZ1B has been implicated in influencing cell fate in response to cell stress [68–71]. As the 1 μM concentration has subtherapeutic effects on cell migration while the 10 μM concentration is therapeutic, the observed alterations in BAZ1B are associated with therapeutic response. Given its biological role, current findings support the idea that in the current system, altered BAZ1B abundance and phosphorylation may be playing a role in mediating cell stress response to therapy-induced migrastasis.

## 4. Discussion

By integrating three separate proteomics datasets, involving whole-cell, membrane-localized, and temporal– and concentration-dependent phosphoproteomic analytics, we establish a new framework for the therapeutic induction of migrastasis (**Figure 6**). At the cellular level, KBU2046 inhibits the vesicular transport system, which then disrupts the cycling of plasma membrane proteins, such as integrin β1, which is a critical cell adhesion protein. Continued cycling of integrins is necessary to support the dynamic formation of focal adhesions at the leading edge of the cell and their removal at the lagging edge. Disruption of this dynamic process by KBU2046 leads to stasis of focal adhesions and the inability of cells to move. The static formation of focal adhesions induces mechanical stress, activating the MAPK14 stress response pathway. This would normally trigger pathways that would induce cell death, but the decreased abundance of BAZ1B removes this response.

**Figure 6.**
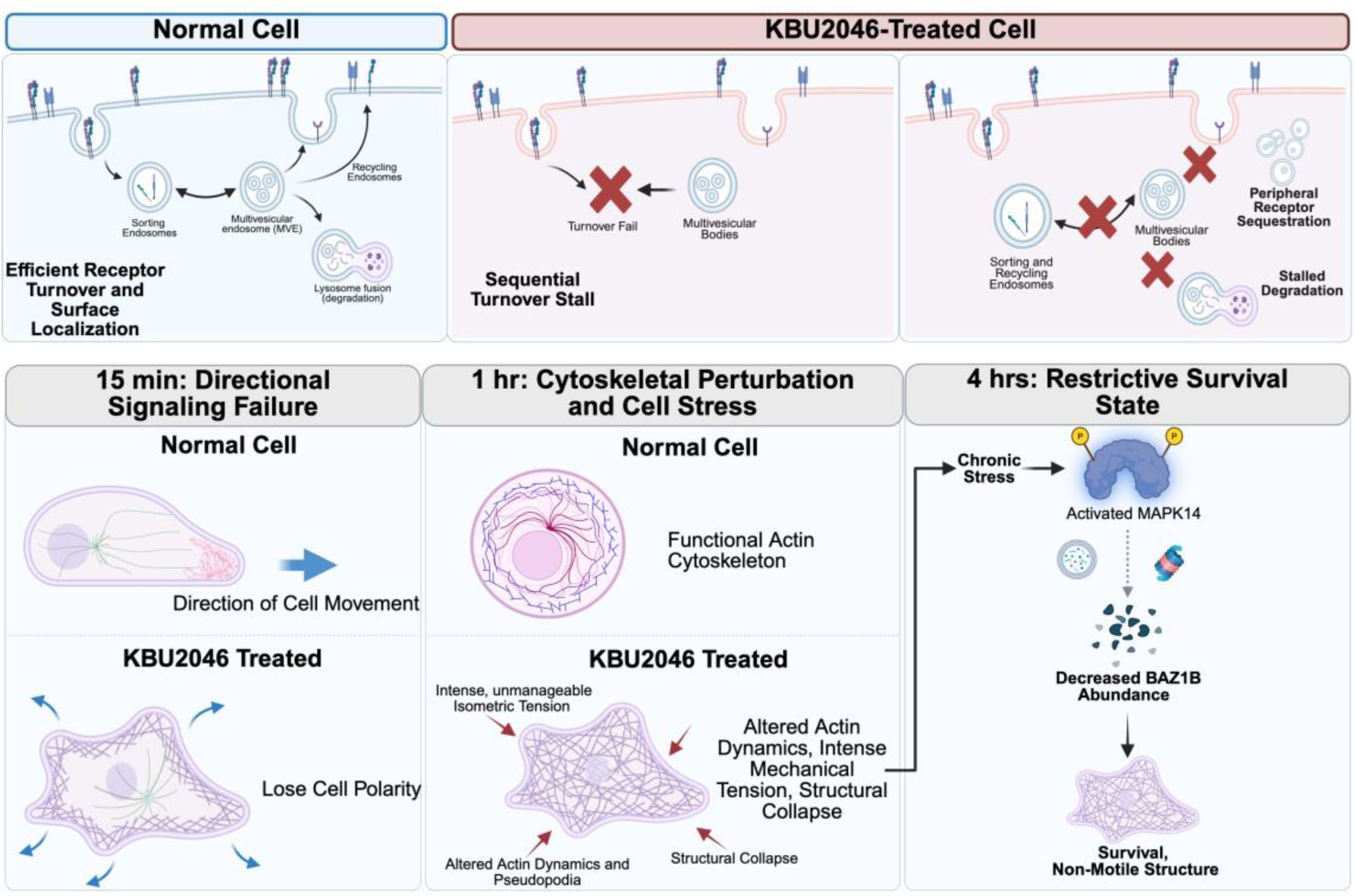
Temporal model of KBU2046 induced functional restriction. (**Top**) Attenuation of vesicular transport decreases membrane protein turnover. (**Bottom**) Proposed temporal progression outlining the cellular response across evaluated time points, detailing alterations in structural architecture, mechanical tension, and kinase network engagement.

Experimental and clinically approved drugs that act by disrupting the cytoskeleton cause widespread toxicity. Consistent with the lack of cytotoxicity associated with KBU2046, our data suggest an alternative mechanism: instead of acting as a cytotoxic agent that destroys cellular components, KBU2046 functions as a regulator of spatial and temporal processes. It modifies intracellular trafficking and the organization of the migration machinery, thereby inhibiting cell movement without causing broad proteome degradation. This finding provides a new window into understanding how cells regulate a fundamental biological process. It also provides a mechanistic basis for a new therapeutic strategy. The reality of the latter is supported by previously reported systemic therapeutic efficacy and lack of toxicity [8, 12].

One aspect of the regulatory mechanism is the physical preservation of the cellular architecture. Evaluation of the whole-cell proteome demonstrates conservation of migration components in the face of KBU2046-induced migrastasis, with structural elements remaining at levels comparable to vehicle controls. The stable abundance of these proteins suggests migrastatic arrest induced by KBU2046 is driven by functional restriction rather than a breakdown of cytoskeletal networks.

To map the nature of the functional restriction, spatial analyses and membrane proteomics were evaluated. Immunofluorescence imaging reveals a peripheral sequestration of activated integrin β1. Correspondingly, global spatial analysis of the proteins with decreased abundance indicates a targeted enrichment for transport vesicle compartments. Furthermore, membrane proteomics reveals an increased enrichment of plasma membrane organization pathways alongside a reduction in specific hypoxia– and progesterone-response signatures, which are established drivers of cellular invasion [45, 46]. These findings support that the decreased abundance of vesicular transport-related proteins deprives the cell of the ability to internalize and redistribute adhesion structures, resulting in the accumulation of intact motility hardware at the cell periphery. The resulting spatial disconnect breaks the receptor turnover cycle required for continuous forward movement. The failure in receptor recycling explains how cell movement is restricted without degrading the migration machinery. Further, this physical restriction induces mechanical stress. It has been shown in other systems that plasma membrane repair acts as a compensatory response to mechanical tension [72], and consistent with this, we observed an increased enrichment of this machinery in our current system. Given that maintaining cellular structural machinery accounts for a substantial fraction of total cellular energetic demands [73], it stands to reason that KBU2046 induction of mechanical stress would lead to increases in energy-generating machinery, and this is what we observed.

Complementing the spatial proteomic framework, temporal– and concentration-dependent phosphoproteomic data delineate dynamic and adaptive changes in cellular signaling networks. Considering phosphoproteomic data in an integrated fashion, selective effects on processes related to cell movement are identified. Given the documented phenotypic function of KBU2046, these represent anticipated findings. However, anticipated should not be taken as unimportant. The signal was clear and selective, and given the broad nature of the screens, its identification of enrichment of cell movement processes makes a powerful statement. It independently corroborates findings reported to date, demonstrating both this phenotypic function as well as the specificity of effect. Of very high interest, and not previously recognized, is the dynamic nature of these signaling events. Specifically, cell movement processes are enriched at the early 15-minute time point, but only with the subtherapeutic 1 μM dose, at the 1-hour time point with both subtherapeutic and therapeutic doses, and at the 4-hour time point, it is not observed with either dose.

Associated analytics provide a mechanistic explanation for temporal– and concentration-dependent changes in enrichment profiles that can be generally characterized as a broad cellular compensatory response to structure-induced stress. This is supported by considering the broad patterns of phosphoproteomics profiles. Both 1 and 10 μM doses favor phosphosites with increased abundance over those with decreased abundance, and this is seen over all time points evaluated. However, for 10 μM, the number of phosphosites with altered abundances (i.e., exhibiting increased or decreased phosphorylation) progressively increases with time, while for 1 μM, they progressively decrease. Further, for 10 μM, the percentage of phosphosites with increased abundance steadily decreases with time, while for 1 μM, there is a marked biphasic high-low-high pattern seen with the progression of time.

A consideration of specific signaling nodes provides additional support for cellular compensatory responses that are temporal– and concentration-dependent. Identification of enrichment in MET signaling was only seen at the 4-hour time point. Vesicular transport is necessary to maintain MET signaling, and as we show that KBU2046 disrupts vesicular transport, this provides a mechanistic rationale. Of high interest, it has been shown that the co-internalization of integrin β1 with c-Met and its subsequent trafficking to autophagy-related endosomal compartments is required to sustain functional MET signaling, a cooperative “inside-in” pathway essential for metastatic survival [74]. Knowing that MAPK14 is downstream of MET and has been shown to be involved in the cellular response to structural stress, it was then shown to be phosphorylated at specific sites known to activate it [66, 75], and this was seen with both 1 and 10 µM doses. An important response difference was then identified by examining substrates downstream of MAPK14, demonstrating that only the 10 µM dose decreased its phosphorylation. BAZ1B influences cell fate in response to cell stress; its continuous presence promotes apoptosis, whereas its deactivation is required to initiate survival and repair pathways [68–71]. Our demonstration that the aggregate abundance of BAZ1B was decreased is consistent with reports showing that the MAPK14 stress network acts as an indirect trigger [71], engaging protein degradation pathways [76–78] to clear targeted proteoforms in response to mechanical tension [79].

## 5. Limitations and Future Directions

The current analysis used the PC3 cell line, an aggressive model of androgen-independent, castration-resistant PCa derived from a bone metastasis. While PC3 cells offer a robust framework for evaluating metastatic migration, relying on a single in vitro system presents potential biological limitations. Future studies should aim to evaluate these findings across a broader panel of cell lines and in patient-derived organoids to account for genomic heterogeneity. Furthermore, because mass spectrometry only captures static snapshots of cellular states, resolving the kinetic progression of this intracellular transport inhibition will require live-cell microscopy tracking of fluorescently tagged integrins and MET receptors. Additionally, the analytical methodologies used in this study rely on peptide-level measurements, which limit the ability to fully characterize intact proteoforms. Proteome complexity arises from combinations of genetic variations, alternative splicing, and post-translational modifications that create distinct protein species [80, 81]. Future intact protein or top-down proteomic analysis is needed to identify the specific proteoforms mediating the response to KBU2046 and define the mechanism of migrastasis.

It will also be important for future investigations to evaluate key changes in protein abundance by an orthogonal assay. Further, dedicated investigations will be required to ascertain the importance of potential regulatory proteins. For example, for BAZ1B, it will be important to confirm changes in protein expression, e.g., by Western blot, and to then conduct a series of BAZ1B-specific engineering investigations. In this manner, it will be possible to ascertain in the current system its role in maintaining cellular viability under stress, including that related to mechanical and DNA damage, and/or potential cooperation with RUNX2, as supported by reports in the literature [82]. Related, it will be important for future studies to determine the pathways driving BAZ1B protein clearance and its impact on apoptosis.

The proteomic adaptations identified here provide insight into potential cellular survival mechanisms engaged during physical restriction. Future strategies could explore exploiting these newly defined stress networks, evaluating whether targeting these vulnerabilities may optimize a combination therapeutic approach, as supported by prior reports by us [12].

## 6. Conclusions

This study uses a quantitative dynamic approach to evaluate the biochemical mechanism of action of KBU2046. The data suggest a functional profile distinct from conventional cytotoxic interventions that rely on broad structural degradation. Rather than degrading the core cellular architecture, the compound restricts the spatial dynamics required for persistent cell migration. Immobilizing the migratory apparatus through spatial decoupling and modulation of dynamic signaling uncouples cellular migration from basal viability. Ultimately, integrating these three proteomic analyses yields a working model in which KBU2046 inhibits cellular migration by restricting intracellular transport networks, achieving targeted functional arrest without inducing generalized cytotoxicity.

## Supplementary Materials

The following supporting information can be downloaded. Supplementary Materials and Methods: Detailed Proteomic Sample Preparation and Bioinformatic and Statistical Analyses; Supplementary Table S1: Top 10 Compartment Enrichment Analysis of Proteins with Decreased Abundance Following KBU2046 Treatment; Supplementary Table S2: Top 10 Compartment Enrichment Analysis of Proteins with Increased Abundance Following KBU2046 Treatment; Supplementary Table S3: Quantitative profile of individual MAPK14 phosphorylation entries corresponding to Figure 5B; Supplementary Figure S1: Phosphosite Residue Composition Sensitivity and Distribution; Supplementary Figure S2: Standard Threshold Volcano Plot of Membrane Proteins; Supplementary Figure S3: High Stringency Threshold Volcano Plot of Membrane Proteins; Supplementary Figure S4: Quantitative evaluation of BAZ1B protein abundance and site-specific phosphorylation status; Supplementary Figure S5: Expanded pathway enrichment heatmaps from temporal phosphoproteomic analysis; Representative Confocal Images Raw Data Folder: Uncropped immunofluorescence microscopy files corresponding to the manuscript Figure 1; Extra Supplementary Files (Bioinformatics Raw Data) Folder: Comprehensive raw data files utilized for bioinformatics analysis. Supplementary Data S1: Curated gene lists used for compartment and adhesome annotation; Supplementary Data S2: Whole-cell proteomics source data and bioinformatics outputs, including PCA, volcano plot, motility hardware heatmap, and curated compartment analysis outputs; Supplementary Data S3: Membrane proteomics source data and bioinformatics outputs, including PCA, volcano plot, gene ontology enrichment, and membrane purity validation outputs; Supplementary Data S4: Time-course phosphoproteomics source data and bioinformatics outputs, including PCA, volcano plot, IPA analysis, phosphosite quality-control outputs, and MAPK14/p38 mechanistic analysis tables; Proteomics Experiments and LC-MS Details Folder: Proteome Discoverer export files and instrument method parameter files, including LC method and MS method.

## Author Contributions

Conceptualization, R.C.B., W.C., and H.C.-H.L.; development of methodology, W.C., S.R., C.G., H.C.-H.L., N.T.W., F.Q., J.W.Z., K.L.O. and R.C.B.; investigation, W.C., F.Q., J.W.Z., K.L.O., and R.C.B.; data analysis, W.C., S.R., C.G., H.C.-H.L., N.T.W., and R.C.B.; resources, R.C.B. and C.G.; data curation, W.C. and S.R.; writing—original draft preparation, W.C.; writing—review and editing, W.C., S.R., H.C.-H.L., F.Q., J.W.Z., K.L.O., N.T.W., C.G. and R.C.B..; visualization, W.C. and S.R.; supervision, R.C.B., C.G., H.C.-H.L., and N.T.W.; funding acquisition, R.C.B.. All authors have read and agreed to the published version of the manuscript.

## Institutional Review Board Statement

Not applicable.

## Informed Consent Statement

Not applicable.

## Data Availability Statement

The mass spectrometry (MS) proteomics data have been deposited to the ProteomeXchange Consortium via the PRIDE partner repository with the dataset identifiers PXD077603, PXD077611, and PXD077844. Reviewer access details: Log in to the PRIDE website (https://www.ebi.ac.uk/pride/login) using the following details: Project 1 accession: PXD077603, Token: fLNGY3Suf0AT. Project 2 accession: PXD077611, Token: zBxMmK2DjosP. Project 3 accession: PXD077844, Token: ryQJqR4CmjEo. Additional data will be made available upon request.

## Supporting information

Supplementary Materials (Bioinformatics Raw Data)Supplementary Materials and Methods

## Acknowledgments

We wish to express our gratitude to Amarnath (Amar) Natarajan, Grinu Mathew and Kabhilan Mohan for their valuable discussions and guidance on experimental protocols and approaches. We thank Sophie Alvarez, Mike Naldrett, and the Proteomics & Metabolomics Facility (RRID:SCR_021314), Nebraska Center for Biotechnology at the University of Nebraska-Lincoln, as well as Melinda Wojtkiewicz and Dragana Noe and the University of Nebraska Medical Center Multiomics Mass Spectrometry Core Facility (RRID: SCR_012539), for providing mass spectrometry analysis services and project support. We acknowledge use of the University of Nebraska Medical Center – UNMC Advanced Microscopy Core Facility, RRID:SCR_022467, P20 GM103427 (NIGMS, NE-INBRE), P30 GM106397 (NIGMS, NCS), P20GM130447 (NIGMS, CoNDA), P30 CA036727 (NCI, Buffett Cancer Center), S10RR02730 (NIH), S10OD030486 (NIH), Nebraska Research Initiative, UNMC Vice Chancellor for Research Office; UNMC Bioinformatics and Systems Biology Core (RRID: SCR_023767) for providing computational resources. We additionally thank James R. Talaska, Camille Hennerberg, and Heather Jensen-Smith of the UNMC AMCF for their assistance. We are grateful to Suzanne L. Topalian for the generous gift of HPV-transformed primary cell lines: 1532CPTX. Grammarly was used for superficial text editing. During the preparation of this manuscript, the authors also used BioRender to generate the graphical abstract and Figures 2A, 3A, 4A, and 6. Created in BioRender. Chen, C. (2026) https://BioRender.com/vfwsr0b, https://BioRender.com/ld50w54, https://BioRender.com/fn7vogc.

