## Supplementary Materials (Bioinformatics Raw Data)Supplementary Materials and Methods for "Integrative Proteomic Analysis Implicates Inhibition of Intracellular Protein Trafficking in Therapy-Induced Migrastasis in Prostate Cancer": MSmethod_Bergan.pdf

Orbitrap Exploris 480 Method Summary

Creator: F24NB93\Thermo Last Modified: 06/28/2022 15:35:30 by F24NB93\Thermo

Global Settings
- Use Static Source Gasses
- Use Ion Source Settings from Tune = Not Checked
- Method Duration (min)= 155
- Ion Source Type = NSI
- Spray Voltage = Static
- Spray Voltage: Positive Ion (V) = 1800
- Spray Voltage: Negative Ion (V) = 600
- Gas Mode = Static
- Infusion Mode (LC)= False
- Ion Transfer Tube Temp (°C) = 280
- APPI Lamp = Not in use
- Total Carrier Gas Flow = 4
- FAIMS Mode = Standard Resolution
- Application Mode = Peptide
- Pressure Mode = Standard
- Expected Peak Width (s) = 30
- Default Charge State = 2
- Advanced Peak Determination = False
- Xcalibur AcquireX enabled for method modifications = False

Experiment 1
- Experiment Name = MS
- Start Time (min) = 0
- End Time (min) = 155
- Cycle Time (sec) = 2

Scan MasterScan
- Orbitrap Resolution = 120000
- Scan Range (m/z) = 350-1200
- Microscans = 1
- AGC Target = Standard
- RF Lens(%) = 40
- Maximum Injection Time Mode = Auto
- FAIMS Voltages On = True
- DataType = Profile
- FAIMS CV = -45
- Polarity = Positive
- Source Fragmentation = False
- Scan Description =

Filter MIPS
- Filter Type = MIPS
- Filter Label =
- MIPS Mode = Peptide
- Relax Restrictions when too few Precursors are Found = False

Filter IntensityThreshold
- Filter Type = IntensityThreshold
- Filter Label =
- Intensity Filter Type = IntensityThreshold
- Minimum Intensity = 5000

Filter ChargeState
- Filter Type = ChargeState
- Filter Label =
- Include charge state(s) = 2-5
- Include undetermined charge states = False

Filter DynamicExclusion
- Filter Type = DynamicExclusion
- Filter Label =
- Share Dynamic exclusion list with other selected dynamic exclusion filters = False
- Dynamic Exclusion Mode = Custom
- Exclude after n times = 1
- Exclusion duration (s) = 45
- Excl. Mass Width = ppm
- Mass tolerance high = 10
- Mass tolerance low = 10
- Exclude isotopes = True
- Perform dependent scan on single charge state per precursor only = True

Filter Purity
- Filter Type = Purity
- Filter Label =
- Purity Threshold (%) = 70
- Purity Window = 0.7

Data Dependent Properties
- Data Dependent Mode= Cycle Time

Scan Event 1

Scan ddMSnScan
- Multiplex Ions Enabled = False
- Isolation Window = Custom
- Isolation Window (m/z) = 0.7
- Isolation Offset = Off
- Maximum number of multiplexed ions = 0
- Reported Mass = Offset Mass
- Collision Energy Mode = Fixed
- Collision Energy Type = Normalized
- HCD Collision Energy (%) = 36
- Orbitrap Resolution = 45000
- First Mass (m/z) = 110
- Microscans = 1
- AGC Target = Standard
- Maximum Injection Time Mode = Auto
- DataType = Centroid
- Source Fragmentation = False
- Scan Description =
