## Supplementary Materials (Bioinformatics Raw Data)Supplementary Materials and Methods for "Integrative Proteomic Analysis Implicates Inhibition of Intracellular Protein Trafficking in Therapy-Induced Migrastasis in Prostate Cancer": LCmethod_RG.pdf

---- Overview ----
Name: New Instrument Method
Comment:
Run time: 155.000 [min]
Instrument: _U3000_Nano2_Lumos on vcrdrc1050lumosms
Description:
---- Script ----
initial Instrument Setup
Sampler.LowDispersionMode: Off
Sampler.WashSpeed: 4.000 [µl/s]
Sampler.WashVolume: 50.000 [µl]
Sampler.PunctureDepth: 8.000 [mm]
Sampler.SampleHeight: 1.500 [mm]
Sampler.WasteSpeed: 4.000 [µl/s]
Sampler.DispenseDelay: 2.000 [s]
Sampler.DispSpeed: 2.000 [µl/s]
Sampler.DrawSpeed: 0.200 [µl/s]
Sampler.DrawDelay: 5.000 [s]
Sampler.RinseBetweenReinjections: Yes
Sampler.FlushVolume: 5.000 [µl]
Sampler.TransVialPunctureDepth: 11.000 [mm]
Sampler.TransLiquidHeight: 5.000 [mm]
Sampler.TransportVialCapacity: Unlimited
Sampler.LastTransportVial: R1
Sampler.FirstTransportVial: R1
Sampler.InjectMode: ulPickUp
Sampler.LoopWashFactor: 2.000
Sampler.PumpDevice: "LoadingPump"
Sampler.TempCtrl: On
Sampler.Temperature.Nominal: 5.0 [°C]
Sampler.ReadyTempDelta: None
Sampler.Temperature.LowerLimit: 4.0 [°C]
Sampler.Temperature.UpperLimit: 45.0 [°C]
PumpModule.LoadingPump.%A.Equate: "%A"
PumpModule.LoadingPump.%B.Equate: "%B"
PumpModule.LoadingPump.%C.Equate: "%C"
PumpModule.LoadingPump.Pressure.LowerLimit: 0 [bar]
PumpModule.LoadingPump.Pressure.UpperLimit: 500 [bar]
PumpModule.LoadingPump.MaximumFlowRampUp: 50 [µl/min²]
PumpModule.LoadingPump.MaximumFlowRampDown: 50 [µl/min²]
PumpModule.NC_Pump.%A.Equate: "%A"
PumpModule.NC_Pump.%B.Equate: "%B"
PumpModule.NC_Pump.Pressure.LowerLimit: 0 [bar]
PumpModule.NC_Pump.Pressure.UpperLimit: 800 [bar]
PumpModule.NC_Pump.MaximumFlowRampUp: 0.300 [µl/min²]
PumpModule.NC_Pump.MaximumFlowRampDown: 0.300 [µl/min²]
ColumnOven.TempCtrl: On
ColumnOven.Temperature.Nominal: 36.0 [°C]
ColumnOven.Temperature.LowerLimit: 25.0 [°C]
ColumnOven.Temperature.UpperLimit: 75.0 [°C]
ColumnOven.EquilibrationTime: 0.5 [min]
ColumnOven.ReadyTempDelta: None
ColumnOven.ValveRight: 1_2
0.000 [min] Equilibration
PumpModule.LoadingPump.Flow.Nominal: 4.000 [µl/min]
PumpModule.LoadingPump.%B.Value: 0.0 [%]
PumpModule.LoadingPump.%C.Value: 0.0 [%]

PumpModule.LoadingPump.Curve: 5
PumpModule.NC_Pump.Flow.Nominal: 0.300 [µl/min]
PumpModule.NC_Pump.%B.Value: 9.0 [%]
PumpModule.NC_Pump.Curve: 5
0.000 [min] Inject Preparation
Wait Sampler.Ready And PumpModule.LoadingPump.Ready And PumpModule.NC_Pump.Ready And ColumnOven.Ready
0.000 [min] Inject
Sampler.Inject
0.000 [min] Start Run
PumpModule.LoadingPump.LoadingPump_Pressure.AcqOn
PumpModule.NC_Pump.NC_Pump_Pressure.AcqOn
0.000 [min] Run
1.000 [min]
PumpModule.LoadingPump.Flow.Nominal: 4.000 [µl/min]
PumpModule.LoadingPump.%B.Value: 0.0 [%]
PumpModule.LoadingPump.%C.Value: 0.0 [%]
PumpModule.LoadingPump.Curve: 5
10.000 [min]
PumpModule.LoadingPump.Flow.Nominal: 4.000 [µl/min]
PumpModule.LoadingPump.%B.Value: 0.0 [%]
PumpModule.LoadingPump.%C.Value: 0.0 [%]
PumpModule.LoadingPump.Curve: 5
PumpModule.NC_Pump.Flow.Nominal: 0.300 [µl/min]
PumpModule.NC_Pump.%B.Value: 9.0 [%]
PumpModule.NC_Pump.Curve: 5
ColumnOven.ValveRight: 6_1
100.000 [min]
PumpModule.NC_Pump.Flow.Nominal: 0.300 [µl/min]
PumpModule.NC_Pump.%B.Value: 25.0 [%]
PumpModule.NC_Pump.Curve: 5
130.000 [min]
PumpModule.NC_Pump.Flow.Nominal: 0.300 [µl/min]
PumpModule.NC_Pump.%B.Value: 45.0 [%]
PumpModule.NC_Pump.Curve: 5
135.000 [min]
PumpModule.NC_Pump.Flow.Nominal: 0.300 [µl/min]
PumpModule.NC_Pump.%B.Value: 99.0 [%]
PumpModule.NC_Pump.Curve: 5
140.000 [min]
PumpModule.NC_Pump.Flow.Nominal: 0.300 [µl/min]
PumpModule.NC_Pump.%B.Value: 99.0 [%]
PumpModule.NC_Pump.Curve: 5
ColumnOven.ValveRight: 1_2
145.000 [min]
PumpModule.NC_Pump.Flow.Nominal: 0.300 [µl/min]
PumpModule.NC_Pump.%B.Value: 9.0 [%]
PumpModule.NC_Pump.Curve: 5
155.000 [min]
PumpModule.NC_Pump.Flow.Nominal: 0.300 [µl/min]
PumpModule.NC_Pump.%B.Value: 9.0 [%]
PumpModule.NC_Pump.Curve: 5
155.000 [min] Stop Run
PumpModule.LoadingPump.LoadingPump_Pressure.AcqOff
PumpModule.NC_Pump.NC_Pump_Pressure.AcqOff
