## Supplementary Materials (Bioinformatics Raw Data)Supplementary Materials and Methods for "Integrative Proteomic Analysis Implicates Inhibition of Intracellular Protein Trafficking in Therapy-Induced Migrastasis in Prostate Cancer": MSmethod_RG.pdf

Orbitrap Fusion Lumos Method Summary

Creator: VCRDRC1050LUMOS\Administrator Last Modified: 10/08/2021 12:02:50 by VCRDRC1050LUMOS\Administrator

Global Settings
- Use Static Source Gasses
- Use Ion Source Settings from Tune = Not Checked
- Method Duration (min)= 155
- Ion Source Type = NSI
- Spray Voltage = Static
- Spray Voltage: Positive Ion (V) = 1900
- Spray Voltage: Negative Ion (V) = 600
- Gas Mode = Static
- Infusion Mode (LC)= False
- Sweep Gas (Arb) = 0
- Ion Transfer Tube Temp (°C) = 275
- APPI Lamp = Not in use
- FAIMS Mode = Not Installed
- Internal Mass Calibration = EASY-IC
- Application Mode = Peptide
- Pressure Mode = Standard
- Default Charge State = 2
- Advanced Peak Determination = True

Experiment 1
- Experiment Name = Universal Method
- Start Time (min) = 0
- End Time (min) = 155
- Cycle Time (sec) = 3

Scan MasterScan
- Desired minimum points across the peak = 6
- MSn Level = 1
- Use Wide Quad Isolation = True
- Detector Type = Orbitrap
- Orbitrap Resolution = 120K
- Mass Range = Normal
- Scan Range (m/z) = 375-1500
- Maximum Injection Time (ms) = 50
- AGC Target = 400000
- Normalized AGC Target = 100%
- Microscans = 1
- Maximum Injection Time Type = Custom
- RF Lens (%) = 30
- Use ETD Internal Calibration = True
- DataType = Profile
- Polarity = Positive
- Source Fragmentation = False
- Scan Description =
- Enhanced Resolution Mode = Off

Filter MIPS
- Relax Restrictions when too few Precursors are Found = True
- MIPS Mode = Peptide

Filter ChargeState

Include charge state(s) = 2-5
Include undetermined charge states = False

Filter DynamicExclusion
Exclude after n times = 1
Exclusion duration (s) = 60
Mass Tolerance = ppm
Mass tolerance low = 10
Mass tolerance high = 10
Use Common Settings = False
Exclude isotopes = True
Perform dependent scan on single charge state per precursor only = False

Filter IntensityThreshold
Maximum Intensity = 1E+20
Minimum Intensity = 25000
Relative Intensity Threshold = 0
Intensity Filter Type = IntensityThreshold

Data Dependent Properties
Data Dependent Mode= Cycle Time

Scan Event 1

Scan ddMSnScan
Desired minimum points across the peak = 6
MSn Level = 2
Isolation Mode = Quadrupole
Enable Intelligent Product Acquisition for MS2 Isolation = False
Isolation Window = 1.6
Isolation Offset = Off
Reported Mass = Original Mass
Multi-notch Isolation = False
Scan Range Mode = Auto
Scan Priority= 1
Collision Energy Mode = Fixed
ActivationType = HCD
Collision Energy (%) = 35
Detector Type = Orbitrap
Orbitrap Resolution = 15K
Maximum Injection Time (ms) = 22
AGC Target = 50000
Inject ions for all available parallelizable time = False
Normalized AGC Target = 100%
Microscans = 1
Maximum Injection Time Type = Dynamic
Use ETD Internal Calibration = False
DataType = Centroid
Polarity = Positive
Source Fragmentation = False
Scan Description =
Time Mode = Unscheduled
Enhanced Resolution Mode = Off
