## Supplementary Materials (Bioinformatics Raw Data)Supplementary Materials and Methods for "Integrative Proteomic Analysis Implicates Inhibition of Intracellular Protein Trafficking in Therapy-Induced Migrastasis in Prostate Cancer": Supplementary Figures and Tables_updated061726.docx

| **Supplementary Table S1**. Top 10 Compartment Enrichment Analysis of Proteins with Decreased Abundance Following KBU2046 Treatment. | | | | | |
| --- | --- | --- | --- | --- | --- |
| **Index** | **Name** | **P-value** | **Adjusted p-value** | **Odds Ratio** | **Combined score** |
| 1 | TRANSPORT VESICLE MEMBRANE | 0.0001305 | 0.02313 | 17.54 | 156.89 |
| 2 | COATED VESICLE MEMBRANE | 0.0002224 | 0.02313 | 15.19 | 127.79 |
| 3 | COATED VESICLE | 0.001089 | 0.07548 | 9.84 | 67.11 |
| 4 | CLATHRIN VESICLE COAT | 0.001497 | 0.07785 | 39.28 | 255.48 |
| 5 | TRANSPORT VESICLE | 0.002328 | 0.08193 | 7.94 | 48.17 |
| 6 | CLATHRIN COAT | 0.002363 | 0.08193 | 30.77 | 186.11 |
| 7 | ER TO GOLGI TRANSPORT VESICLE MEMBRANE | 0.004324 | 0.1285 | 22.31 | 121.45 |
| 8 | VESICLE COAT | 0.005332 | 0.1386 | 19.96 | 104.45 |
| 9 | RECYCLING ENDOSOME MEMBRANE | 0.007637 | 0.1531 | 16.48 | 80.31 |
| 10 | MITOCHONDRIAL IRON-SULFUR CLUSTER ASSEMBLY COMPLEX | 0.009217 | 0.1531 | 138.60 | 649.60 |

**Supplementary Table S1**. Gene Ontology compartment analysis using the COMPARTMENTS curated database for the protein fraction with decreased abundance identified in the global whole cell proteomic analysis of PC3 cells treated with 10 µM KBU2046. The table details the top 10 enriched cellular components. Metrics provided include p-value, adjusted p-value, odds ratio, and combined score for each identified compartment. The complete dataset is available in the **Supplementary Materials** (**Bioinformatics Raw Data**).

| **Supplementary Table S2**. Top 10 Compartment Enrichment Analysis of Proteins with Increased Abundance Following KBU2046 Treatment. | | | | | |
| --- | --- | --- | --- | --- | --- |
| **Index** | **Name** | **P-value** | **Adjusted p-value** | **Odds Ratio** | **Combined score** |
| 1 | MITOCHONDRIAL MEMBRANE | 5.831e-7 | 0.0001361 | 7.25 | 104.04 |
| 2 | MITOCHONDRIAL ENVELOPE | 0.000001143 | 0.0001361 | 6.78 | 92.70 |
| 3 | ORGANELLE ENVELOPE | 0.000003735 | 0.0002963 | 5.18 | 64.73 |
| 4 | ORGANELLE INNER MEMBRANE | 0.00001593 | 0.0009477 | 7.00 | 77.28 |
| 5 | MITOCHONDRIAL INNER MEMBRANE | 0.00006663 | 0.003172 | 6.58 | 63.32 |
| 6 | MITOCHONDRION | 0.00009233 | 0.003662 | 4.26 | 39.59 |
| 7 | U2 SNRNP | 0.002825 | 0.08988 | 28.28 | 166.01 |
| 8 | MITOCHONDRIAL OUTER MEMBRANE | 0.003021 | 0.08988 | 7.20 | 41.78 |
| 9 | SITE OF DOUBLE-STRAND BREAK | 0.003604 | 0.09261 | 10.50 | 59.07 |
| 10 | ORGANELLE OUTER MEMBRANE | 0.004467 | 0.09261 | 6.43 | 34.80 |

**Supplementary Table S2**. Gene Ontology compartment analysis using the COMPARTMENTS curated database for the protein fraction with increased abundance identified in the global whole cell proteomic analysis of PC3 cells treated with 10 µM KBU2046. The table details the top 10 enriched cellular components. Metrics provided include p-value, adjusted p-value, odds ratio, and combined score for each identified compartment. The complete dataset is available in the **Supplementary Materials** (**Bioinformatics Raw Data**).

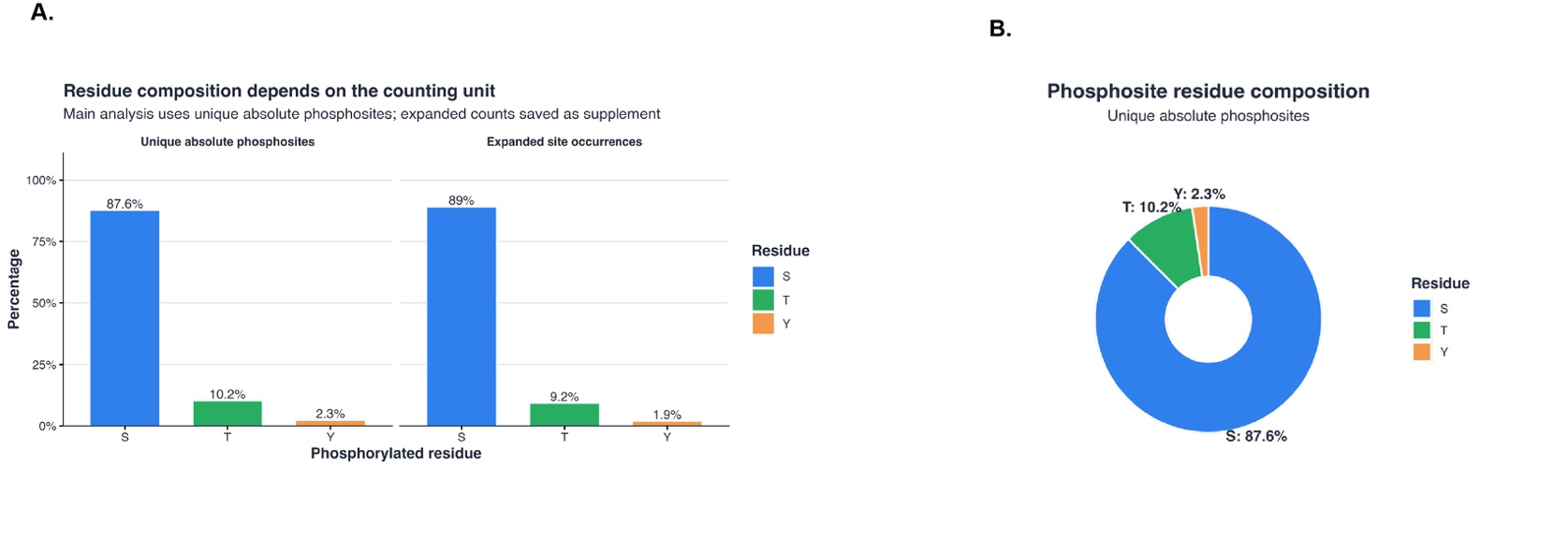

**Supplementary Figure S1. Phosphosite Residue Composition Sensitivity and Distribution**. (**A**) Bar charts comparing the residue composition of phosphorylated residues calculated via unique absolute phosphosites versus expanded site occurrences. (**B**) A pie chart visualizing the proportional residue composition of unique absolute phosphosites, demonstrating a distribution of 87.6% Serine, 10.2% Threonine, and 2.3% Tyrosine. The canonical mammalian residue distribution is robustly maintained irrespective of the bioinformatic counting methodology.

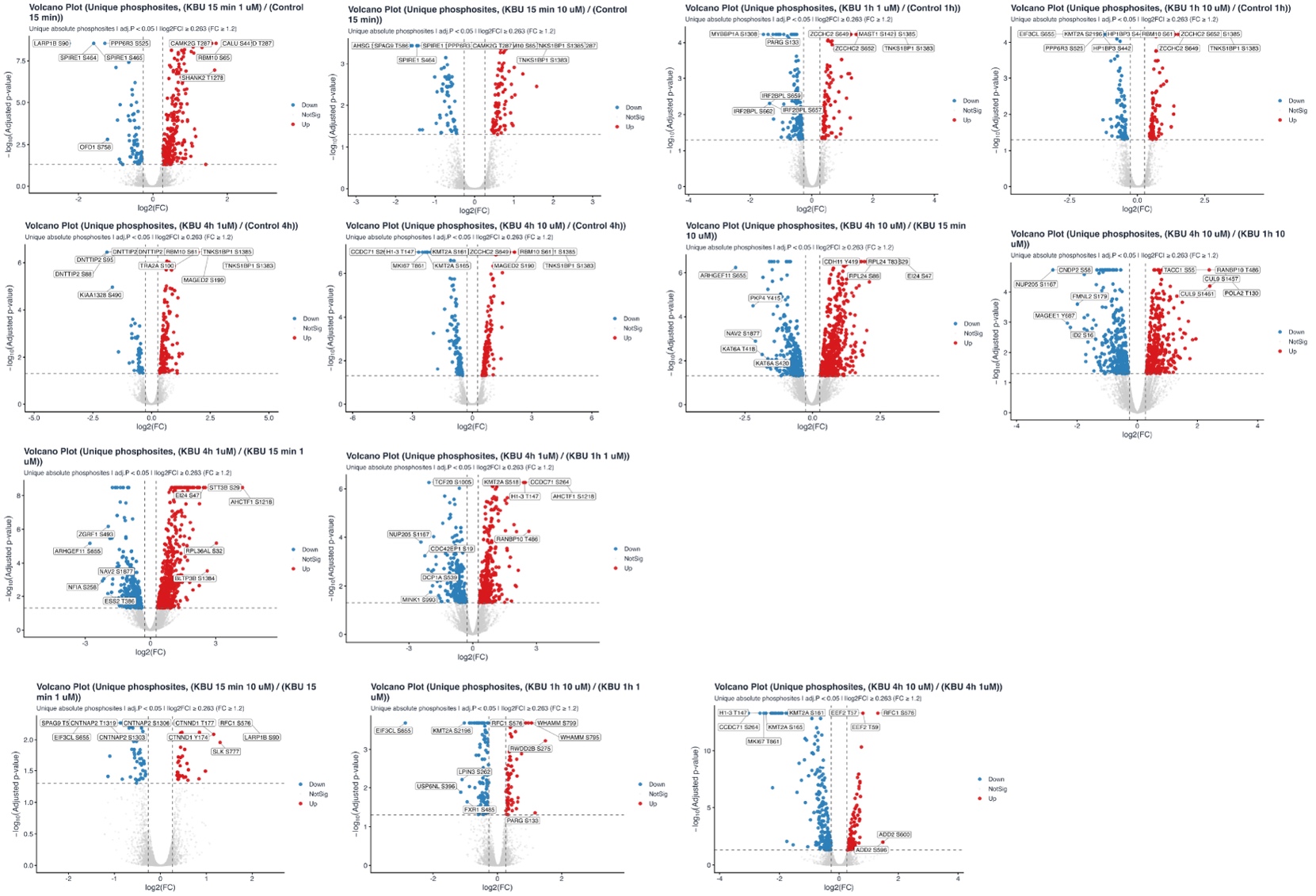

**Supplementary Figure S2. Standard Threshold Volcano Plot of Membrane Proteins.** Volcano plot highlighting the global distribution of unique absolute phosphosites with increased and decreased abundance across KBU2046 treatment conditions and time points. Significant hits are defined by the standard threshold of |Log2 FC| ≥ 0.26 and adjusted p-value < 0.05. Red points indicate phosphosites with increased abundance, blue points indicate phosphosites with decreased abundance, and grey points indicate phosphosites without significant alteration. Individual panels show dose- and time-specific comparisons, including KBU2046-treated samples versus matched controls and selected dose-to-dose comparisons.

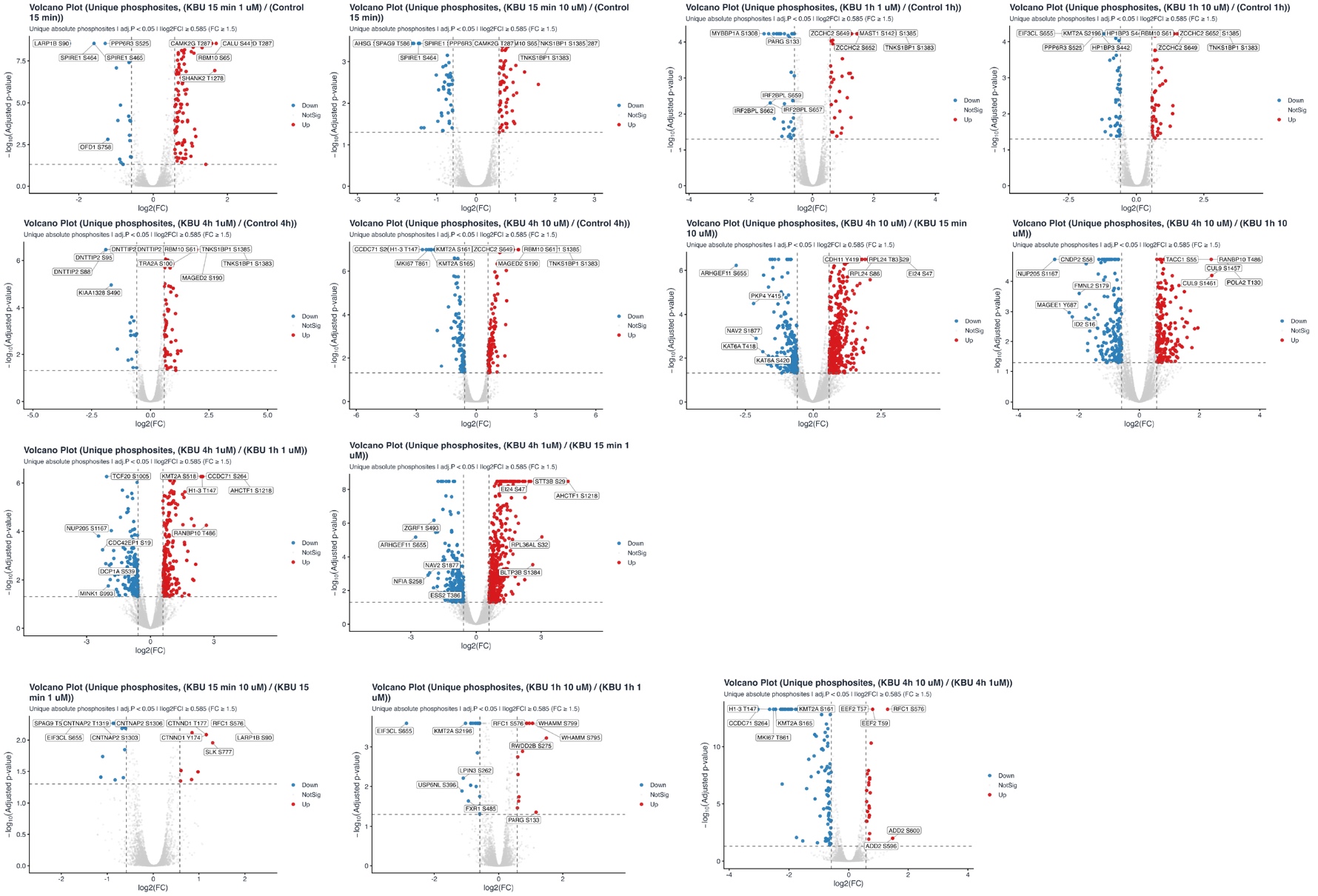

**Supplementary Figure S3. High-Stringency Threshold Volcano Plot of Membrane Proteins.** Volcano plot highlighting the global distribution of unique absolute phosphosites with increased and decreased abundance across KBU2046 treatment conditions and time points under stringent statistical filtering. Significant hits are restricted to a high-stringency threshold of |Log2 FC| ≥ 0.58 and adjusted p-value < 0.05. Red points indicate phosphosites with increased abundance, blue points indicate phosphosites with decreased abundance, and grey points indicate phosphosites without significant alteration.

| **Supplementary Table S3. Quantitative profile of individual MAPK14 phosphorylation entries corresponding to Figure 5B.** | | | | |
| --- | --- | --- | --- | --- |
| **Phosphorylation Site** | **KBU2046 Dose** | **Log2 Fold Change** | **Adjusted p-value** | **Meets Strict Threshold** |
| Site 1: 2xPhospho T180 and Y182 | 1 µM | 1.02 | 7.11E-10 | Yes |
| Site 1: 2xPhospho T180 and Y182 | 10 µM | 1.11 | 3.75E-06 | Yes |
| Site 2: 2xPhospho T180 and Y182 | 1 µM | 0.63 | 9.74E-06 | Yes |
| Site 2: 2xPhospho T180 and Y182 | 10 µM | 0.55 | 0.0489 | No |
| Site 3: 1xPhospho Y182 | 1 µM | 0.04 | 0.977 | No |
| Site 3: 1xPhospho Y182 | 10 µM | -0.10 | 0.988 | No |

**Supplementary Table S3. Quantitative profile of individual MAPK14 phosphorylation entries corresponding to Figure 5B.** This table includes individual MAPK14 peptide phosphorylation entries detected at 4 hours post-treatment with 1 µM or 10 µM KBU2046, compared with the vehicle control. The columns detail the specific phosphorylation site, KBU2046 dose, Log2 FC, adjusted p-value, and whether the entry meets the strict significance threshold (|Log2 FC| ≥ 0.58 and adjusted p-value < 0.05).

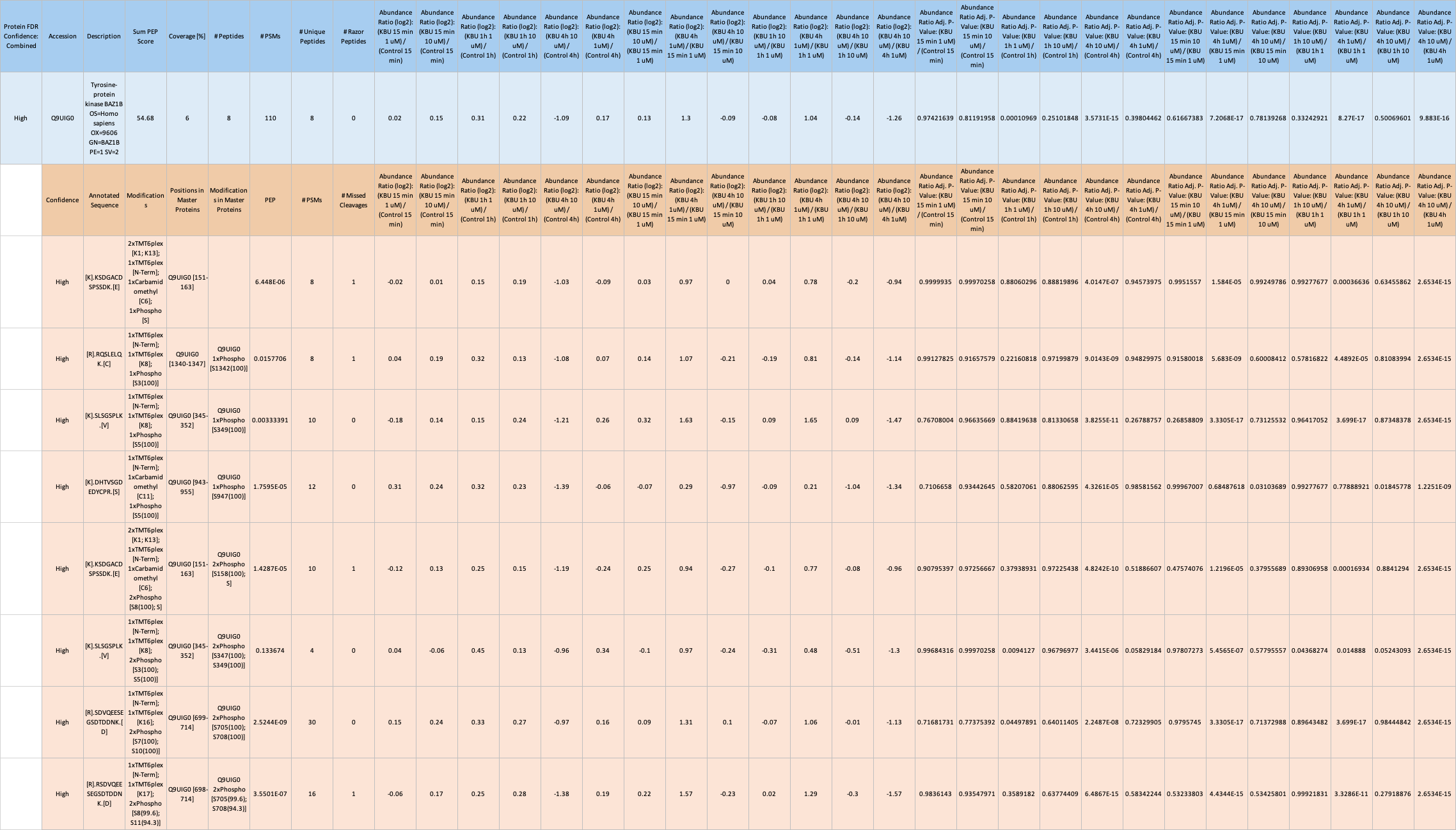

**Supplementary Figure S4**. **Quantitative evaluation of BAZ1B protein abundance and site-specific phosphorylation status.** The data summary contains mass spectrometry quantification for the BAZ1B protein and associated phosphopeptides across evaluated time intervals and treatment concentrations. Values include normalized abundance ratios, adjusted p-values, and specific phosphorylated residues. Detailed data sheets are provided in the **Supplementary Data**.

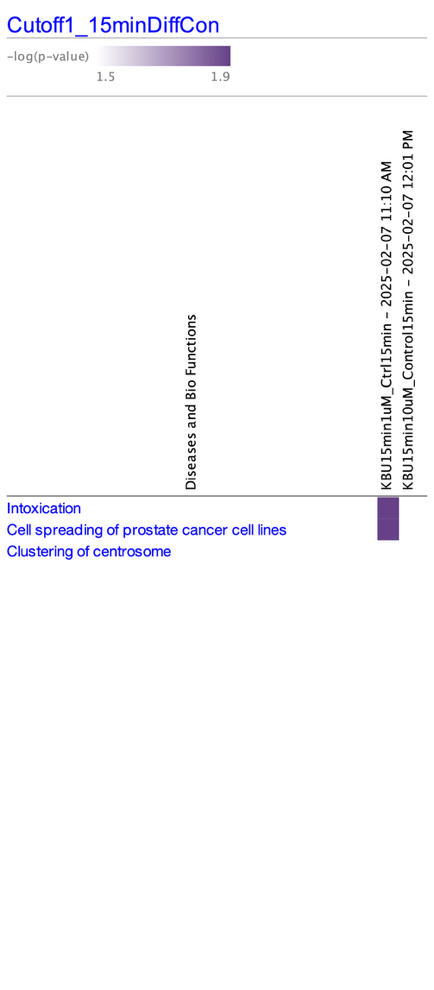

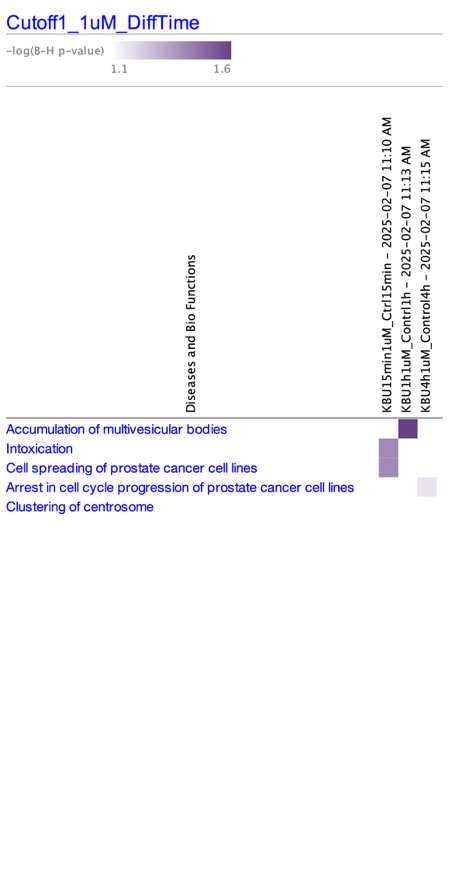

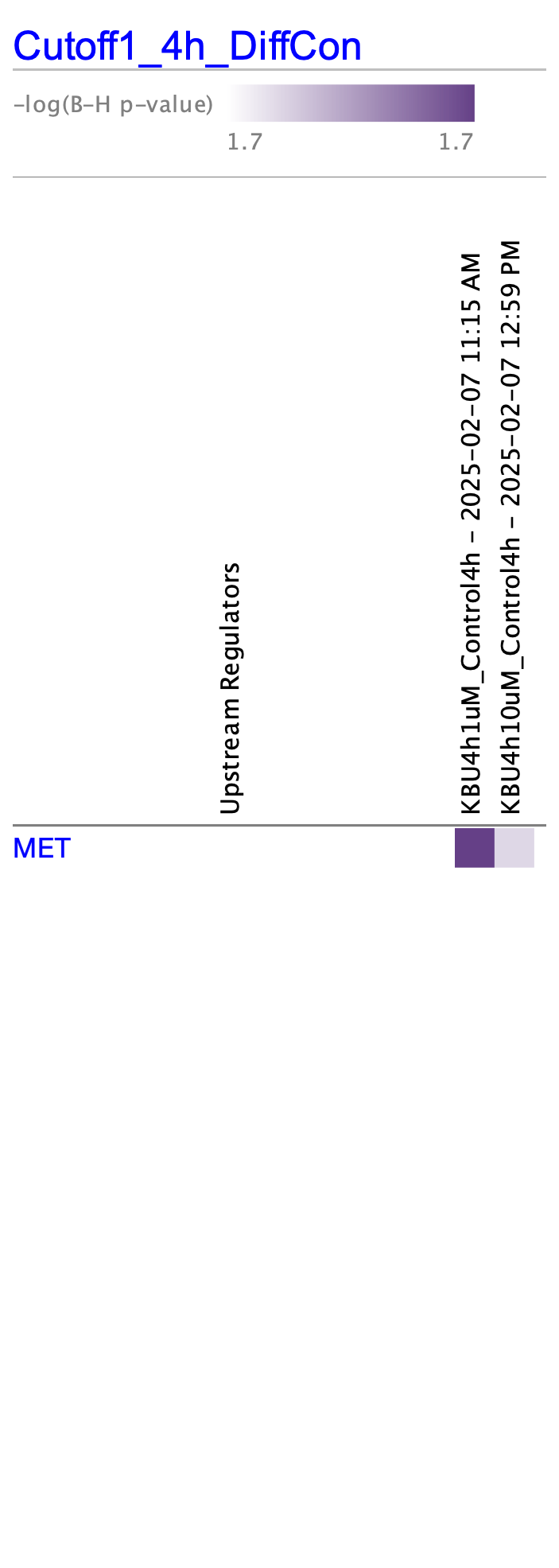

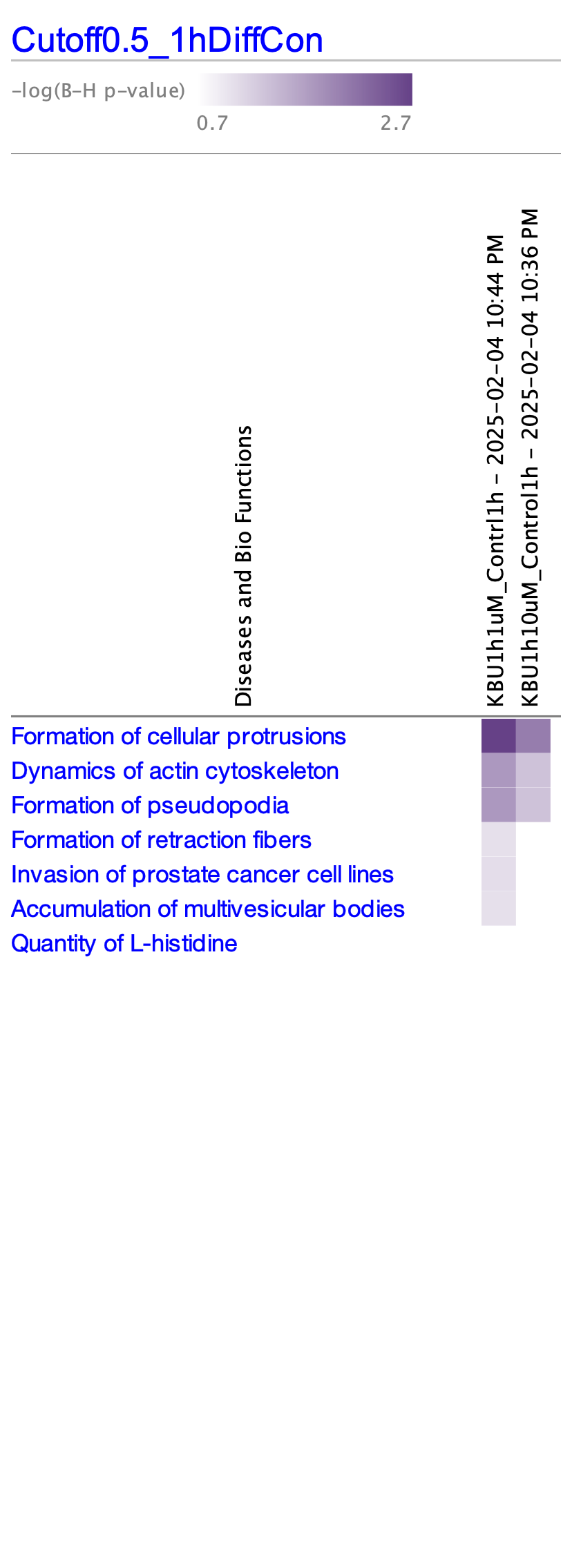

**Supplementary Figure S5.** Expanded pathway enrichment heatmaps from temporal phosphoproteomic analysis. High-resolution uncropped visualizations of the Ingenuity Pathway Analysis Disease and Bio Functions mapping across the evaluated treatment concentrations and time intervals of Figures 4F, 4G, 4H and 5A.
