## Supplementary Materials (Bioinformatics Raw Data)Supplementary Materials and Methods for "Integrative Proteomic Analysis Implicates Inhibition of Intracellular Protein Trafficking in Therapy-Induced Migrastasis in Prostate Cancer": Supplementary Materials and Methods_updated 061726.docx

**S1. Detailed Proteomic Sample Preparation and Mass Spectrometry**

**S1.1. Whole-Cell Label-Free Proteomics**

Exactly 2×10^5^ PC3 cells were seeded into 100 mm petri dishes and allowed to attach overnight. Cells were treated with 10 µM KBU2046 (n = 3) or a corresponding vehicle control (n = 3) for 72 hours. After incubation, plates were placed on ice, and cells were washed 3 times with cold PBS. Cells were scraped on ice using PBS, collected into tubes, and centrifuged at 2000 g at 4 °C for 10 minutes. The cell pellets were resuspended in RIPA buffer (89900; Thermo Scientific, Waltham, MA) supplemented with Halt Protease and Phosphatase Inhibitor Single Use Cocktail (100X) (78442; Thermo Scientific, Waltham, MA), 1 M PMSF (MilliporeSigma), and a custom 20X phosphatase inhibitor mixture containing sodium orthovanadate (J60191.AD; Thermo Scientific Chemicals), sodium fluoride (S6776 100G; MilliporeSigma), and beta glycerophosphate (G9422 10G; MilliporeSigma). Samples were kept on ice for 30 minutes, vortexing every 10 minutes to ensure complete lysis. Lysates were then centrifuged at 14000 g at 4 °C for 10 minutes to clear insoluble debris. Supernatants were collected, and protein concentrations were determined utilizing a BCA protein assay kit (A65453; Thermo Scientific, Waltham, MA) prior to downstream processing.

Sample preparation and mass spectrometry were conducted at the University of Nebraska Medical Center Mass Spectrometry and Proteomics Core Facility. Proteins were denatured and reduced at 95 °C for 2 minutes, then centrifuged at 12000 g for 10 minutes. Digestion was performed utilizing the Filter Aided Sample Preparation procedure in 10 kDa centrifugal filter tubes. Proteins were treated with 50 mM dithiothreitol (DTT) in urea (UA) buffer for 30 minutes at 37 °C, and alkylated with 50 mM iodoacetamide (IAA) in UA buffer for 30 minutes in darkness. Following washes, proteins were digested with Promega trypsin at a 1 to 100 weight to weight ratio for 18 hours at 37 °C. Peptides were purified using homemade C18 Empore tips, lyophilized, and acidified with 0.1 % formic acid.

Approximately 1 µg of peptides per sample was loaded onto a nanoflow HPLC Easy nLC 1200 system (Thermo Fisher Scientific, Waltham, MA). Liquid chromatography separation was performed at a flow rate of 300 nl/min over a 155-minute total run time. Analysis was performed on a Thermo Orbitrap Fusion Lumos mass spectrometer (Thermo Fisher Scientific, Waltham, MA) utilizing a data-dependent acquisition method with a 3-second cycle time. Full MS1 scans were acquired in the Orbitrap with a resolution of 120000, a scan range of 375 to 1500 m/z, a maximum injection time of 50 milliseconds, and an automatic gain control target of 400000. Precursor ions with charge states 2 to 5 were isolated using a 1.6 m/z window and fragmented via higher-energy collisional dissociation with a normalized collision energy of 35%. MS2 spectra were acquired in the Orbitrap at a resolution of 15000 with a maximum injection time of 22 milliseconds and an automatic gain control target of 50000. Precursor ions were excluded dynamically for 60 seconds.

Raw mass spectrometry data files were processed utilizing Proteome Discoverer software (Thermo Scientific, Waltham, MA, USA, v2.2) (Thermo Fisher Scientific, Waltham, MA) integrating the SEQUEST HT and COMET search engines. Spectra were searched against the SwissProt human database, assuming trypsin digestion. A 1% false discovery rate cutoff was applied to peptide spectral matches and protein identifications. The Log2 protein abundance ratio was calculated by dividing the average protein abundance of the treated group by the average protein abundance of the control group. Normalized protein abundance tables exported from Proteome Discoverer were used for subsequent relative protein abundance evaluation, and protein abundance tables exported from Proteome Discoverer and normalized prior to analysis were used for downstream relative protein abundance analysis. Mass spectrometry analyses were performed by the University of Nebraska Medical Center Multiomics Mass Spectrometry Core Facility (RRID: SCR 012539).

**S1.2. Membrane Enriched Subcellular Proteomics**

2x10^6^ PC3 cells were seeded into 150 mm petri dishes. Cells were treated with 10 µM KBU2046 (n = 4) and DMSO control (n = 4) for 72 hours. Membrane fractions were isolated to resolve spatial dynamics utilizing the Mem PER Plus Membrane Protein Extraction Kit (89842; Thermo Scientific, Waltham, MA) supplemented with Halt Protease and Phosphatase Inhibitor Single Use Cocktail (100X) (78442; Thermo Scientific, Waltham, MA). Extracted surface-associated proteins were chemically labeled with the Tandem Mass Tag (TMT) 10-plex Mass Tag Labeling Kit (Thermo Fisher Scientific) to enable multiplexed quantification. 100μg of each sample was reconstituted to 100μL with 100mM triethylammonium bicarbonate (TEAB) and labeled. Pierce Quantitative Colorimetric Peptide Assay (ThermoFisher Scientific, Waltham, MA) was used for peptide quantification. Liquid chromatography separation was performed at a flow rate of 300 nl/min over a 155-minute total run time. Analysis was performed on a Thermo Orbitrap Exploris 480 mass spectrometer equipped with a Field Asymmetric Ion Mobility Spectrometry Pro interface (Thermo Fisher Scientific) using a data-dependent acquisition (DDA) method with a 2-second cycle time and a compensation voltage of -45. Full MS1 scans were acquired in the Orbitrap with a resolution of 120000, a scan range of 350 to 1200 m/z, an automatic maximum injection time, and a standard automatic gain control target. Precursor ions with charge states 2 to 5 were isolated using a 0.7 m/z window and fragmented via higher-energy collisional dissociation with a normalized collision energy of 36%. MS2 spectra were acquired in the Orbitrap at a resolution of 45000 with a first mass of 110 m/z, an automatic maximum injection time, and a standard automatic gain control target. Precursor ions were excluded dynamically for 45 seconds.

Differentially abundant proteins resultant from Tandem Mass Tag membrane-enriched labeling were identified using Proteome Discoverer software (Thermo Scientific, Waltham, MA, USA, v2.2) (Thermo Fisher Scientific, Waltham, MA) integrated with SEQUEST HT and COMET search engines. Spectra were searched against the SwissProt human database, assuming trypsin digestion. Carbamidomethylation of cysteine and Tandem Mass Tag labeling modifications were set as fixed modifications, while oxidation of methionine was set as a dynamic modification. A 1% false discovery rate cutoff was applied to peptide spectral matches and protein identifications. Normalized protein abundance tables exported from Proteome Discoverer were used for subsequent relative protein abundance evaluation, and protein abundance tables exported from Proteome Discoverer and normalized prior to analysis were used for downstream relative protein abundance analysis. Mass spectrometry analyses were performed by the University of Nebraska Medical Center Multiomics Mass Spectrometry Core Facility (RRID: SCR 012539).

**S1.3. Quantitative Temporal Phosphoproteomics**

Protein Extraction and Digestion: Sample processing was performed in collaboration with the Proteomics and Metabolomics Facility (PMF) at the University of Nebraska-Lincoln. PC3 cells were seeded at a density of 1.5x10^6^ cells per 100 mm dish and incubated overnight. Cultures received treatment with complete media supplemented with either a vehicle control, 1 µM KBU2046, or 10 µM KBU2046 for 15 minutes, 1 hour, and 4 hours, utilizing three biological replicates per condition. Plates were transferred to ice. Adherent cells were washed three times with cold phosphate-buffered saline supplemented with Halt Single Use 100X protease and phosphatase inhibitor cocktail (78442, Thermo Scientific), 1 M phenylmethylsulfonyl fluoride (MilliporeSigma), and a custom 20X blend containing sodium orthovanadate (J60191.AD, Thermo Scientific Chemicals), sodium fluoride (S6776 100G, MilliporeSigma), and beta glycerophosphate (G9422 10G, MilliporeSigma). Cells were scraped on ice in a matched lysis buffer. Lysates were centrifuged at 2000 x g at 4 °C for 10 minutes. The resulting cellular pellet was collected and sent out for analysis at the Proteomics and Metabolomics Facility at the University of Nebraska-Lincoln. Cell pellets were lyzed in a solution of 2 M thiourea, 7 M urea, 10 mM DTT, 0.2 M HEPES, pH 8.0 including a 1x protease inhibitor (cOmplete EDTA-free protease inhibitor cocktail; Roche, Indianapolis, IN, USA) and 2 x phosphatase inhibitor (PhosSTOP, Roche) on ice for 10 min with the proteins subsequently reduced at 37°C for 3 hours on a thermomixer. Protein content was assayed using the CB-X protein assay from G-Biosciences (St Louis, MO, USA). One hundred and seventy micrograms of protein was taken and alkylated with 20 mM iodoacetamide (IAA) for 45 minutes at room temperature, with subsequent quenching of the IAA with equimolar DTT. The samples were diluted 3-fold with water, acetone precipitated and the pellets washed once with 70% ethanol. Each pellet was then digested with a 1:50 enzyme:substrate ratio of Lys-C in 50 µL 0.2 M EPPS, pH 8.5 at 37°C for 2 hours. Following this, a 1:50 ratio of trypsin in 50 µL water was added and digestion carried out for a further 16 hours.

To enable robust quantitative comparisons across the 27 total samples, the experimental design incorporated three multiplexed Tandem Mass Tag 10-plex batches (Thermo Fisher Scientific). An internal reference standard, created by pooling 20 µg aliquots from each of the 27 samples, was included for each 10-plex batch to facilitate precise cross-batch normalization. Each of the 150 µg peptide digests and the internal reference standard were labeled according to the following modified kit protocol. Each sample, in 100mM EPPS buffer, pH 8.5, was labeled with 450 µg of the respective Tandem Mass Tag reagent for 1 hour at RT. The labeling reaction was quenched with 5% hydroxylamine for 15 minutes and 1 µL from each was taken to check that the labeling efficiency was >99%. Labeled samples for each 10*plex set were pooled, dried under vacuum whilst frozen, and desalted utilizing C18 SepPak cartridges (Waters, Milford MA).

Phosphopeptides were enriched from each 10-plex experiment (1.5 mg of total labeled peptides) using titanium dioxide affinity chromatography. Labeled peptides were redissolved in a loading buffer of 2 M lactic acid, 60% acetonitrile, 0.1% trifluoracetic acid (TFA) and incubated with 10 mg titanium dioxide beads (GL Sciences) at 4°C on a shaker for 1.5 hours. Beads were then captured in a 200 µL STAGE tip with 2 layers of C8 membrane (3M Empore, Sigma-Aldrich) and washed sequentially to remove non-phosphorylated peptides using loading buffer followed by 80% acetonitrile, 0.1% TFA. Phosphopeptides were finally eluted using 5% ammonium hydroxide and immediately frozen, and vacuum-dried in a SpeedVac concentrator until dry. Each of the 3 sets of enriched phosphopeptides were then subfractionated into 6 fractions according to the method of Ruprecht *et al.* [1].

Analytical data acquisition was conducted at the Proteomics and Metabolomics Facility (RRID:SCR_021314) within the Nebraska Center for Biotechnology. Enriched phosphopeptides were separated using an UltiMate 3000 RSLCnano system (Thermo Fisher Scientific) coupled to a Thermo Orbitrap Eclipse Tribrid mass spectrometer. Peptides were loaded onto an Acclaim PepMap™ 100 75 um x 2 cm C18 trap column (3 um, 100A) and separated on an EASY-spray™ PepMap Neo C18 column, (75 um x 50 cm), using a 120-minute non-linear gradient during a 150min run at a flow rate of 250 nL/min. The mass spectrometer was operated in positive data-dependent acquisition mode as described previously in Abid *et al*. [2], using High-Field Asymmetric waveform Ion Mobility Spectrometry (FAIMS) to add an additional dimension of separation and multistage activation collision-induced dissociation (MSA-CID) fragmentation and generate MS/MS data. MS1 full scans were acquired in the Orbitrap at a resolution of 120,000 with a mass range of 400 to 1400 m/z. Following MS2, multiple fragment ions were synchronously isolated using synchronous precursor selection (SPS-MS3) for precise reporter ion quantitation and subjected to higher-energy collisional dissociation at a normalized collision energy of 65%. The resulting MS3 reporter ions were detected in the Orbitrap at a resolution of 50,000.

Raw mass spectrometry data files were processed utilizing Proteome Discoverer version [3.1] (Thermo Fisher Scientific). Spectra were searched against the human UniProt database (UP000005640_canonical; 20,578 entries) utilizing the Sequest HT search engine. Search parameters included full tryptic specificity, a maximum of two missed cleavages, a precursor mass tolerance of 15 ppm and fragment mass tolerance of 0.5 Da. Static modifications included cysteine carbamidomethylation and Tandem Mass Tag labeling at peptide N termini and lysine residues. Dynamic modifications for methionine oxidation and serine, threonine, or tyrosine phosphorylation followed by INFERYS rescoring were included in the search. False discovery rates for peptide and protein identification were set to <1 %. Phosphorylation site localization probabilities were calculated utilizing the ptmRS node. Quantitative analysis was performed using reporter ion intensities extracted from MS3 spectra normalized based on total phosphopeptide abundances and scaled to the internal pooled reference standard present in each batch.

**S2. Bioinformatic and Statistical Analyses**

**2.1. General Bioinformatic Workflow and Software**

Unless otherwise stated, downstream bioinformatic analyses were performed in R version 4.5.1 [3] using custom R scripts. Data import, preprocessing, and data manipulation were performed using R libraries including readxl (1.4.5), dplyr (1.1.4), tibble (3.3.0), stringr (1.6.0), readr (2.1.6), janitor (2.2.1), and tidyr (1.3.2). Data visualization was performed using ggplot2 [4] (4.0.1), ggrepel (0.9.6), and pheatmap (1.0.13). R-based enrichment analyses were performed using clusterProfiler [5] (4.16.0), enrichplot (1.28.4), org.Hs.eg.db (3.21.0), and, where applicable, msigdbr (25.1.1) for MSigDB-derived gene-set annotations. Kinase-substrate enrichment analysis was performed using KSEAapp [6], with NetworKIN filtering enabled where indicated [7].

Where applicable, analyses were restricted to high-confidence entries, and protein- or phosphosite-level data were collapsed to one representative entry per gene for downstream gene-level analyses. Dataset-specific background universes were defined from the corresponding detected gene sets after identifier mapping. Detailed thresholds, comparison schemes, and additional analysis-specific tools, including Enrichr/COMPARTMENTS and IPA, are described in the relevant subsections below. Code availability is described in the Data Availability Statement.

**S2.2. Global Whole Cell Proteomic Data Analysis**

Global whole-cell proteomics data were analyzed to assess overall variance structure, identify differentially abundant proteins, characterize enriched functional categories, and visualize curated motility-related protein changes. Principal component analysis (PCA) was performed on Log2(x+1)-transformed normalized protein abundance values. Proteins with missing values in all samples were excluded, remaining missing values were imputed using the per-protein median, and zero-variance proteins were removed prior to PCA. Differential protein abundance was visualized using volcano plots, with significance defined as an adjusted p-value < 0.05 and |Log2 FC| ≥ 1 (equivalent to fold change ≥ 2 or ≤ 0.5). Volcano analysis was restricted to high-confidence master protein entries.

To characterize the spatial reorganization of the global proteome, subcellular localization enrichment analysis was performed. Protein lists with altered abundance, represented by corresponding gene symbols, were generated using the aforementioned |Log2 FC| ≥ 1 and p-value < 0.05 thresholds, and proteins with increased and decreased abundance were analyzed separately. Gene lists were submitted to the Enrichr [8-10] web-based platform to query the COMPARTMENTS Curated (2025) [11] database. Organelle and vesicular enrichment significance was determined based on the adjusted p-value, and significant terms were visualized as −Log10(p-value).

For the motility-related analysis, a curated Motility Hardware panel was constructed by matching the detected proteome to defined gene sets from the Adhesome resource [12] and MSigDB [13, 14] functional networks, including the Leading Edge, Cytoskeleton, Trafficking, and Endosomal categories. Candidates were selected using a robustness-first strategy that prioritized consistent detection across replicates, higher normalized abundance, and greater stability across control replicates, rather than proteins with the largest fold changes. Relative change was visualized as Log2 fold change in a heatmap, with the color scale capped at ±2.0 to improve visual contrast for the subtle, relevant changes observed (approximately −0.28 to +0.68).

**2.3. Membrane Enriched Subcellular Proteomic Analysis.**

Membrane-enriched proteomics data were analyzed to evaluate fractionation purity, identify compartment-specific protein changes, characterize enriched functional categories, and examine membrane-associated receptor candidates. Analyses were generally restricted to high-confidence master protein entries, and protein-level results were collapsed to one representative entry per gene for downstream gene-level analyses. PCA was performed on normalized abundance values from high-confidence master proteins across the biological TMT channels after exclusion of the pooled/reference channel; proteins detected in at least 70% of samples were retained, Log2(x+1) transformation was applied, missing values were imputed by row median, and centered/scaled PCA was used to visualize overall sample relationships by scree plot and PC score plots.

Fractionation purity was evaluated by Gene Ontology Cellular Component enrichment analysis of the top 100 most abundant proteins using a dataset-specific background universe derived from the membrane proteomics dataset after mapping detected genes to Entrez identifiers.

To identify biologically relevant shifts in the membrane compartment, differential abundance was visualized using membrane TMT volcano plots. Statistical significance was defined as Welch p-value < 0.05 together with two fold change thresholds: a standard cutoff of |Log2 FC| ≥ 0.26 (approximately 1.2-fold) and a high-stringency cutoff of |Log2 FC| ≥ 0.58 (approximately 1.5-fold).

To characterize functional reorganization, a Biological Process Gene Ontology over-representation analysis was performed using the clusterProfiler package in R. Differential-abundance gene lists were generated after collapsing to one entry per gene by retaining the smallest Welch p-value, with ties resolved by larger absolute Log2 fold change. Enrichment was performed separately for upregulated, downregulated, and combined gene sets at both thresholds using a dataset-specific background universe.

**2.4. Quantitative Temporal Phosphoproteomic Data Analysis**

Quantitative temporal phosphoproteomic data were analyzed to characterize signaling dynamics across time points and concentrations, identify differentially regulated phosphosites, and evaluate functional pathway reorganization. PCA was performed on high-confidence phosphosite-level features derived from scaled abundance values after collapsing duplicate site identifiers by median abundance and retaining phosphosites observed in at least 50% of the samples; values were Log2(x+1)-transformed, missing values were imputed by row median, and centered/scaled PCA was used to visualize overall sample relationships across time points and concentrations. Signaling dynamics were quantified at 15 minutes, 1 hour, and 4 hours. Downstream phosphosite-counting, residue-composition, and overlap analyses were performed at the level of unique absolute phosphosites to avoid repeated counting of the same site across multiple peptide entries. Unique phosphosites were tracked using a robust site identifier based on the corresponding protein-mapped site annotation, and significance was defined as |Log2 FC| ≥ 0.58 and adjusted p-value < 0.05.

To capture the complexity of phosphoproteomic modulation across time points and concentrations, a comprehensive pairwise analysis was performed. Differential phosphosite-abundance landscapes were visualized using volcano plots for drug-versus-control, time-evolution, and dose comparisons. Candidates were evaluated using a standard threshold of |Log2 FC| ≥ 0.26 (approximately 1.2-fold change) and a stringent threshold of |Log2 FC| ≥ 0.58 (approximately 1.5-fold change), together with adjusted p-value < 0.05.

Functional interpretation was also performed using Ingenuity Pathway Analysis (IPA). Phosphosite-level abundance tables collapsed to one row per gene were analyzed using Core Analysis and Comparison Analysis. For pathway-centric interpretation, a threshold of |Log2 FC| ≥ 0.5 was used to capture broader coordinated signaling changes, whereas a target-centric threshold of |Log2 FC| ≥ 1.0 was used to identify stronger signaling shifts. Results were used to evaluate Canonical Pathways, Upstream Regulators, and Disease and Bio Functions across different time points and concentrations. Canonical Pathways were filtered for |z-score| ≥ 2 and Benjamini–Hochberg-adjusted p-value < 0.05.

For comparative pathway profiling, two parallel contrasts were analyzed: 1 µM versus Control and 10 µM versus Control. To reduce enrichment bias from multiple phosphopeptides derived from the same protein, phosphosite-level measurements were collapsed to the gene level by retaining one representative entry per gene before enrichment analysis. Hit genes were defined using |Log2FC| ≥ 0.58 and Benjamini–Hochberg-adjusted p-value < 0.05.

To track signal propagation, sites were classified dynamically across time points as ON_UP (Log2 FC ≥ 0.58, adjusted p-value < 0.05), ON_DOWN (Log2 FC ≤ −0.58, adjusted p-value < 0.05), or OFF.

Kinase-substrate enrichment analysis was performed in R using KSEAapp. High-confidence phosphopeptides were formatted for KSEAapp using phosphosite annotations, adjusted p-values, and fold-change values, with log2 FC converted to linear fold changes before analysis. KSEA scores and kinase–substrate support tables were generated using the KSEAapp kinase–substrate annotation database with NetworKIN filtering enabled and a NetworKIN cutoff of 5.

To evaluate MAPK14/p38-related adaptive signaling at 4 hours, MAPK14 phosphosite-level changes were first examined, with particular attention to activation-loop T180 and Y182 phosphorylation entries. In parallel, MAPK14 KSEA z-scores and p-values were used to assess whether annotated MAPK14 downstream substrates showed a coordinated phosphorylation response. MAPK14-linked substrate genes from the kinase–substrate support table were then examined individually and visualized by phosphoproteomic Log2 FC after collapsing to one representative phosphopeptide entry per gene.
