## Supplementary figures and images for "Integrative Proteomic Analysis Implicates Inhibition of Intracellular Protein Trafficking in Therapy-Induced Migrastasis in Prostate Cancer"

### 1a_std_up_BP_dotplot.pdf

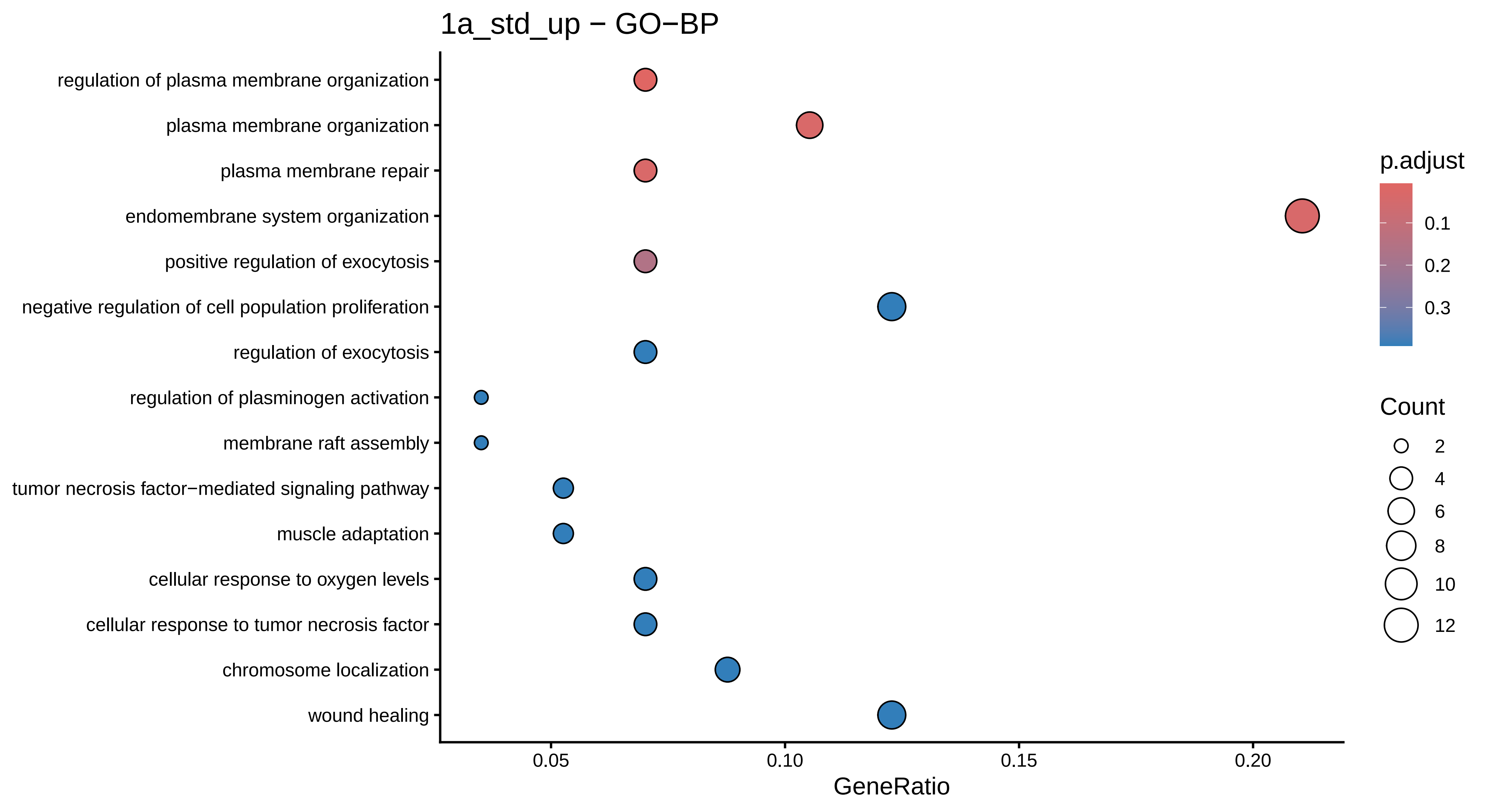

### 1a_std_up_BP_dotplot.png

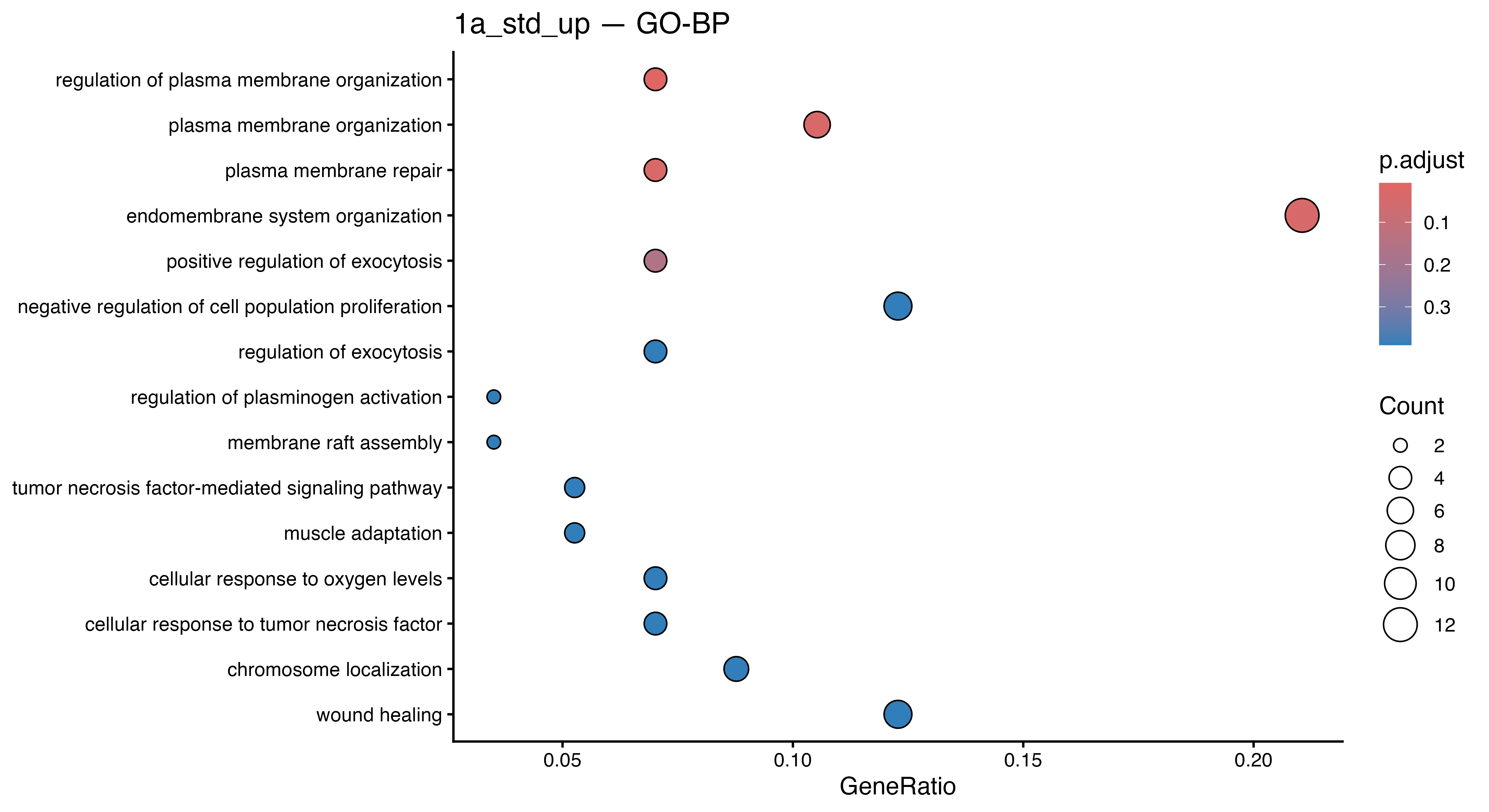

### 1b_std_down_BP_dotplot.pdf

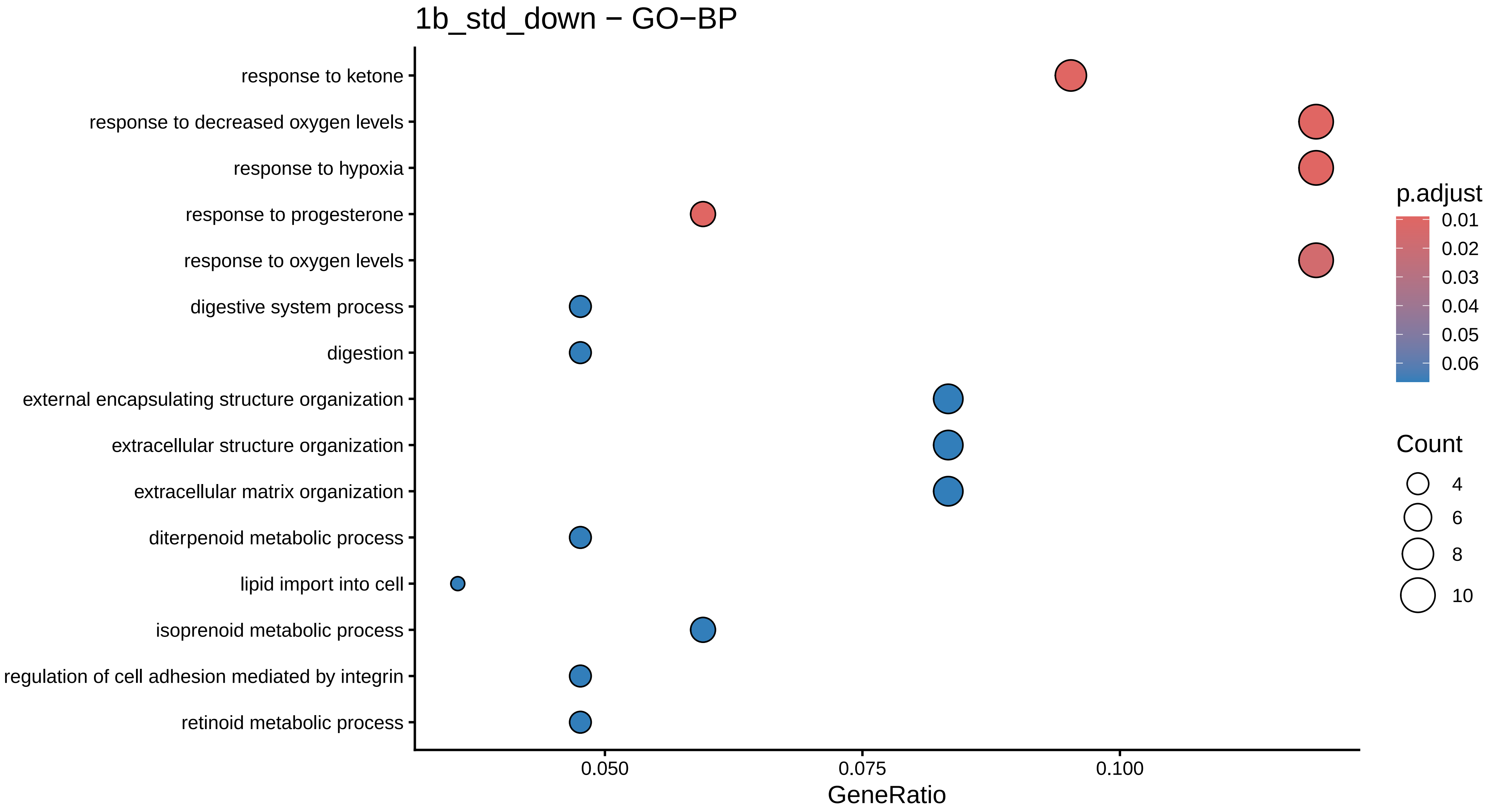

### 1b_std_down_BP_dotplot.png

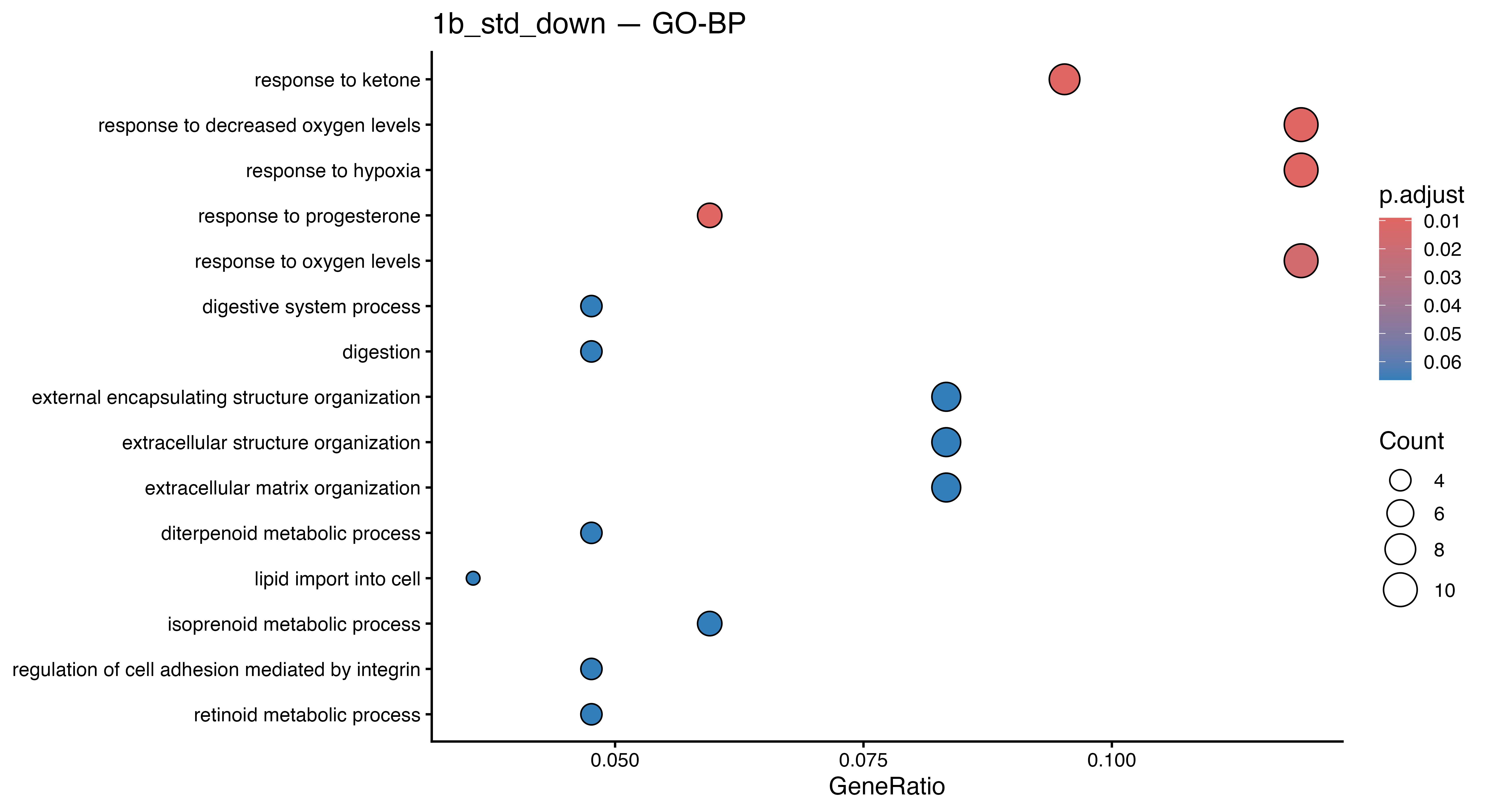

### 1hr_cutoff0.5_1hrDiffCon_Comparison Analysis(visualize BH p-value)_expaned header.png

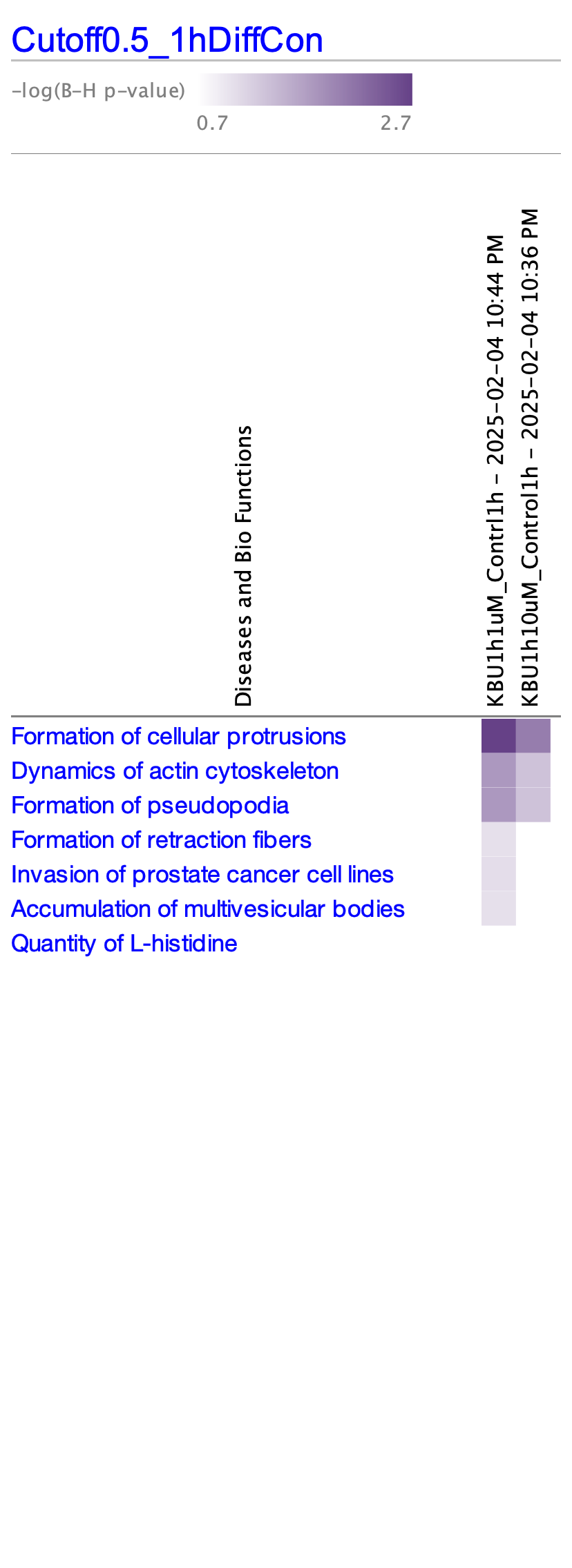

### 1hr_cutoff0.5_1hrDiffCon_Comparison Analysis(visualize BH p-value)_not expaned header.png

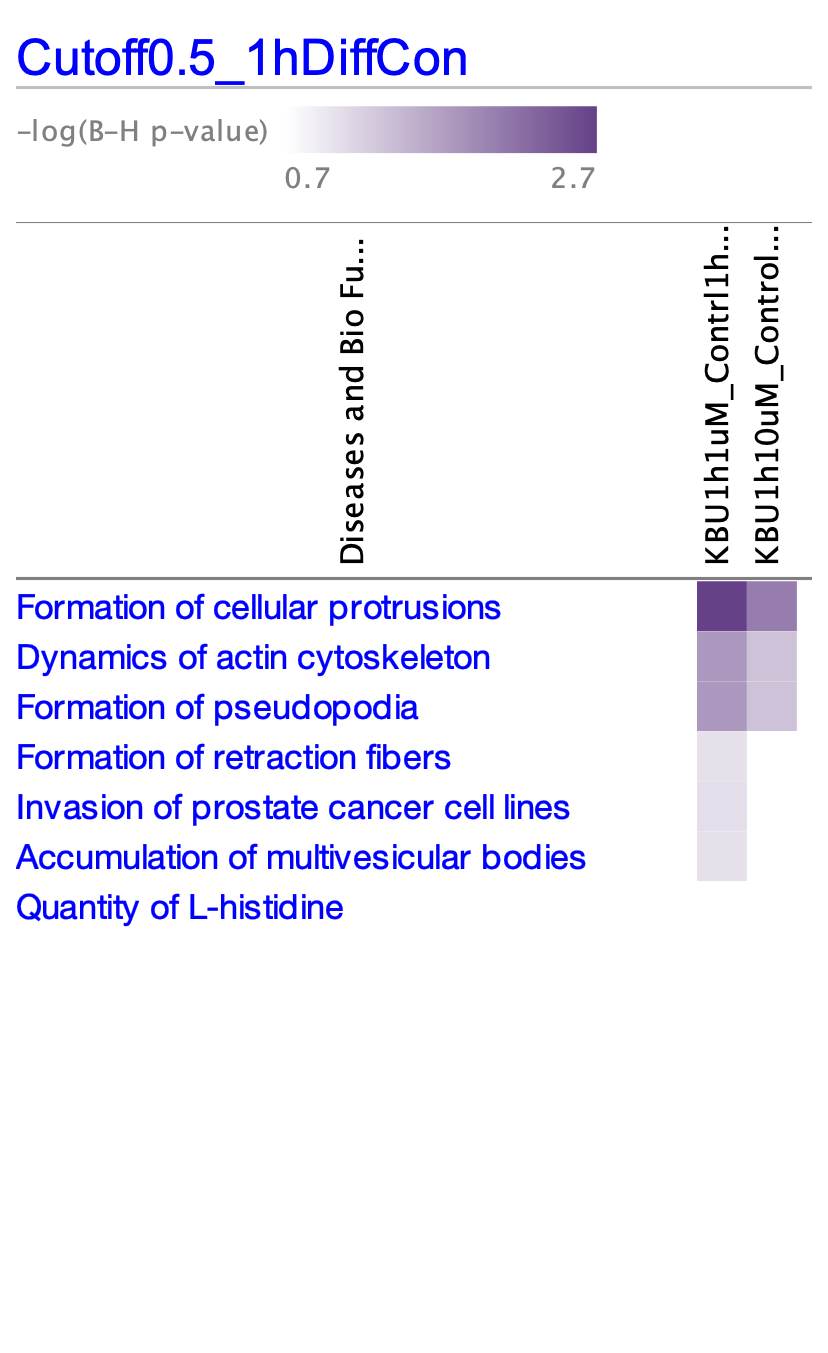

### 1hr_cutoff1_1hrDiffCon_Comparison Analysis(visualize BH p-value)_ expaned header.png

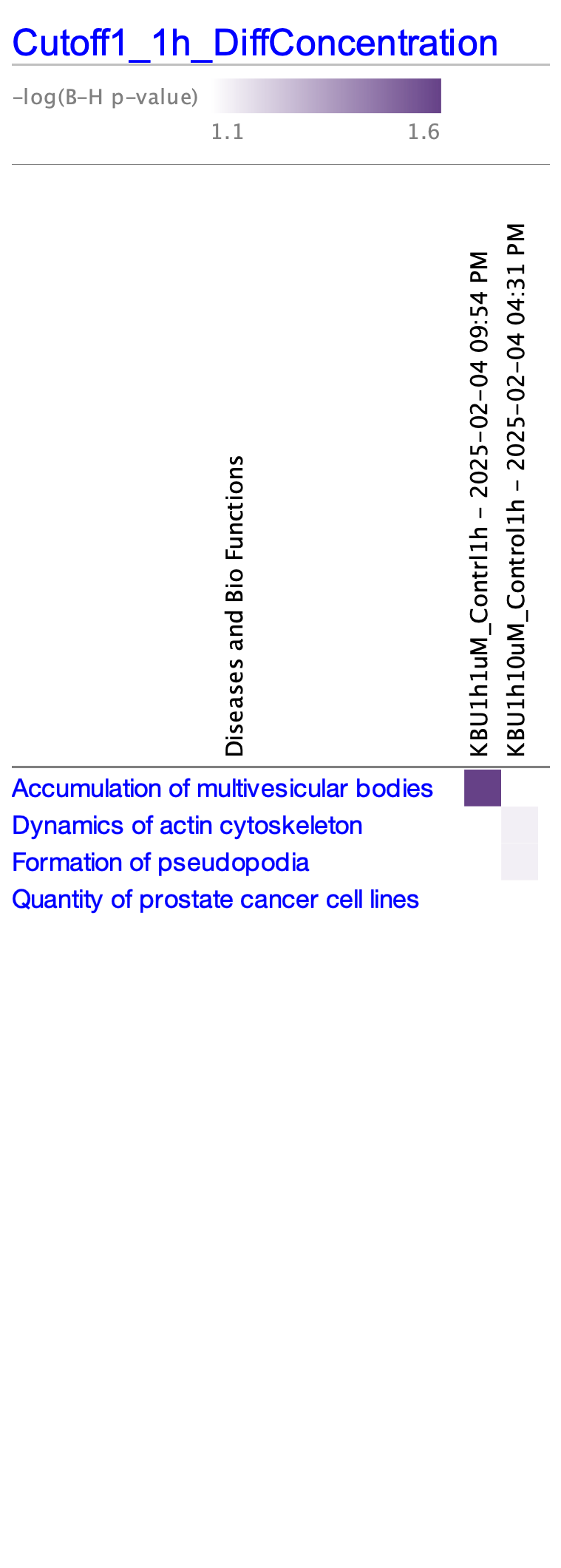

### 1hr_cutoff1_1hrDiffCon_Comparison Analysis(visualize BH p-value)_not expaned header.png

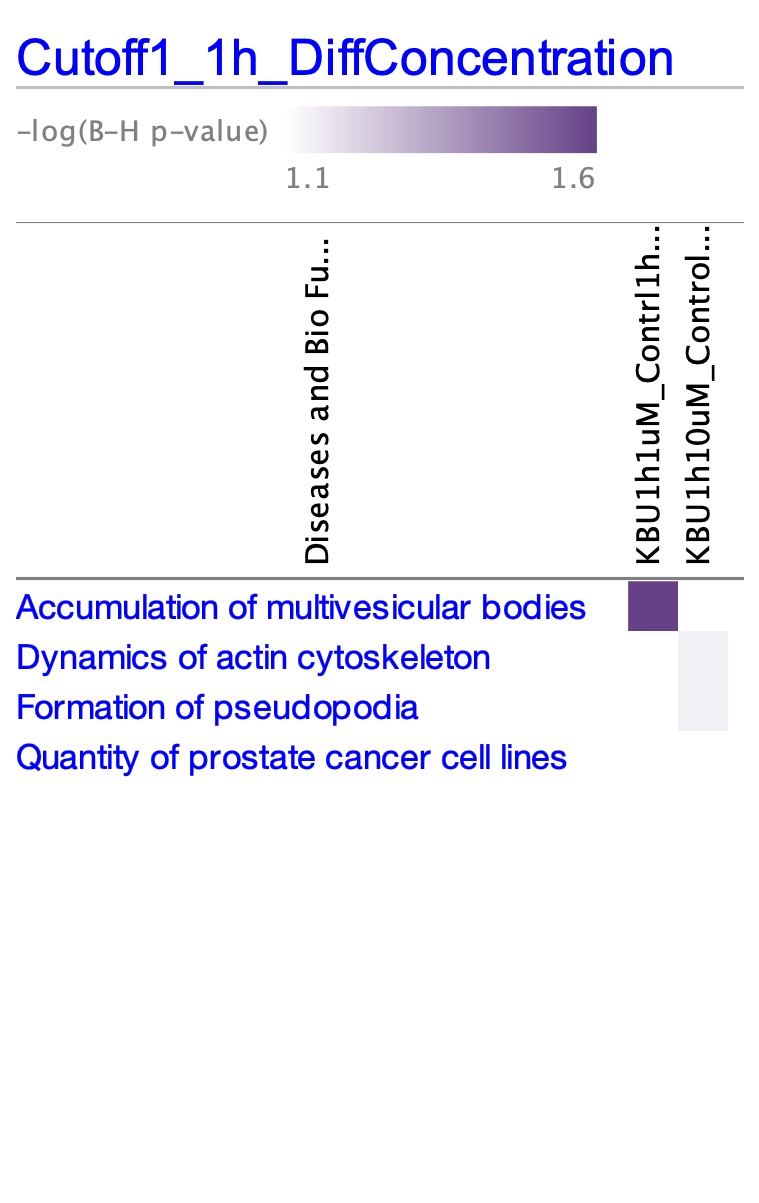

### 2x63x_pc3_DMSO_10UMKBU_3hrs_activated beta1(green)_actin(red)_DMSO_01.tif

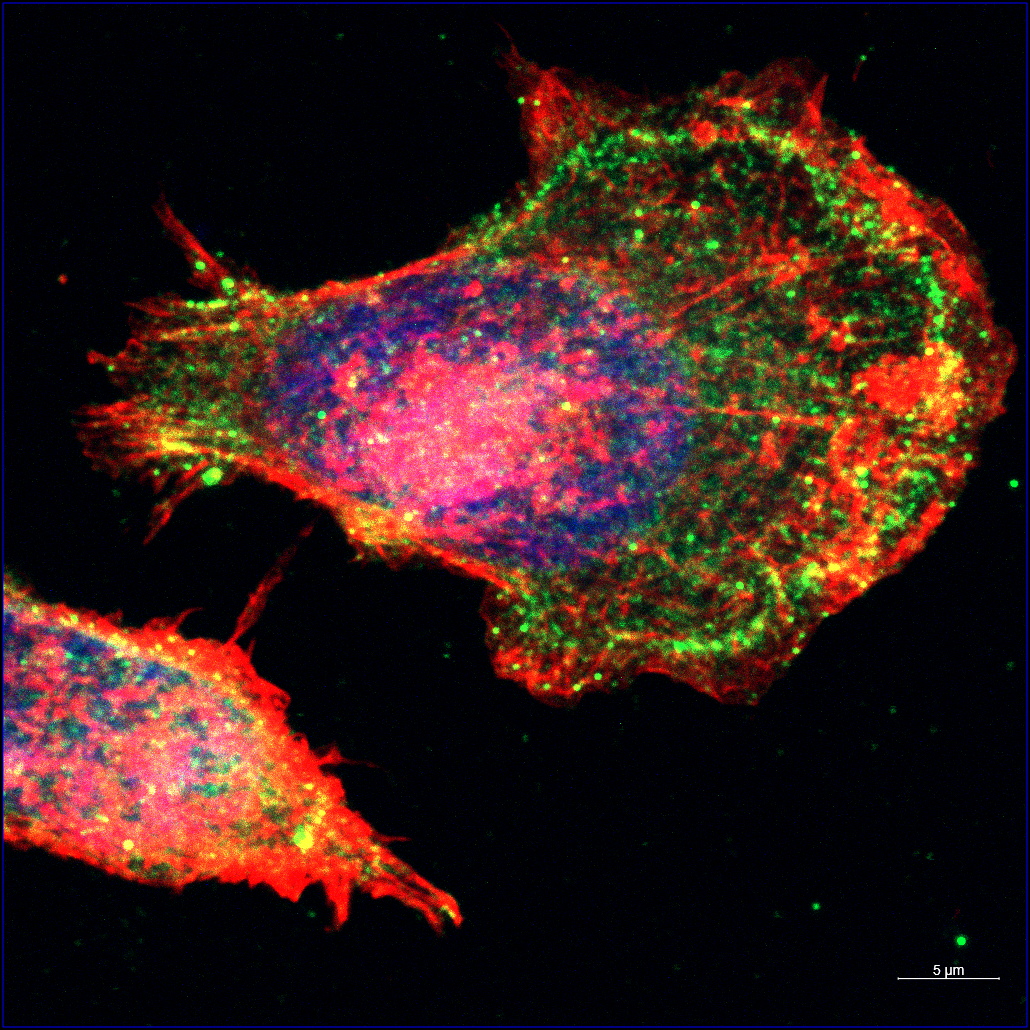

### 2x63x_pc3_DMSO_10UMKBU_3hrs_activated beta1(green)_actin(red)_DMSO_01_AF488-T3.tif

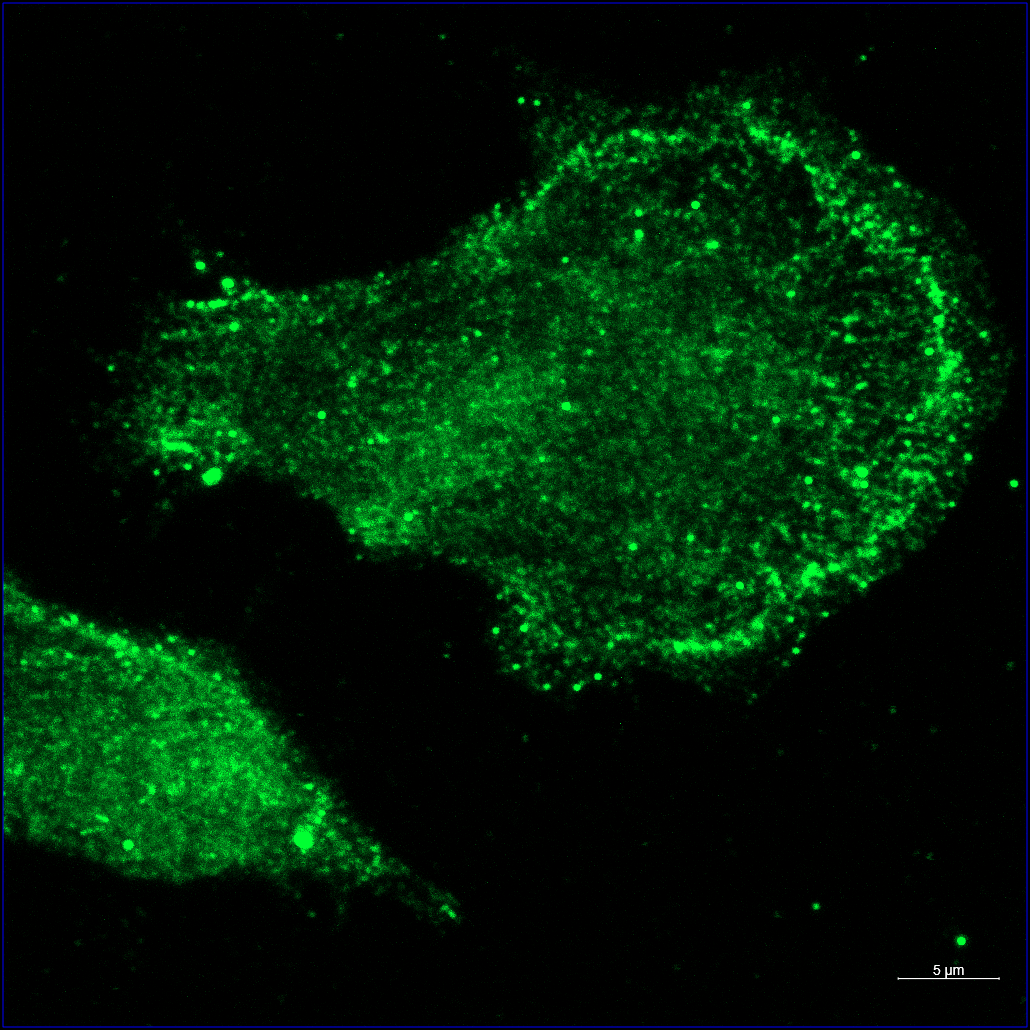

### 2x63x_pc3_DMSO_10UMKBU_3hrs_activated beta1(green)_actin(red)_DMSO_01_AF488-T3_ORG.tif

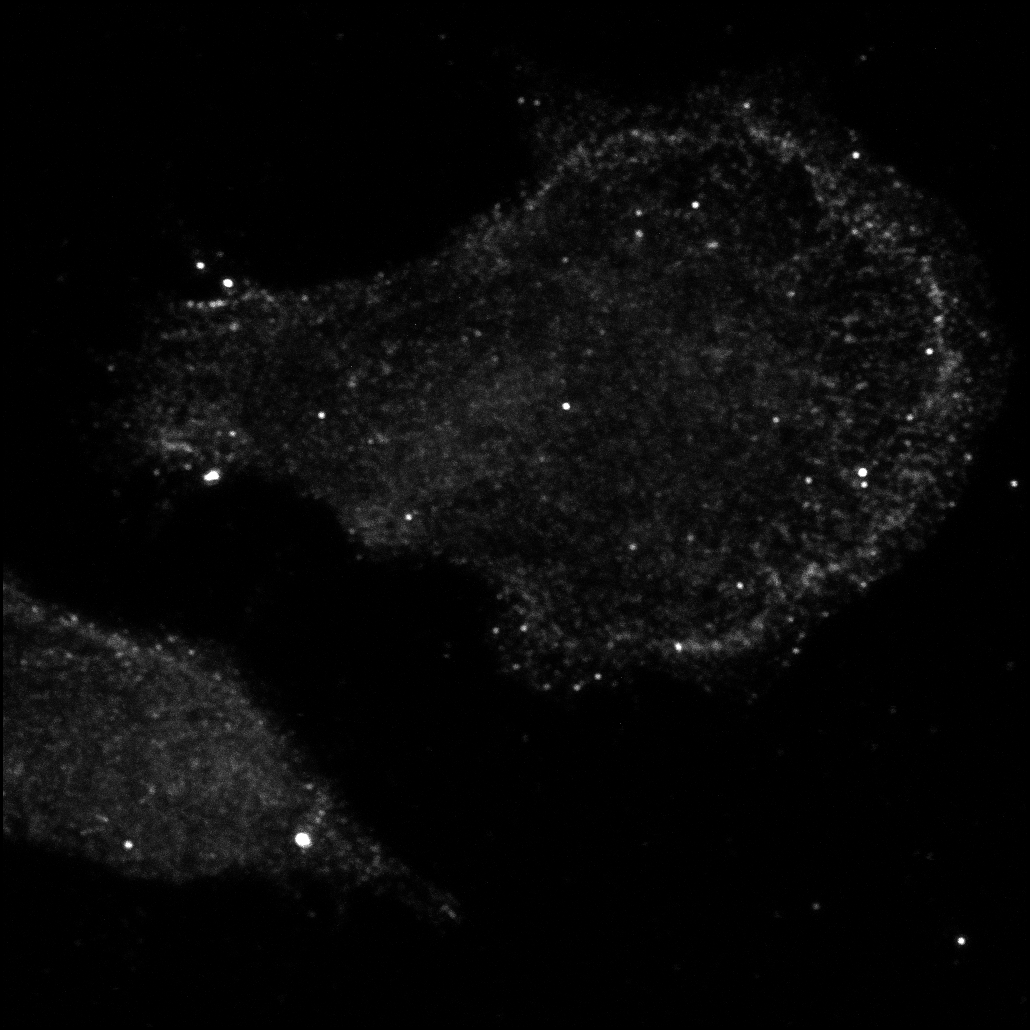

### 2x63x_pc3_DMSO_10UMKBU_3hrs_activated beta1(green)_actin(red)_DMSO_01_AF568-T2.tif

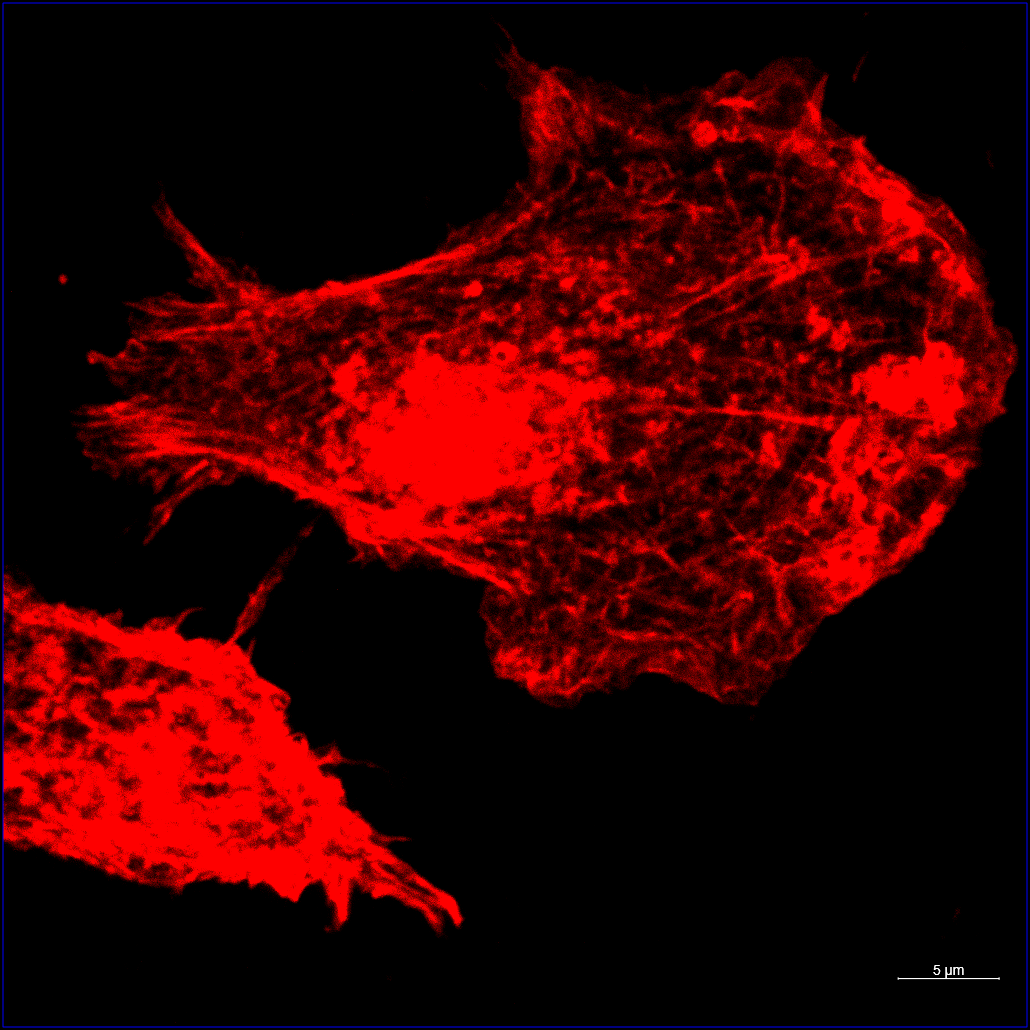

### 2x63x_pc3_DMSO_10UMKBU_3hrs_activated beta1(green)_actin(red)_DMSO_01_AF568-T2_ORG.tif

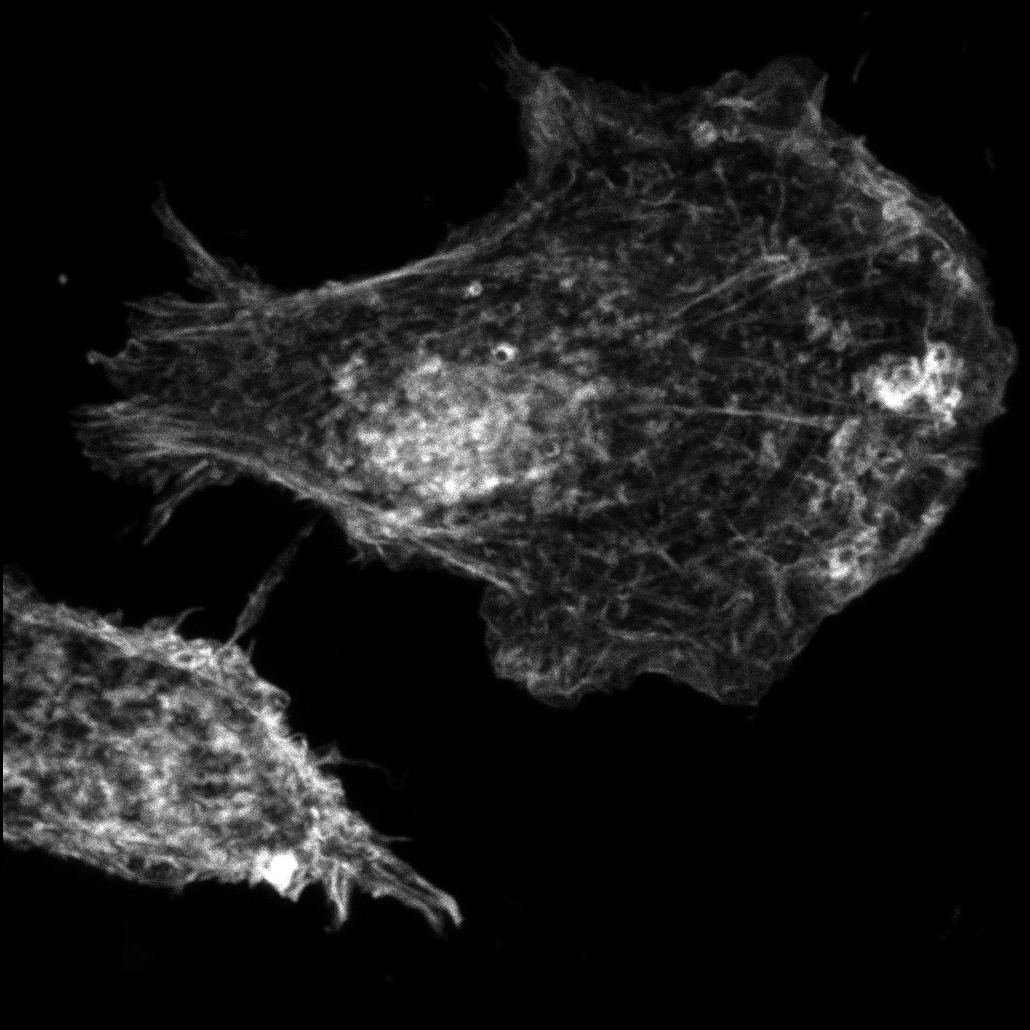

### 2x63x_pc3_DMSO_10UMKBU_3hrs_activated beta1(green)_actin(red)_DMSO_01_DAPI-T4.tif

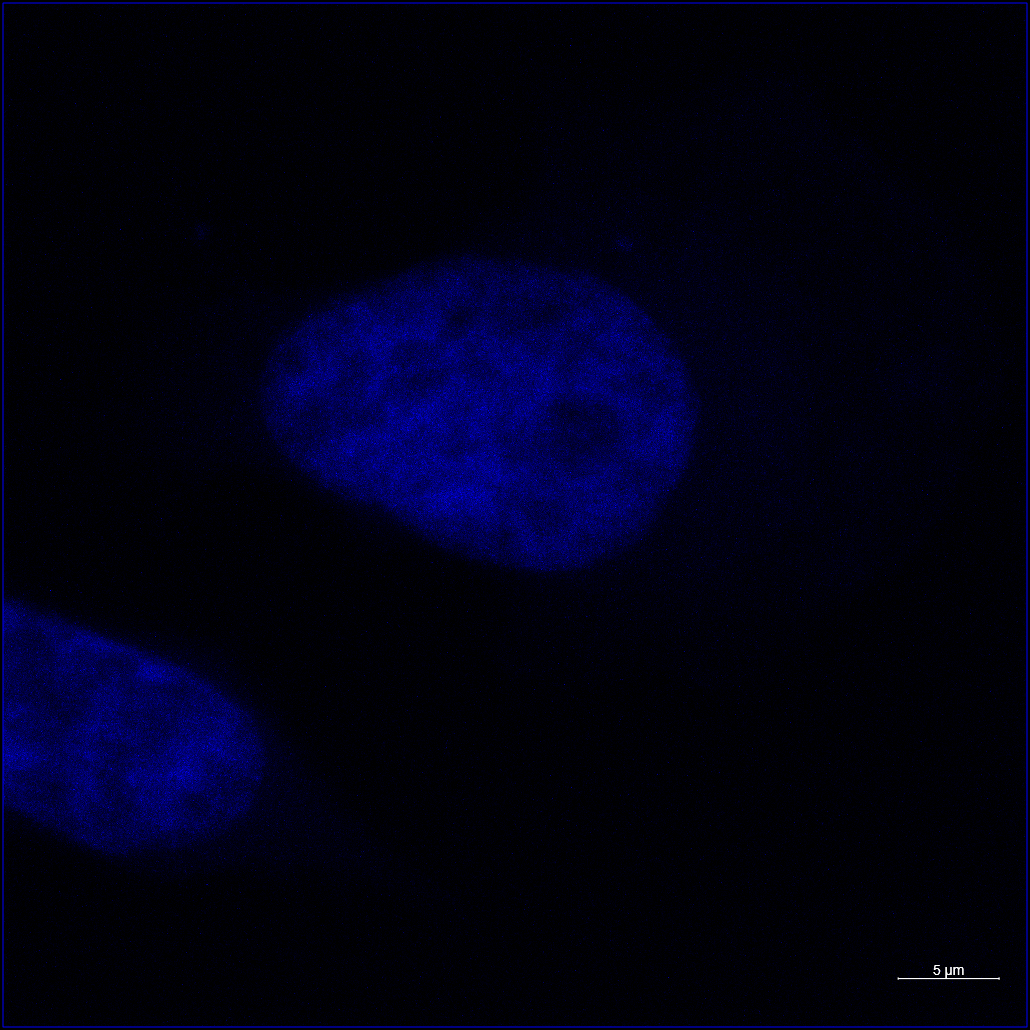

### 2x63x_pc3_DMSO_10UMKBU_3hrs_activated beta1(green)_actin(red)_DMSO_01_DAPI-T4_ORG.tif

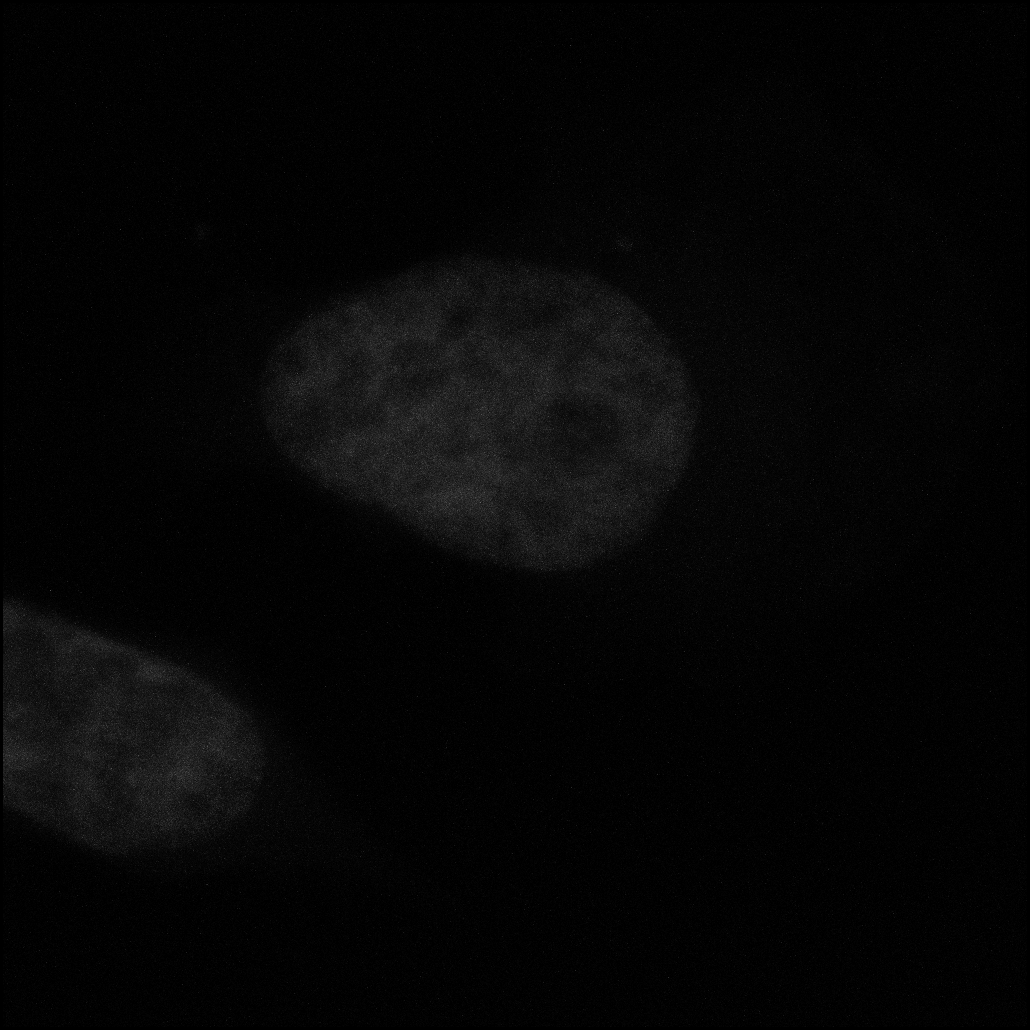

### 2x63x_pc3_DMSO_10UMKBU_3hrs_activated beta1(green)_actin(red)_KBU_02.tif

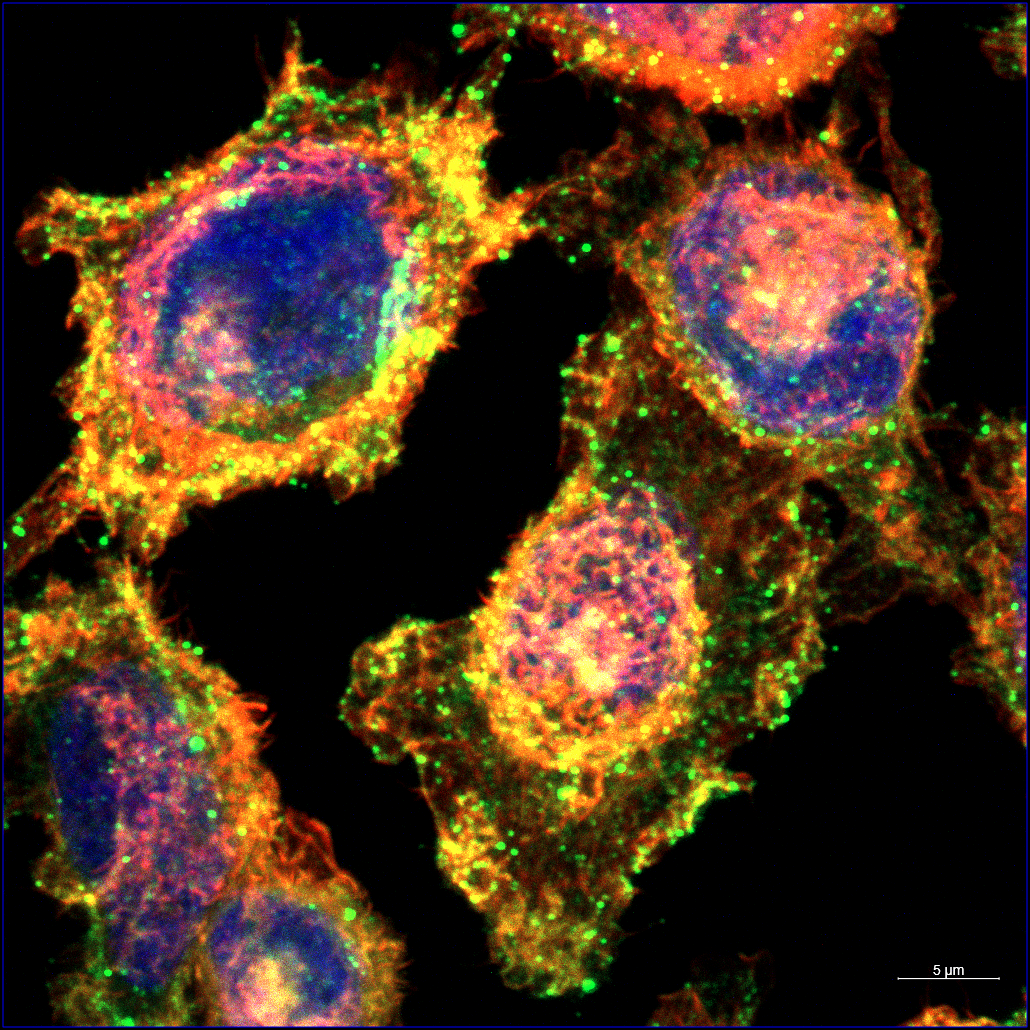

### 2x63x_pc3_DMSO_10UMKBU_3hrs_activated beta1(green)_actin(red)_KBU_02_AF488-T3.tif

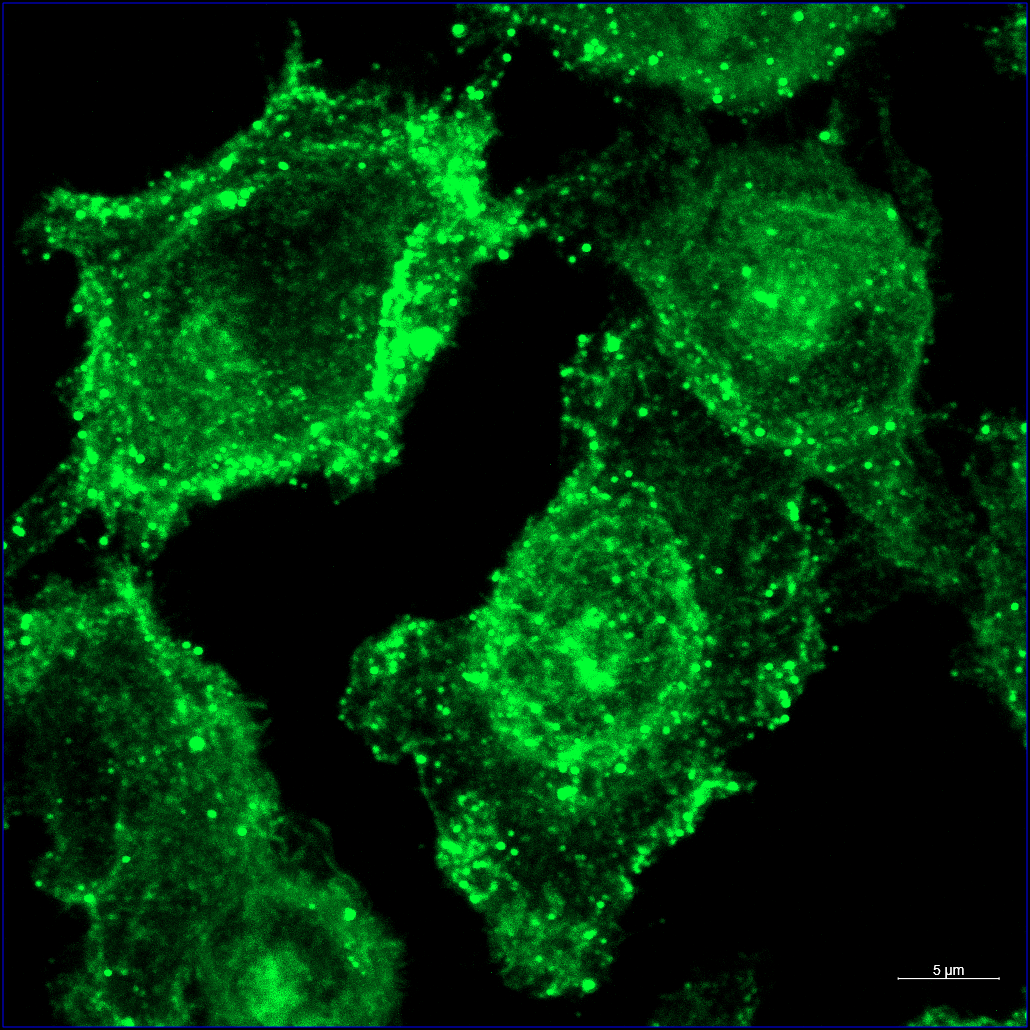

### 2x63x_pc3_DMSO_10UMKBU_3hrs_activated beta1(green)_actin(red)_KBU_02_AF488-T3_ORG.tif

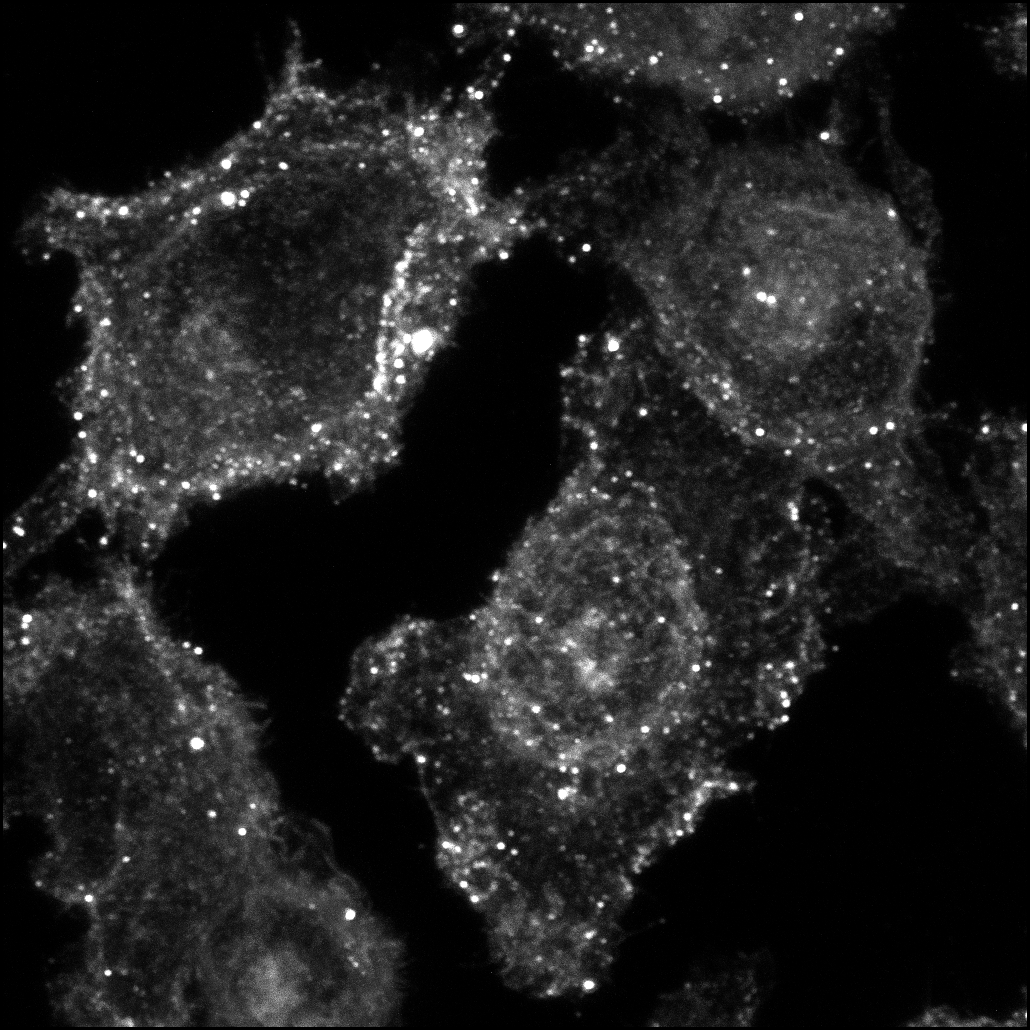

### 2x63x_pc3_DMSO_10UMKBU_3hrs_activated beta1(green)_actin(red)_KBU_02_AF568-T2.tif

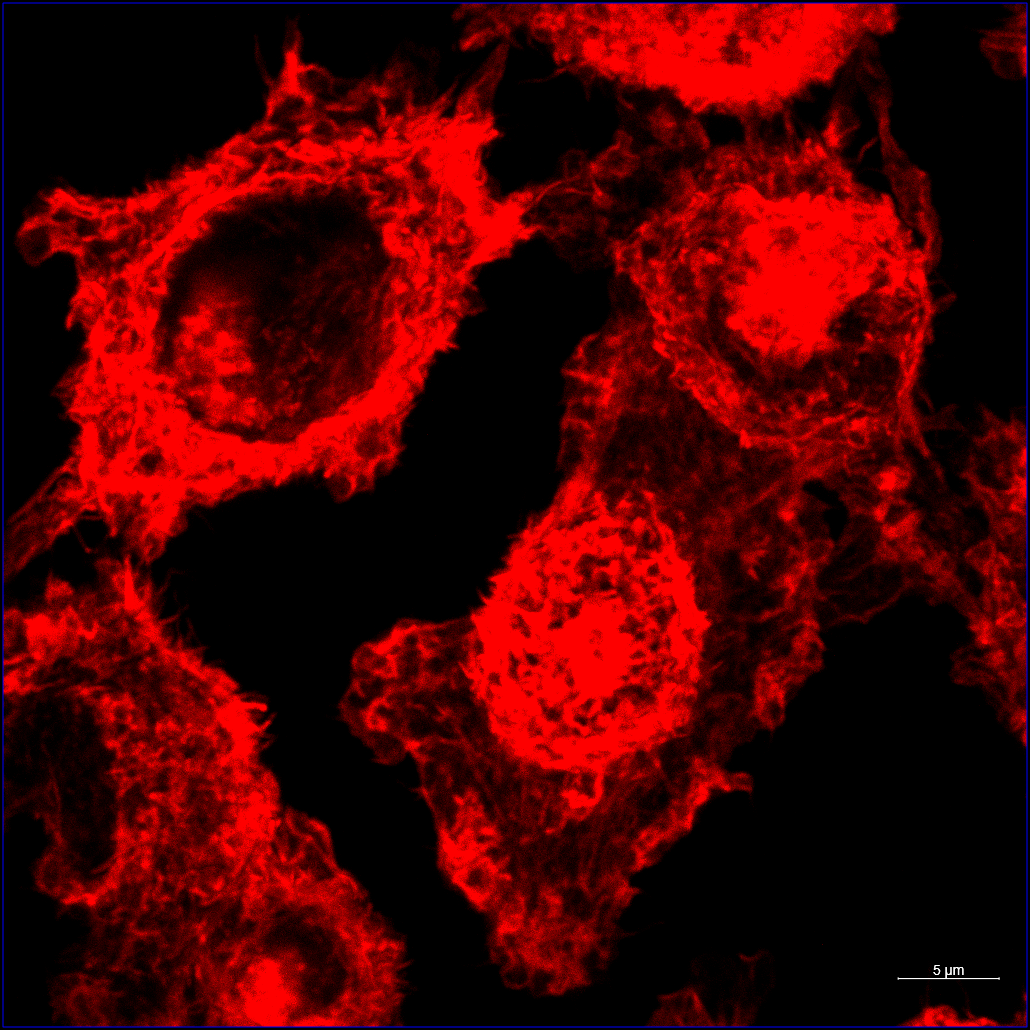

### 2x63x_pc3_DMSO_10UMKBU_3hrs_activated beta1(green)_actin(red)_KBU_02_AF568-T2_ORG.tif

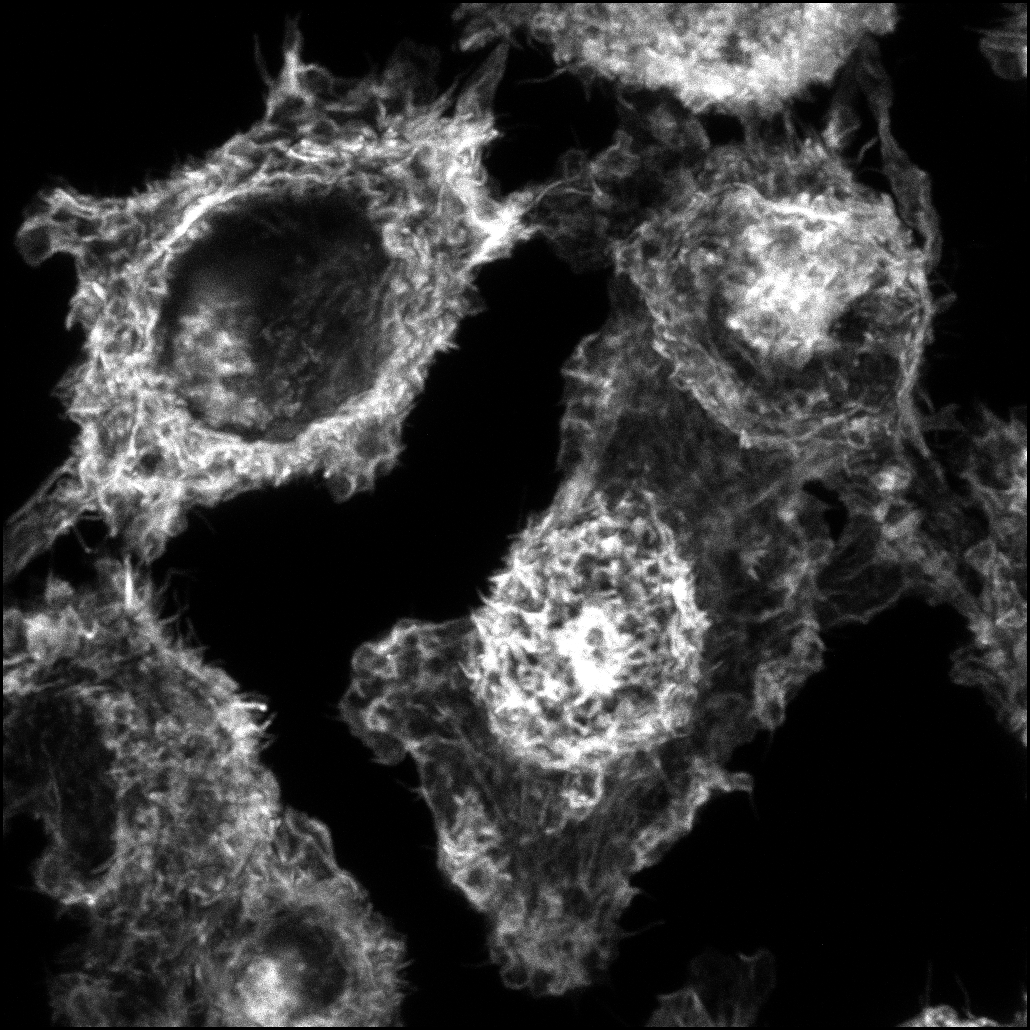

### 2x63x_pc3_DMSO_10UMKBU_3hrs_activated beta1(green)_actin(red)_KBU_02_DAPI-T4.tif

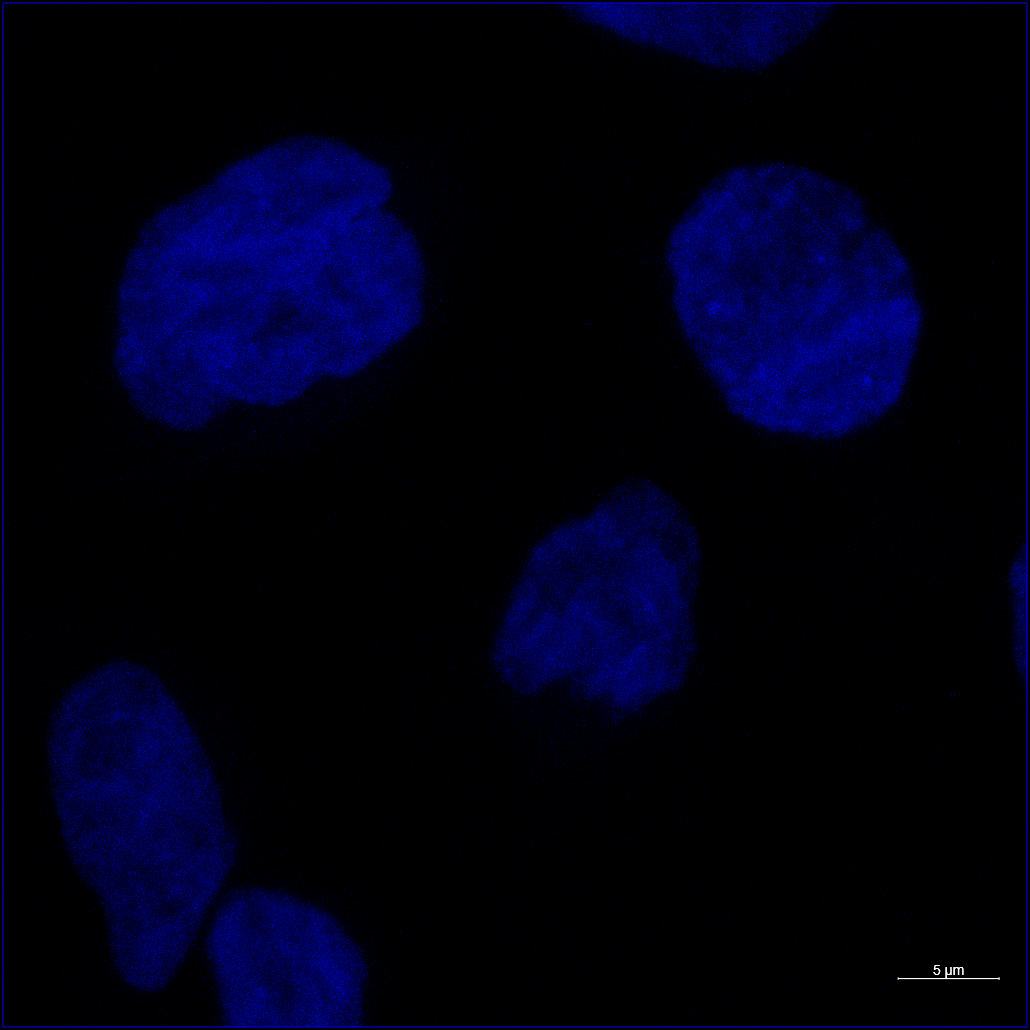

### 2x63x_pc3_DMSO_10UMKBU_3hrs_activated beta1(green)_actin(red)_KBU_02_DAPI-T4_ORG.tif

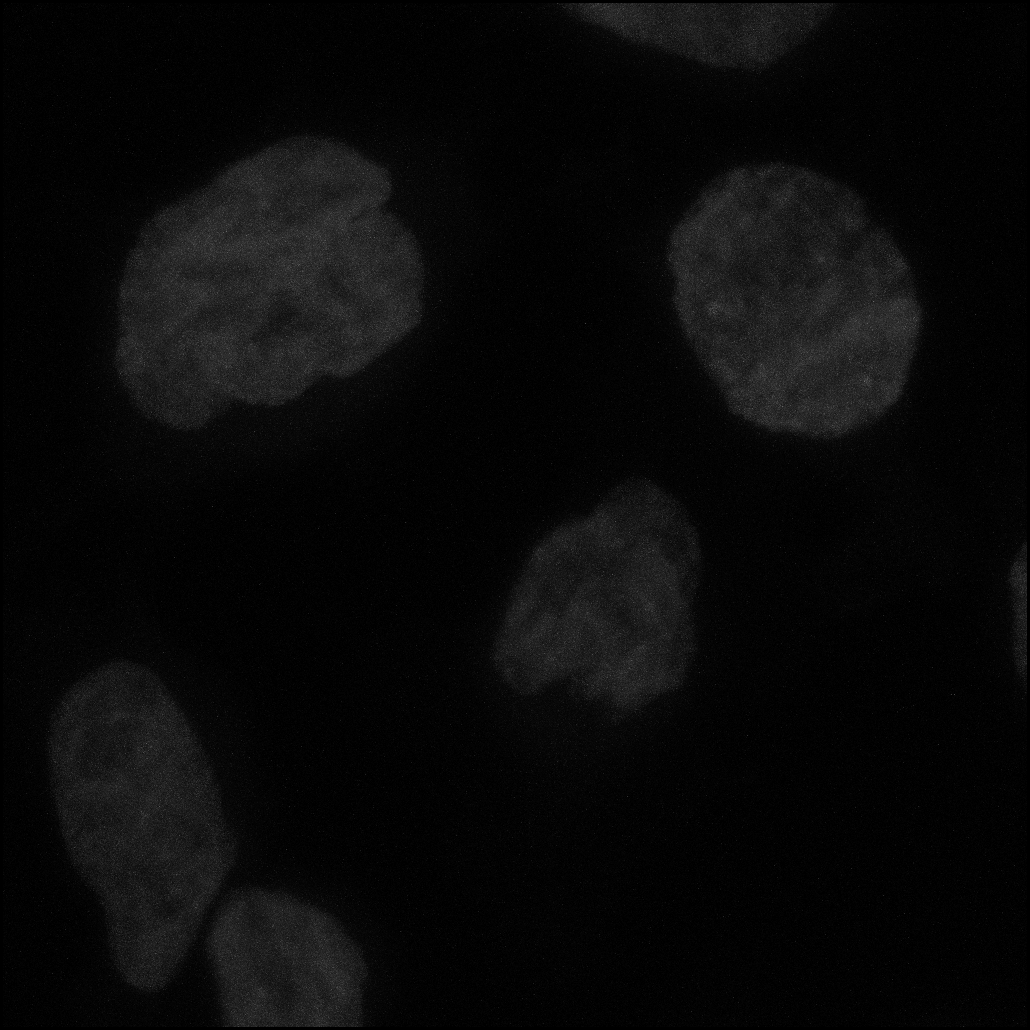

### 15mins_cutoff1_15minDiffCon_Comparison Analysis(visualize BH p-value)_expaned header.png

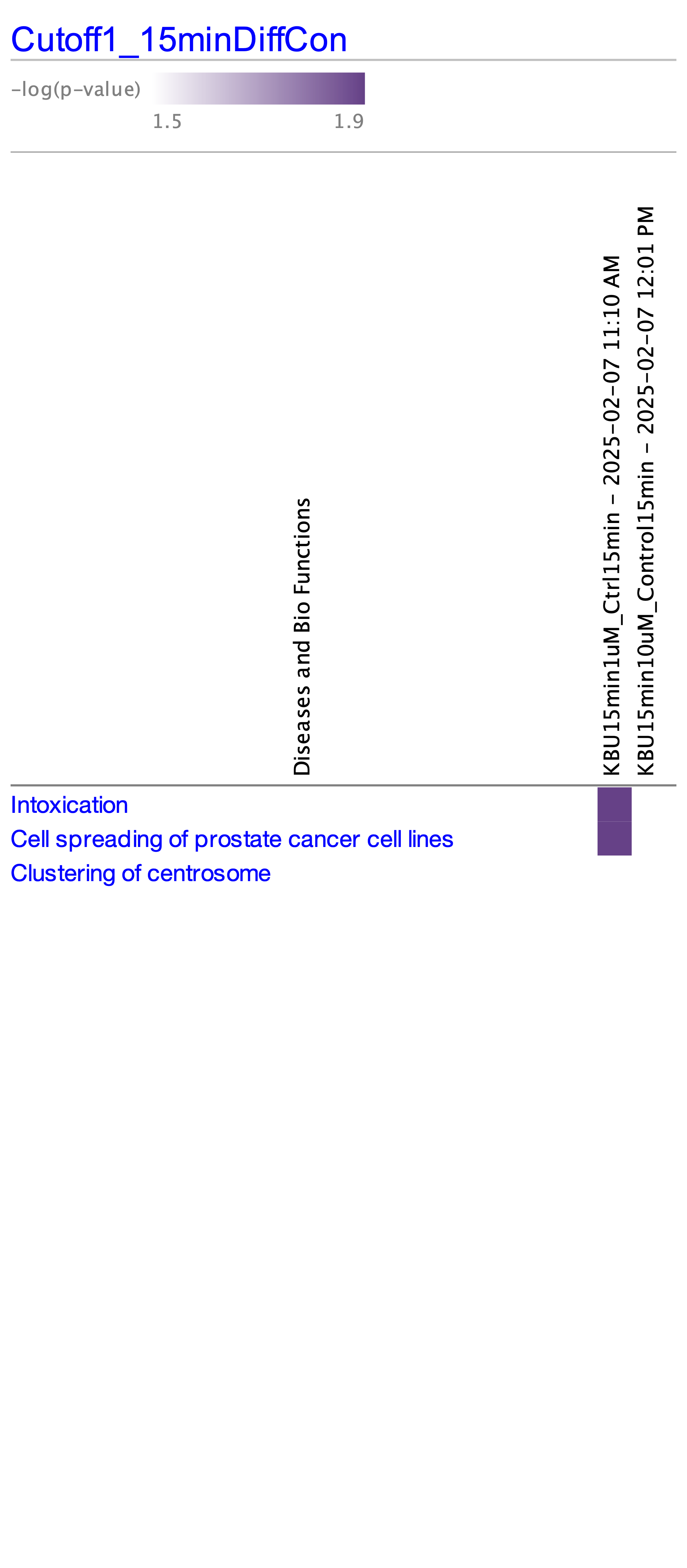

### 15mins_cutoff1_15minDiffCon_Comparison Analysis(visualize BH p-value)_not expaned header.png

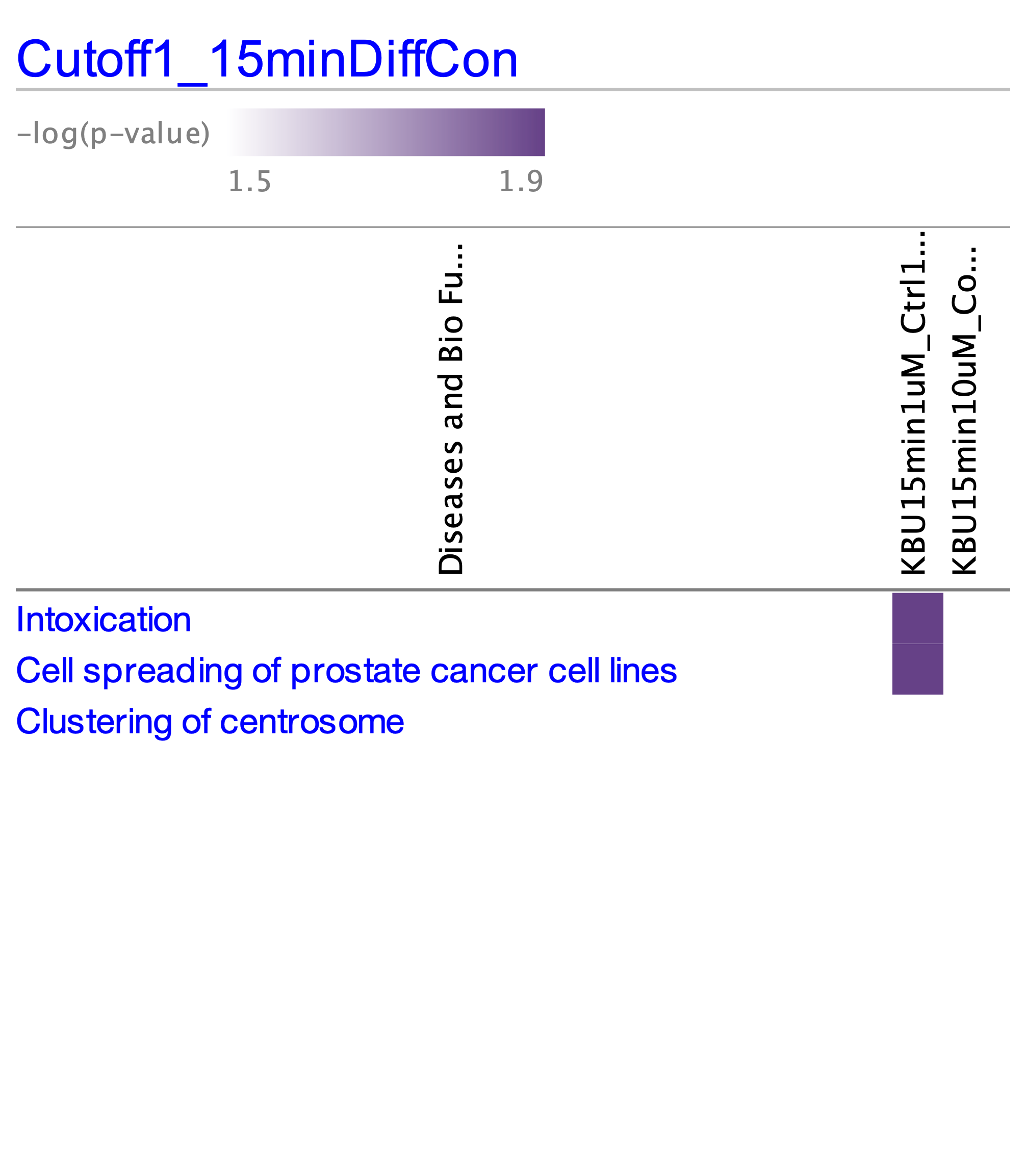

### 40x_1532C_DMSO_10UMKBU_2hrs_activated beta1(green)_actin(red)_kbu_04.tif

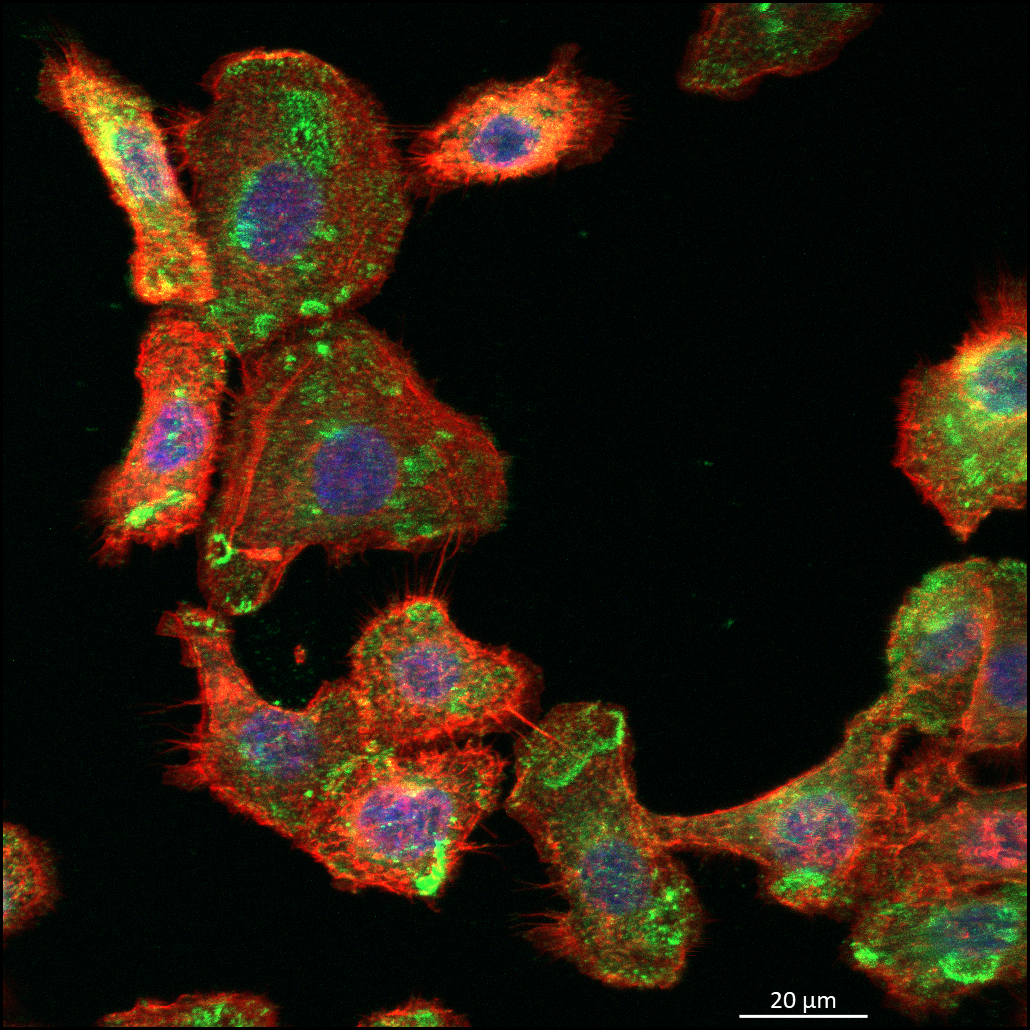

### 40x_1532C_DMSO_10UMKBU_2hrs_activated beta1(green)_actin(red)_kbu_04_AF488-T3.tif

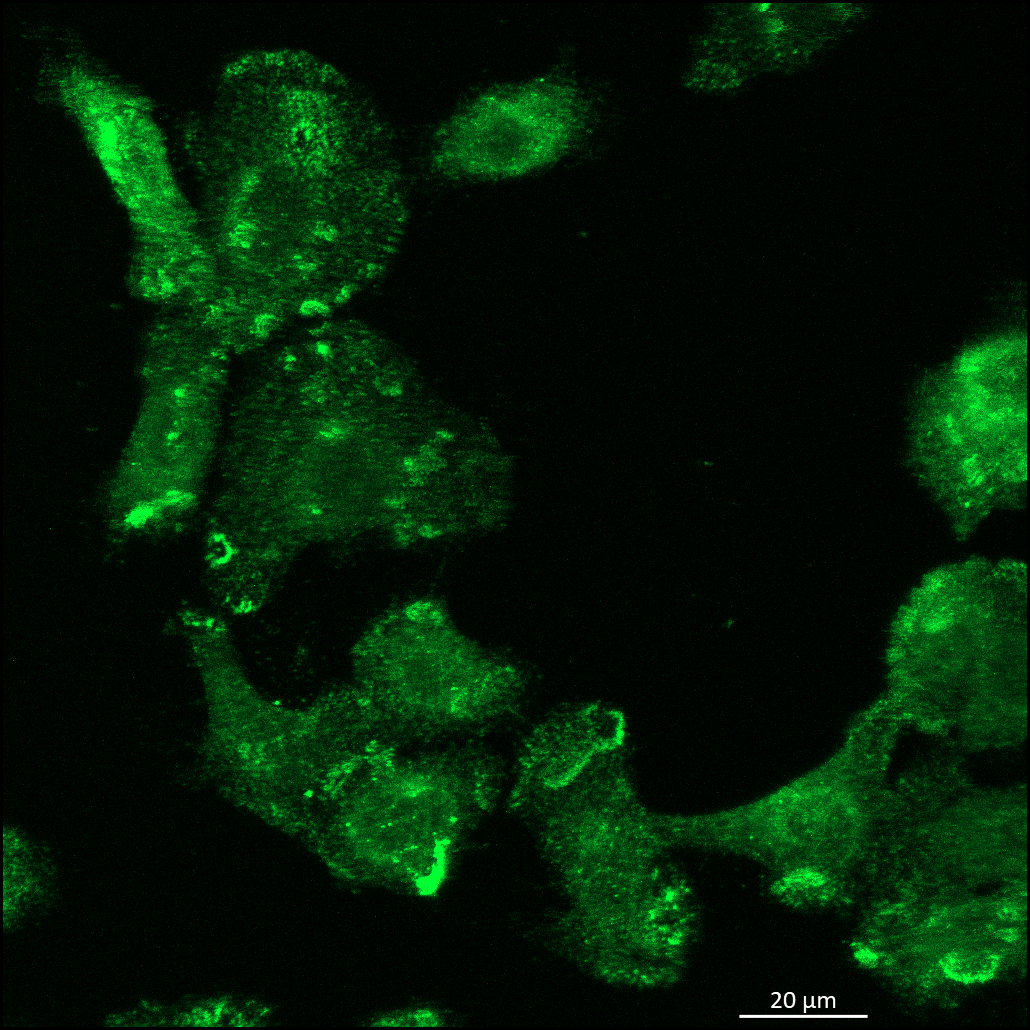

### 40x_1532C_DMSO_10UMKBU_2hrs_activated beta1(green)_actin(red)_kbu_04_AF488-T3_ORG.tif

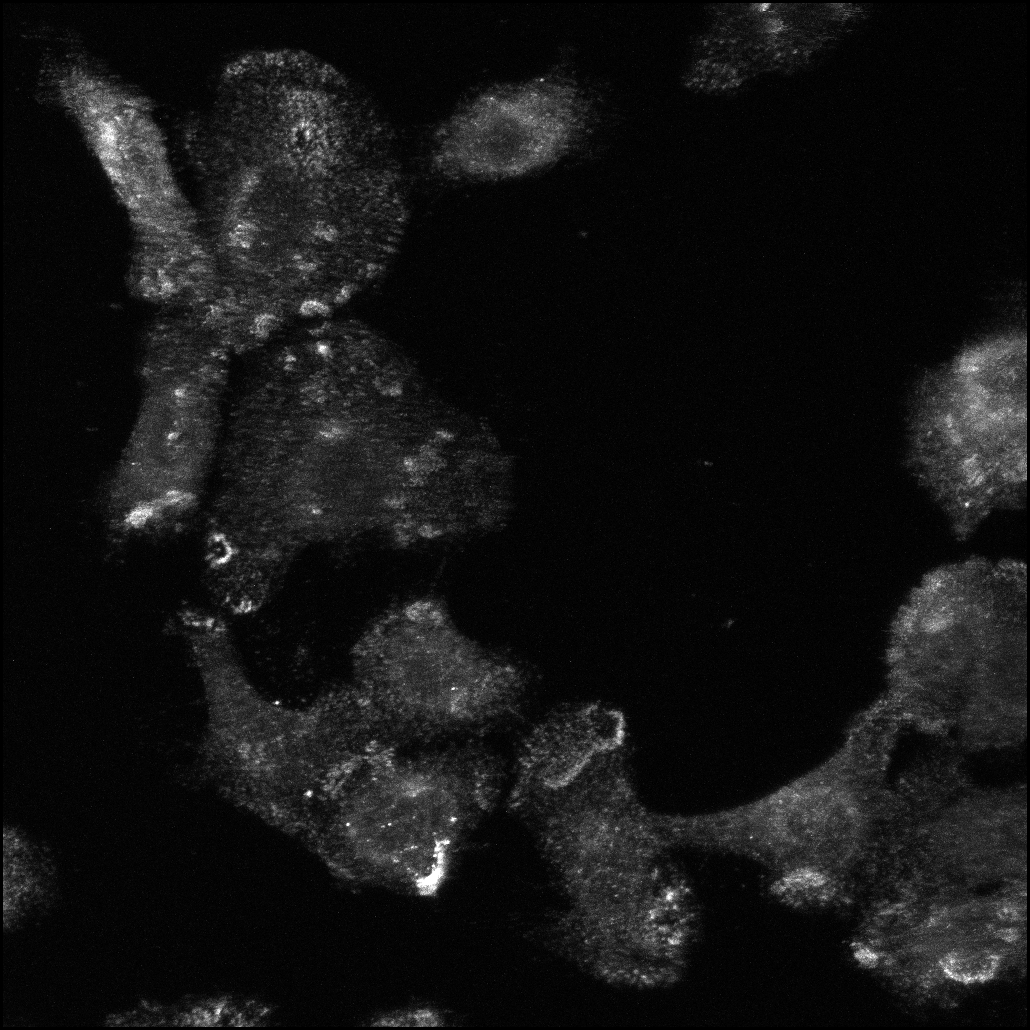

### 40x_1532C_DMSO_10UMKBU_2hrs_activated beta1(green)_actin(red)_kbu_04_AF568-T2.tif

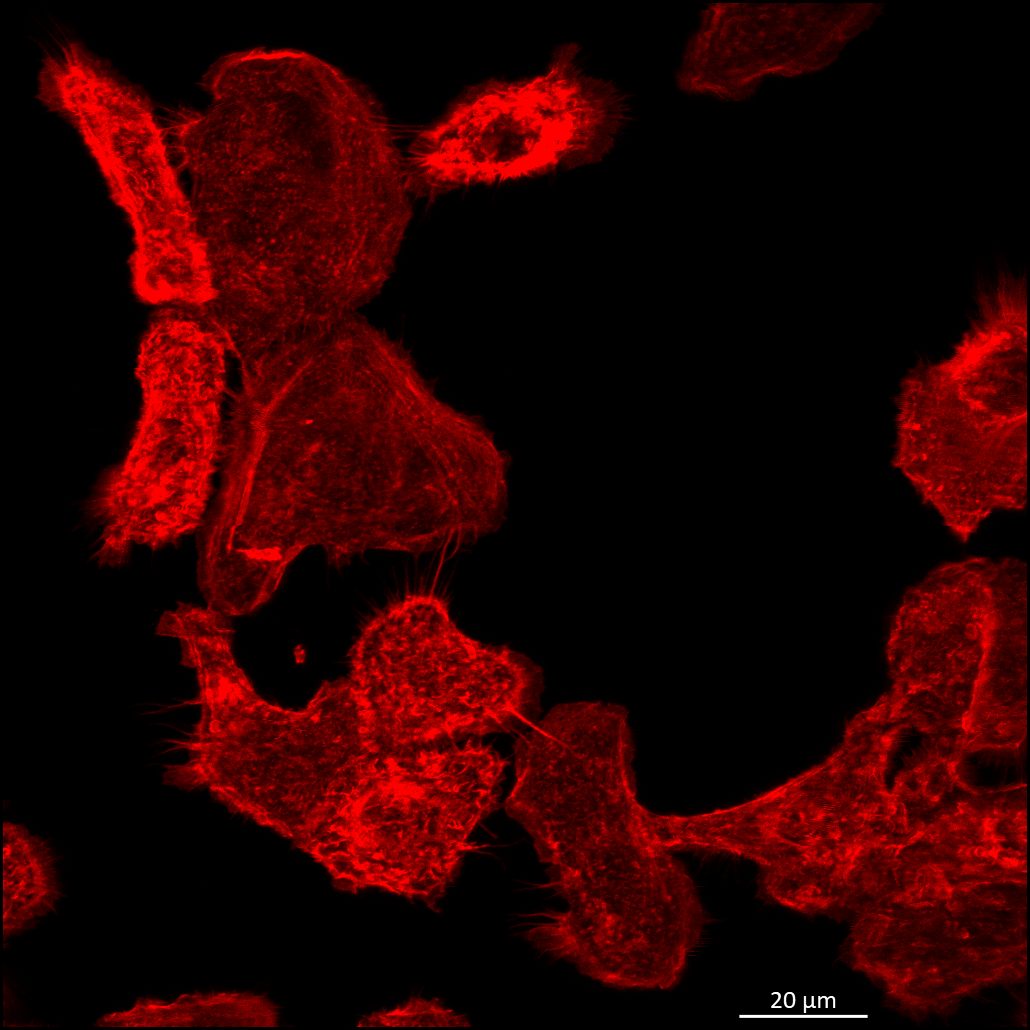

### 40x_1532C_DMSO_10UMKBU_2hrs_activated beta1(green)_actin(red)_kbu_04_AF568-T2_ORG.tif

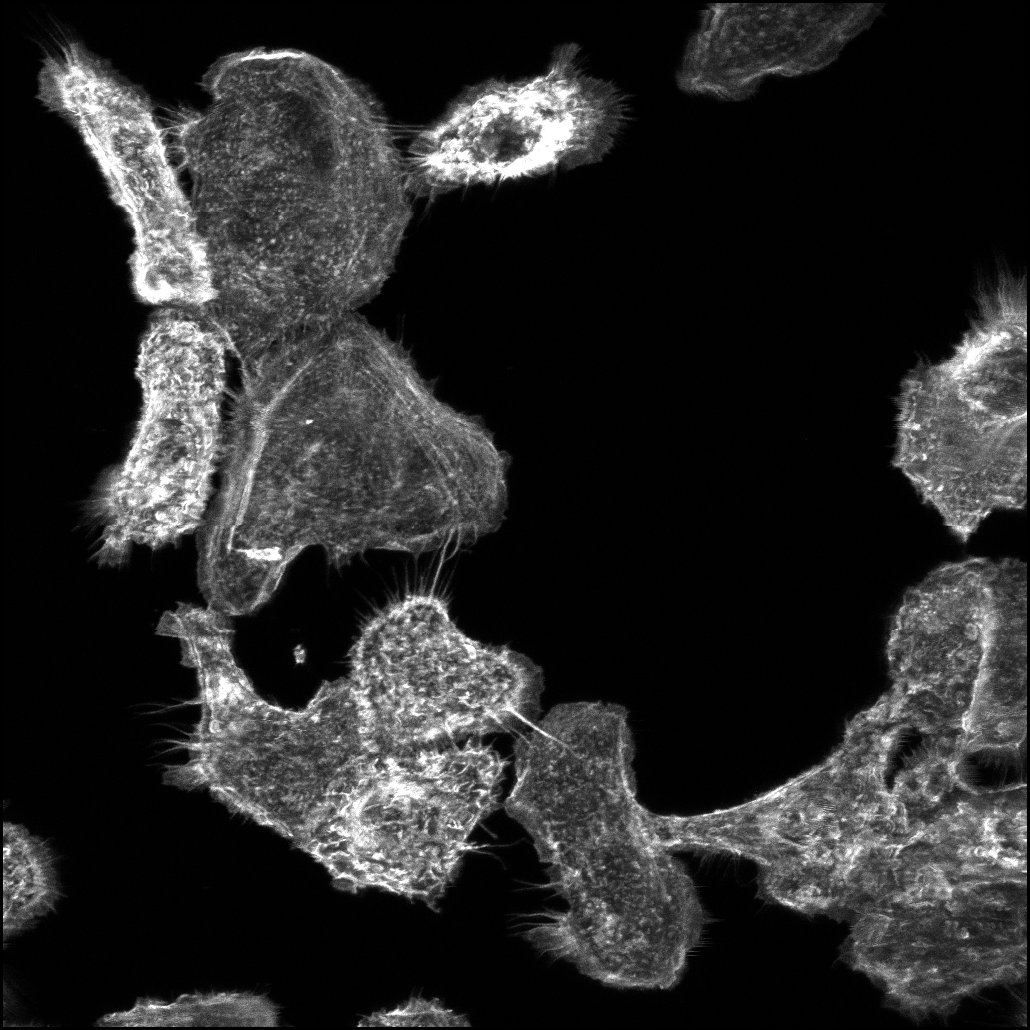

### 40x_1532C_DMSO_10UMKBU_2hrs_activated beta1(green)_actin(red)_kbu_04_DAPI-T4.tif

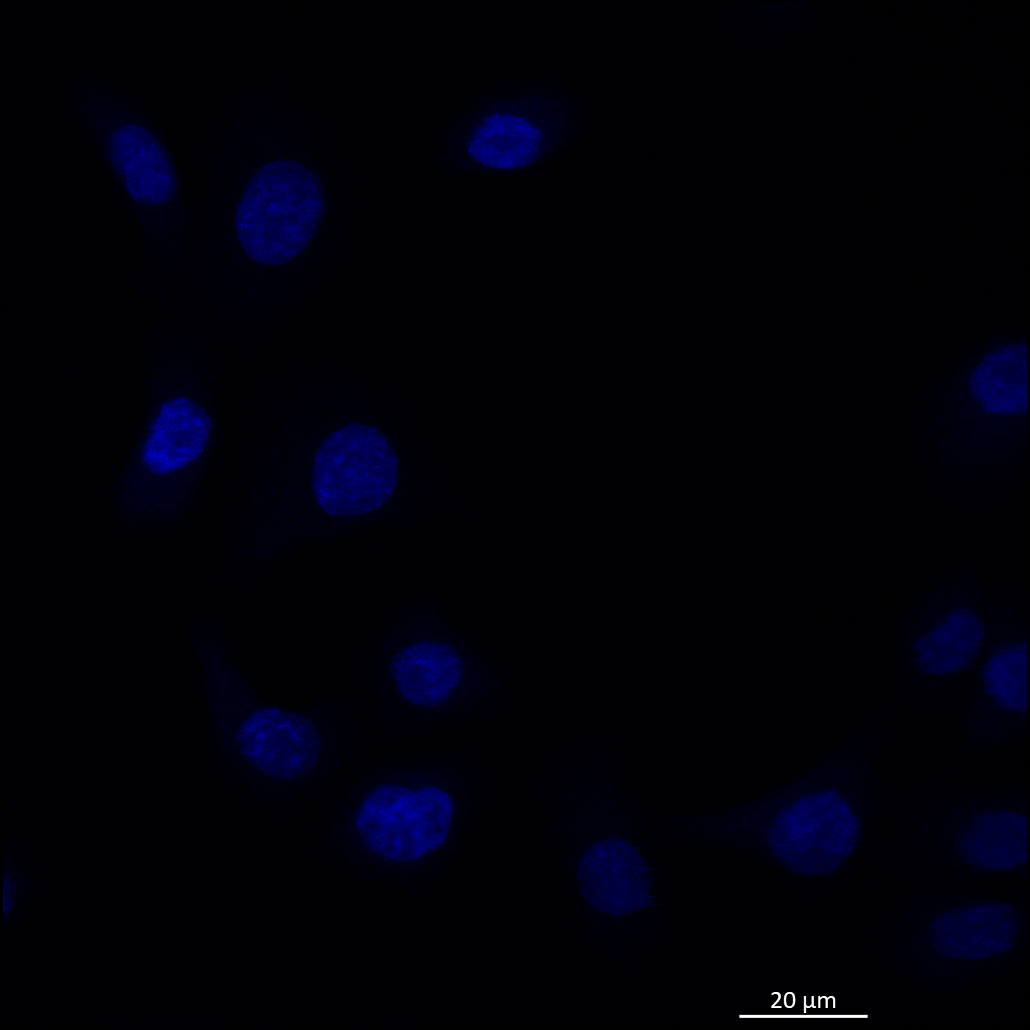

### 40x_1532C_DMSO_10UMKBU_2hrs_activated beta1(green)_actin(red)_kbu_04_DAPI-T4_ORG.tif

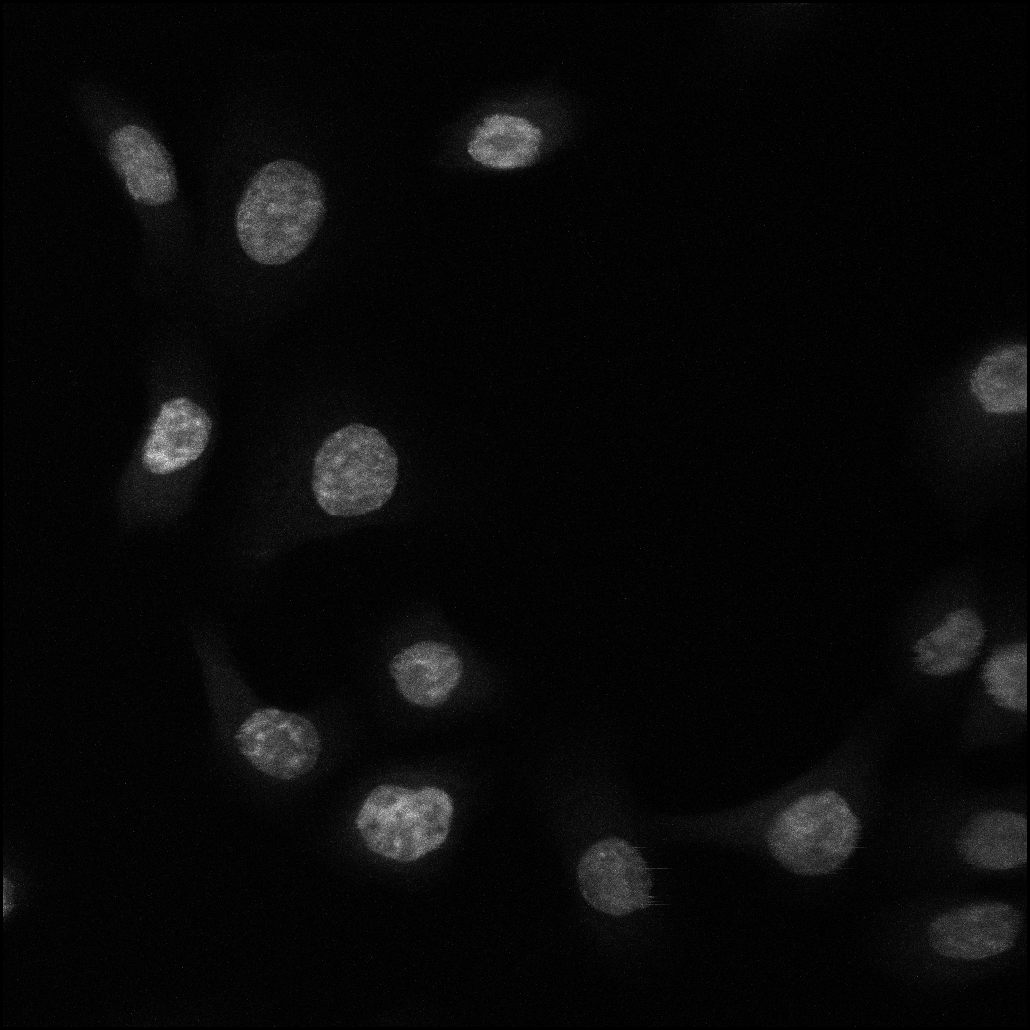

### Cutoff1_1uM_DiffTime_Comparison Analysis(visualize BH p-value)_expaned header.png

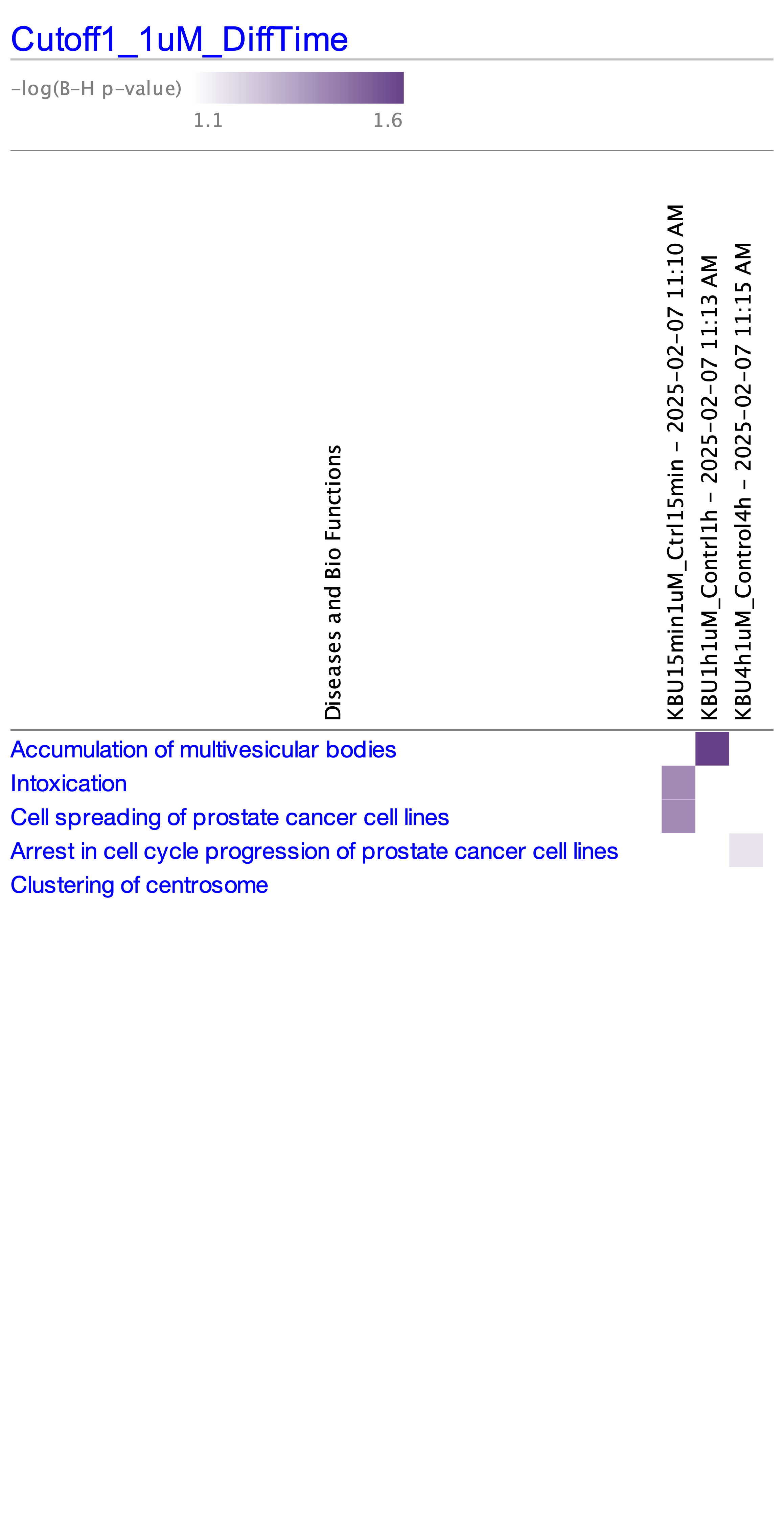

### Cutoff1_1uM_DiffTime_Comparison Analysis(visualize BH p-value)_not expaned header.png

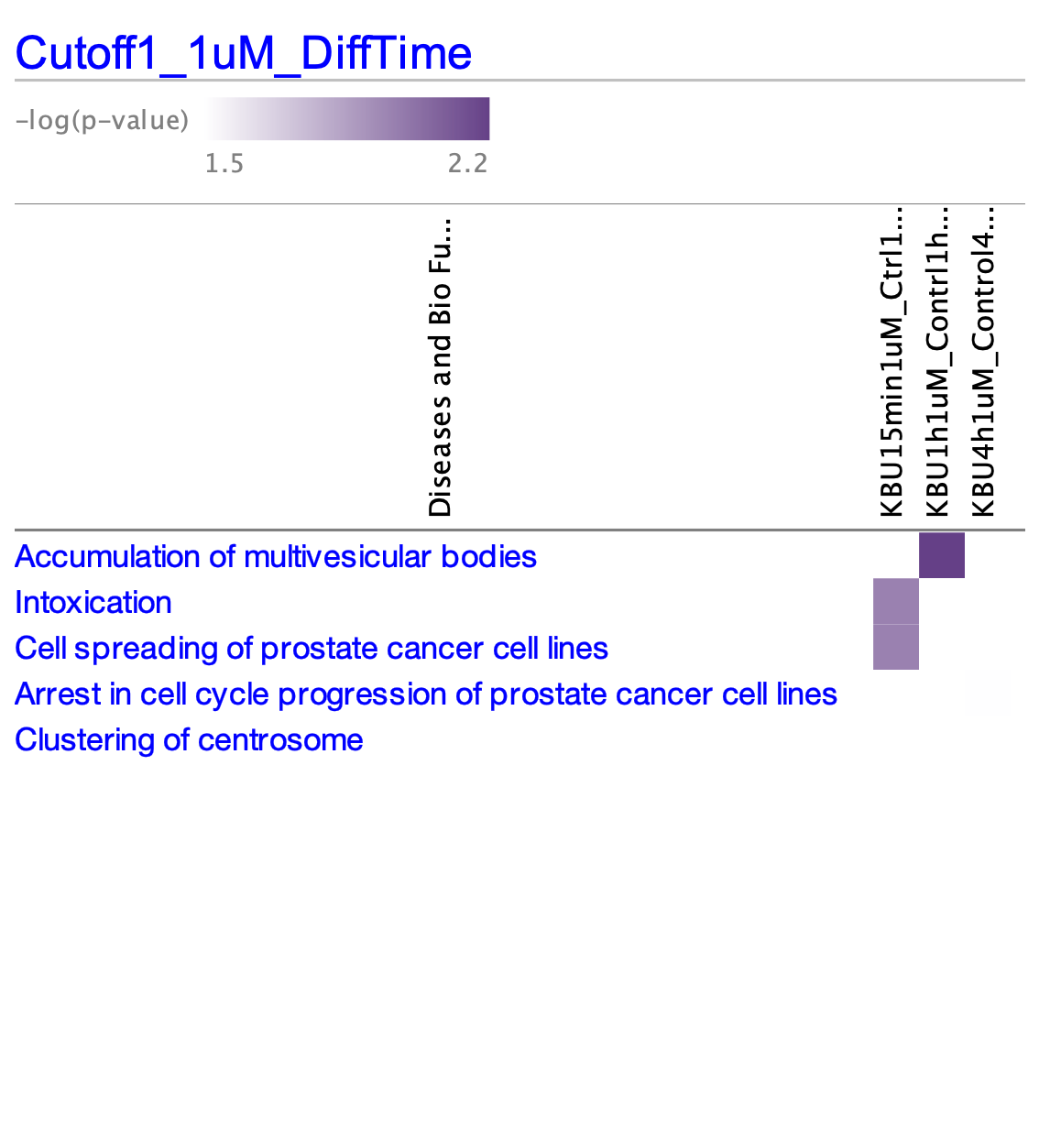

### Enrichr_results_bar_whole cell up.png

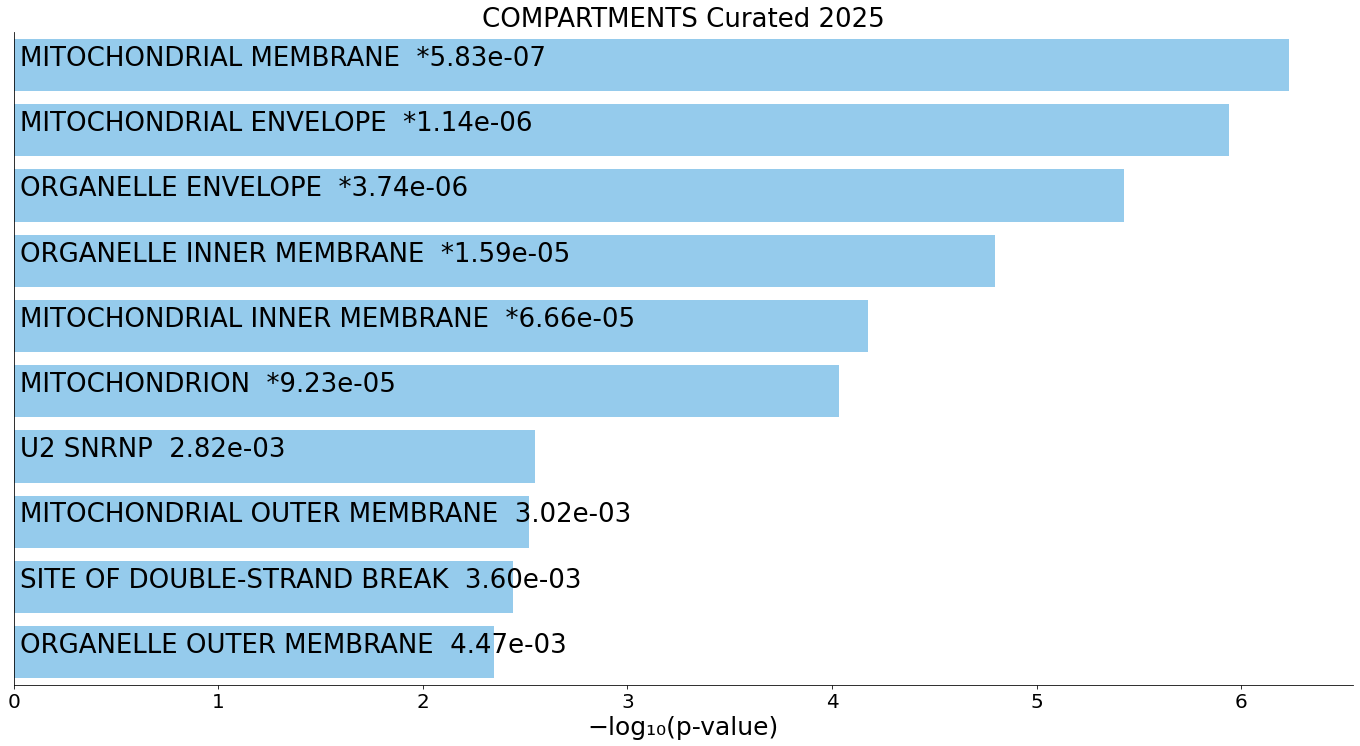

### Enrichr_results_bar_whole cell_down.png

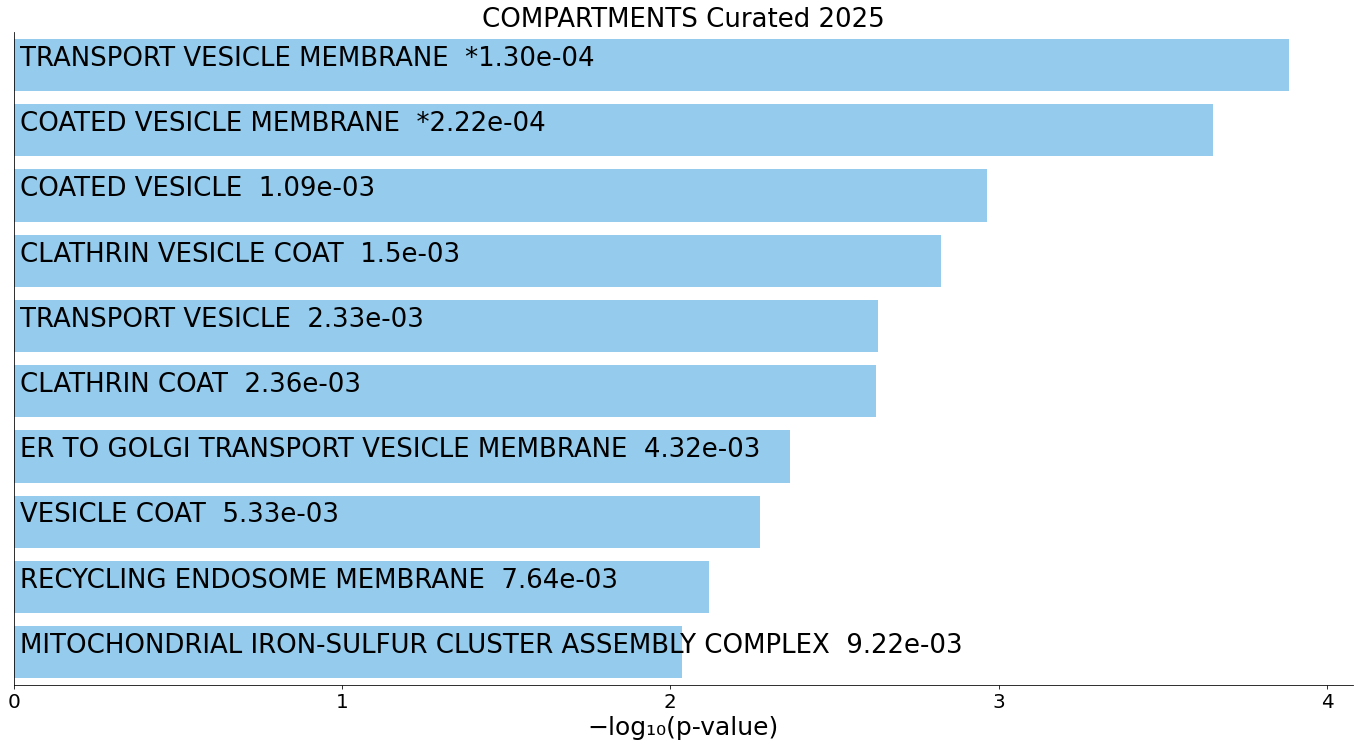

### Enrichr_results_bar_whole cell_down.png 1,360├ù746 pixels.pdf

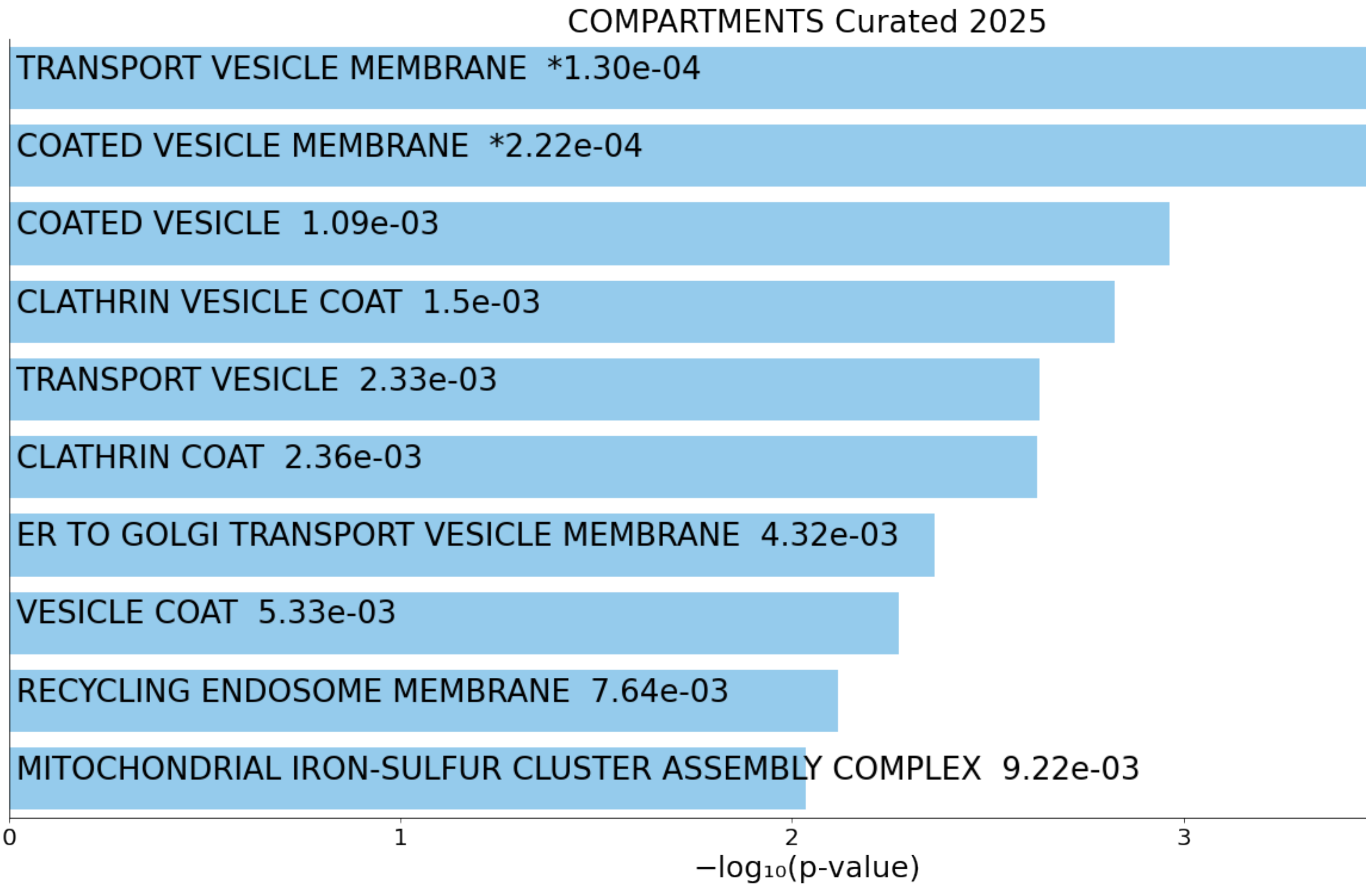

### Enrichr_results_bar_whole cell_up.png 1,360├ù746 pixels.pdf

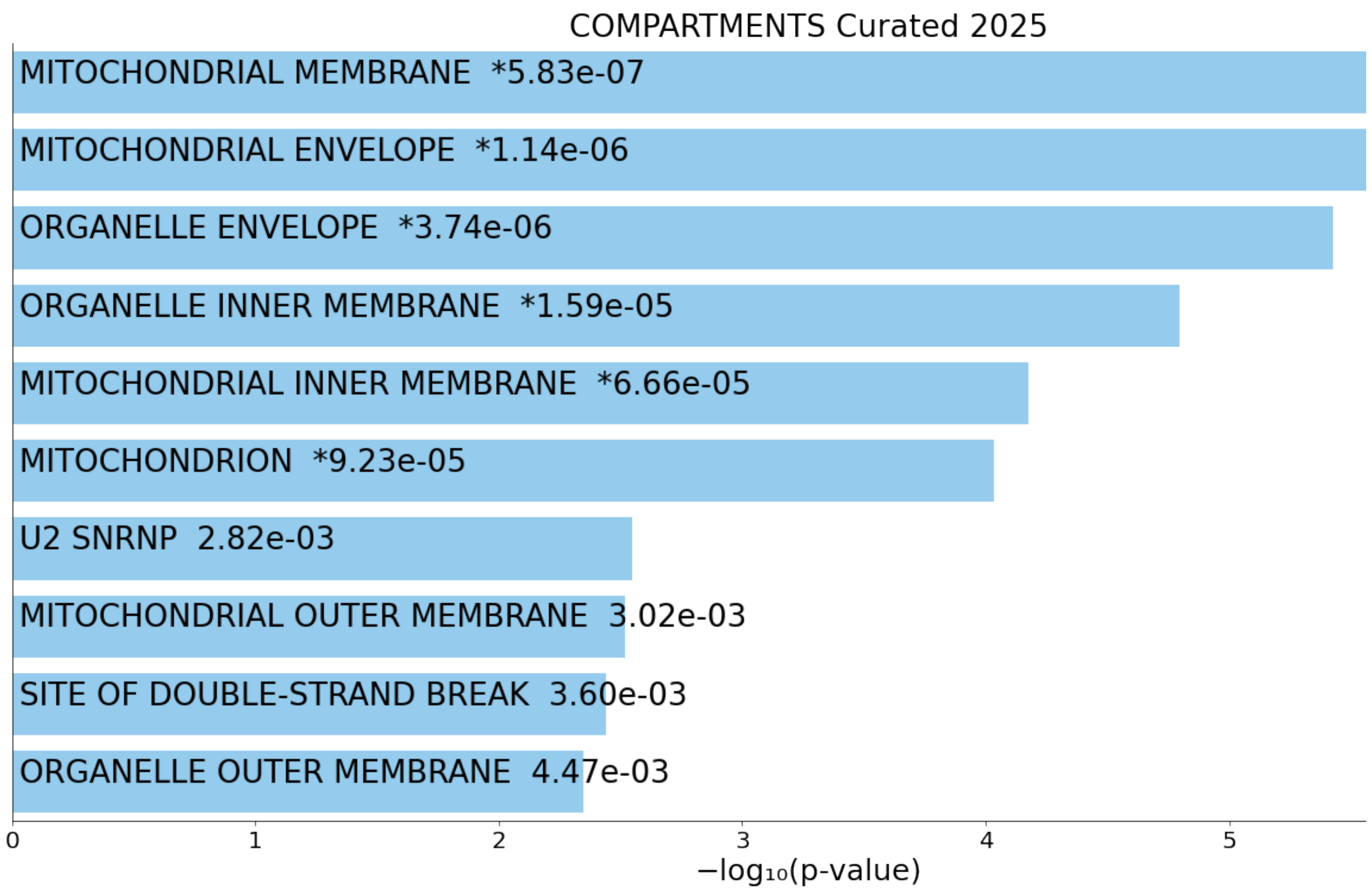

### Fig1_ResidueDistribution_Main_UniqueSites_Bar.pdf

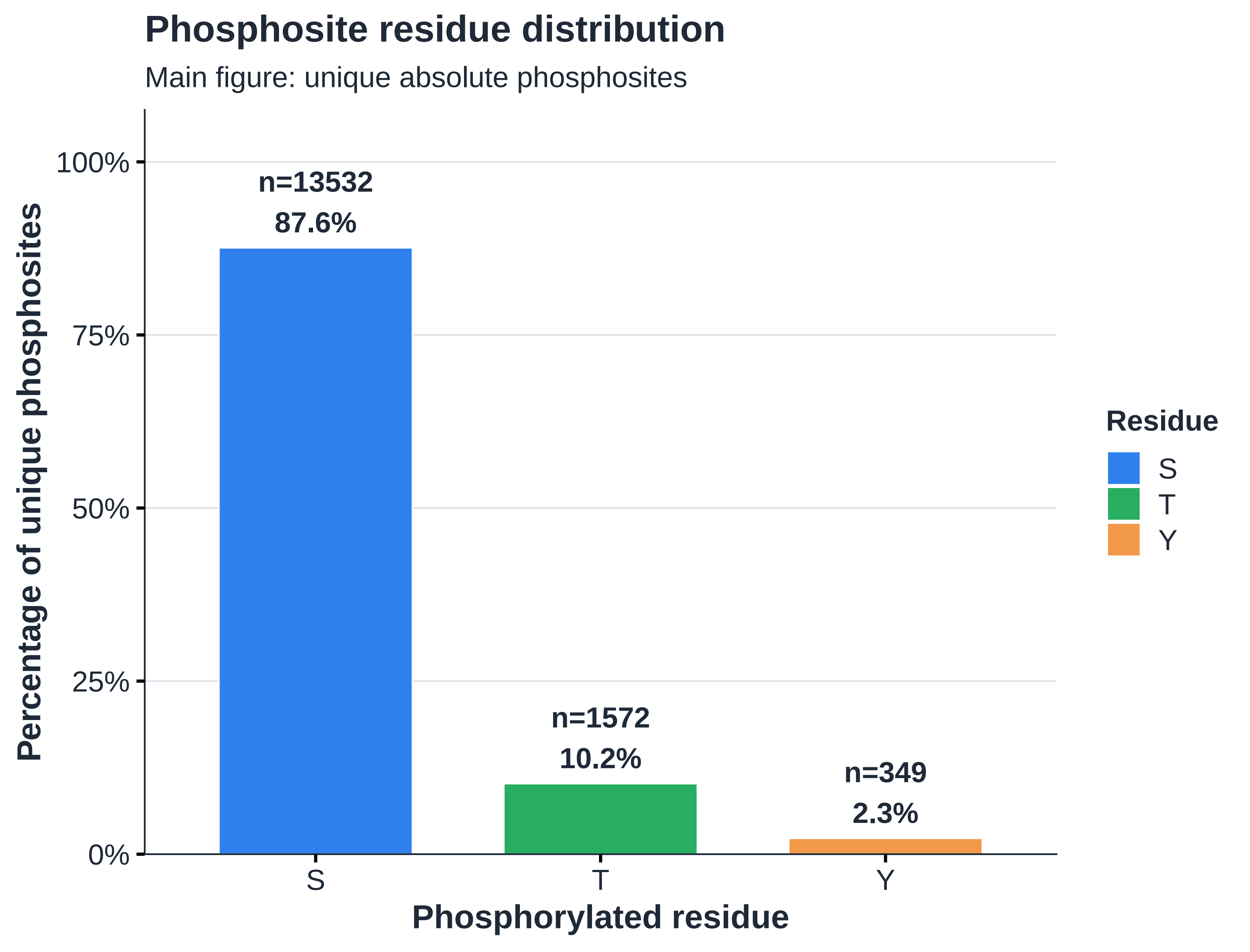

### Fig1_ResidueDistribution_Main_UniqueSites_Bar.png

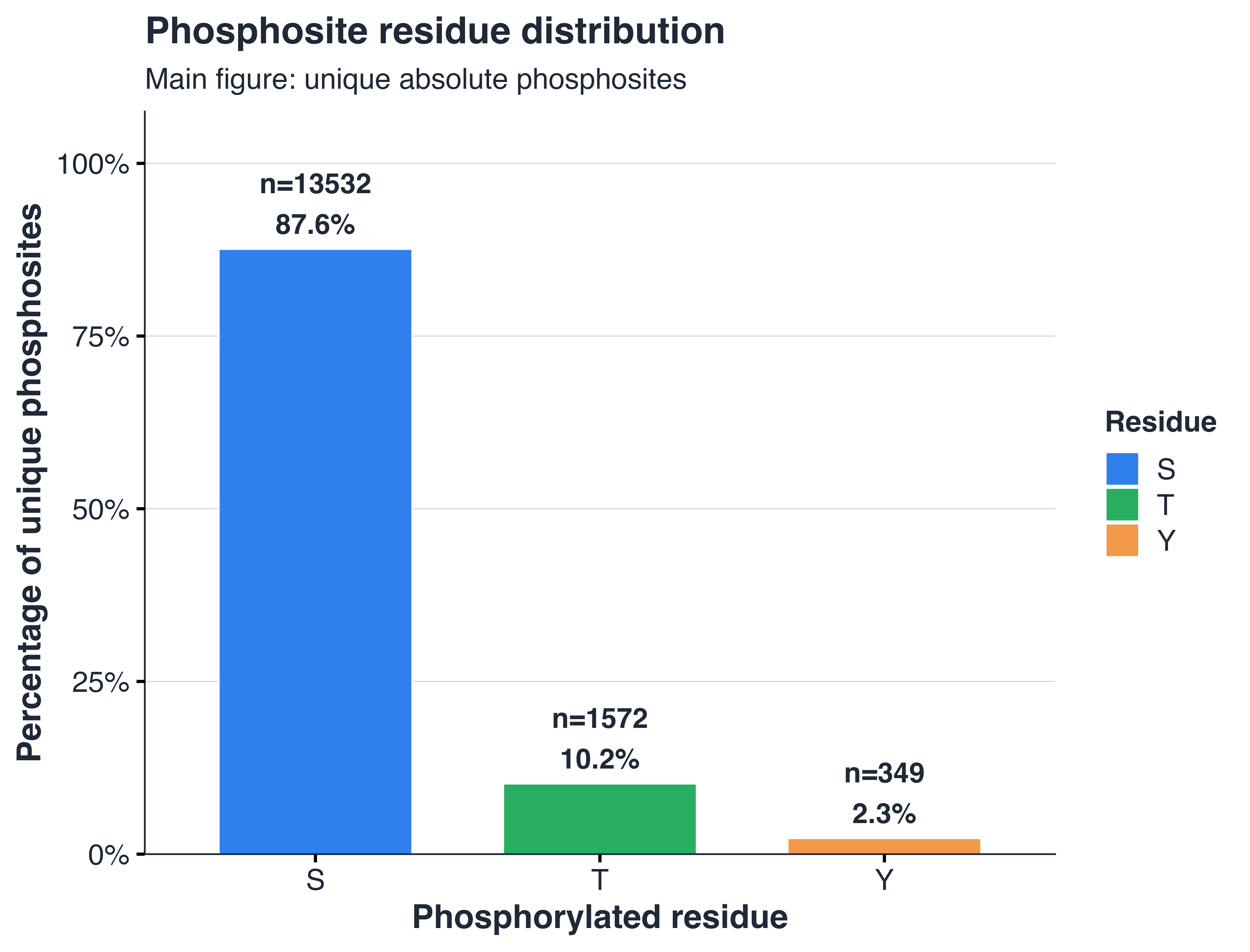

### Fig3_ResidueDistribution_Significant_UniqueSites_byTime.pdf

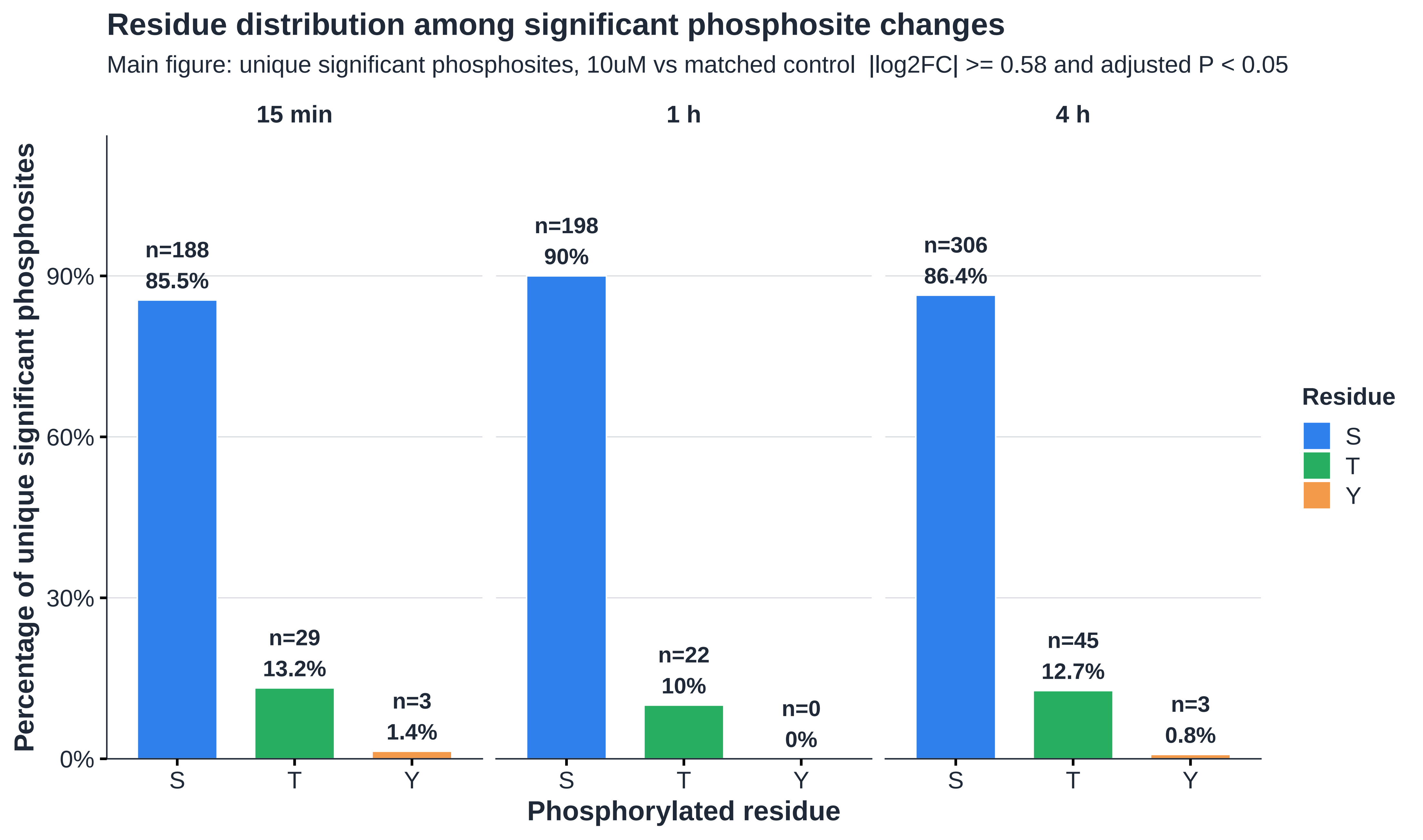

### Fig3_ResidueDistribution_Significant_UniqueSites_byTime.png

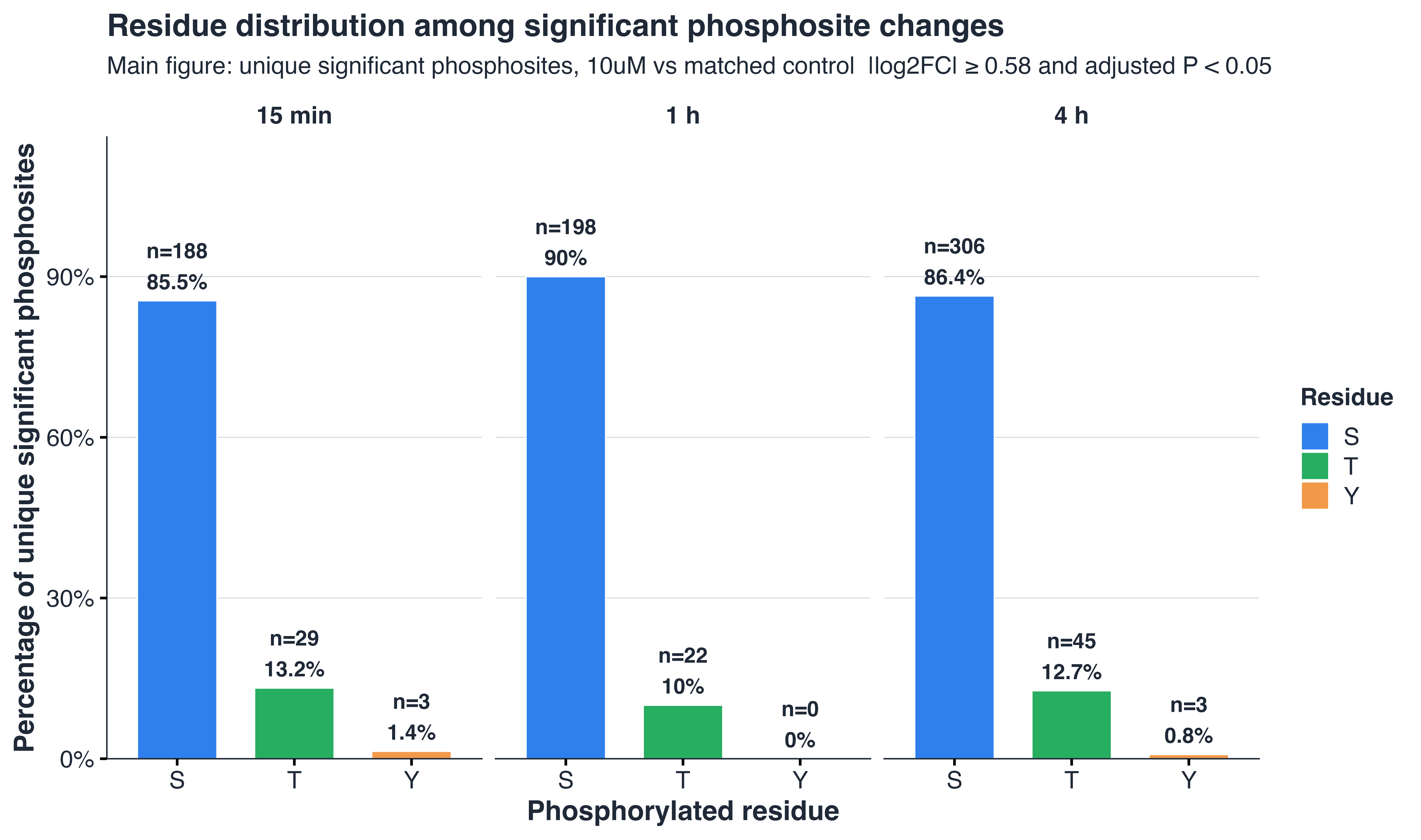

### Fig4_Venn_UniqueSignificantPhosphosites_10uM_vsCtrl.pdf

Unique significant phosphosites across time

10uM vs matched control; |log2FC| >= 0.58 and adjusted P < 0.05

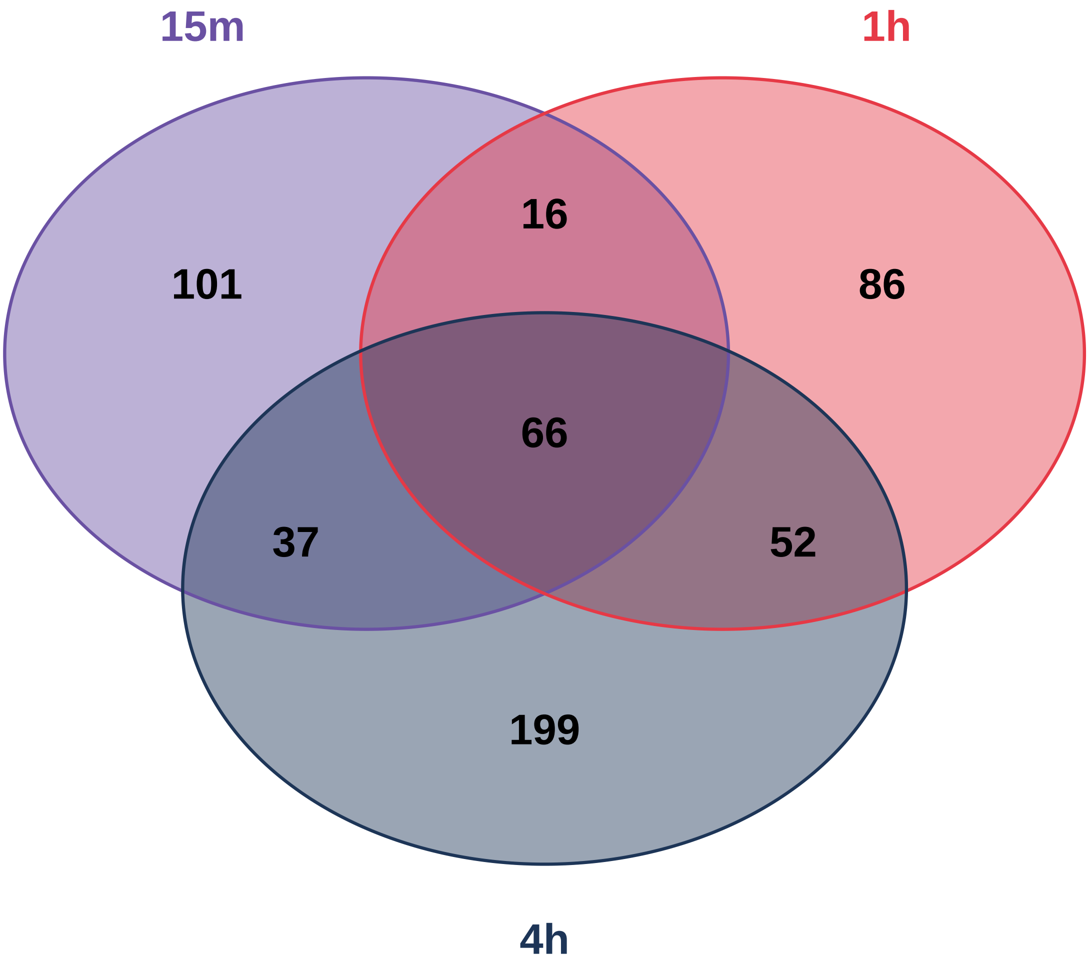

### Fig4_Venn_UniqueSignificantPhosphosites_10uM_vsCtrl.png

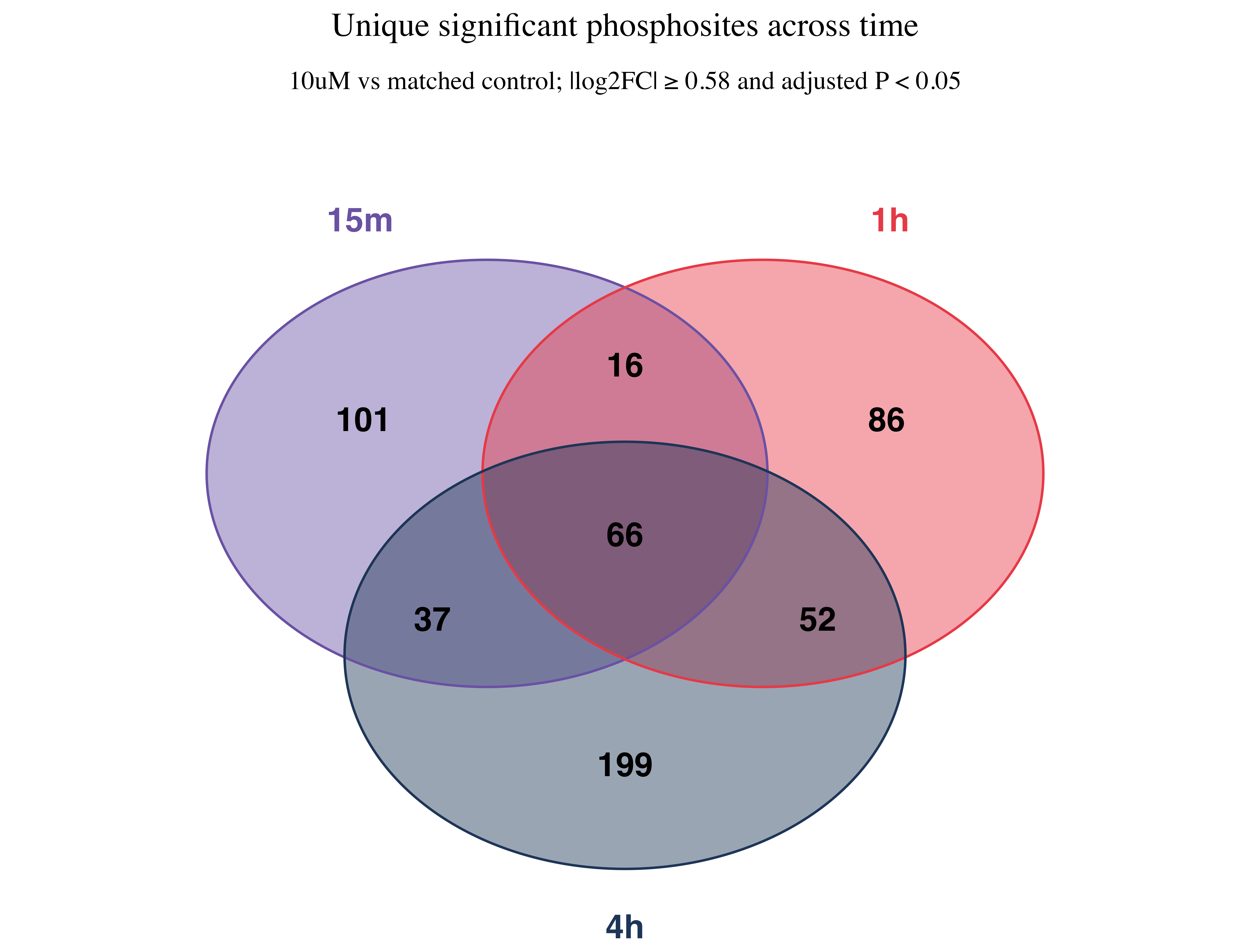

### Figure5D_MAPK14_Substrates_Heatmap_4h.pdf

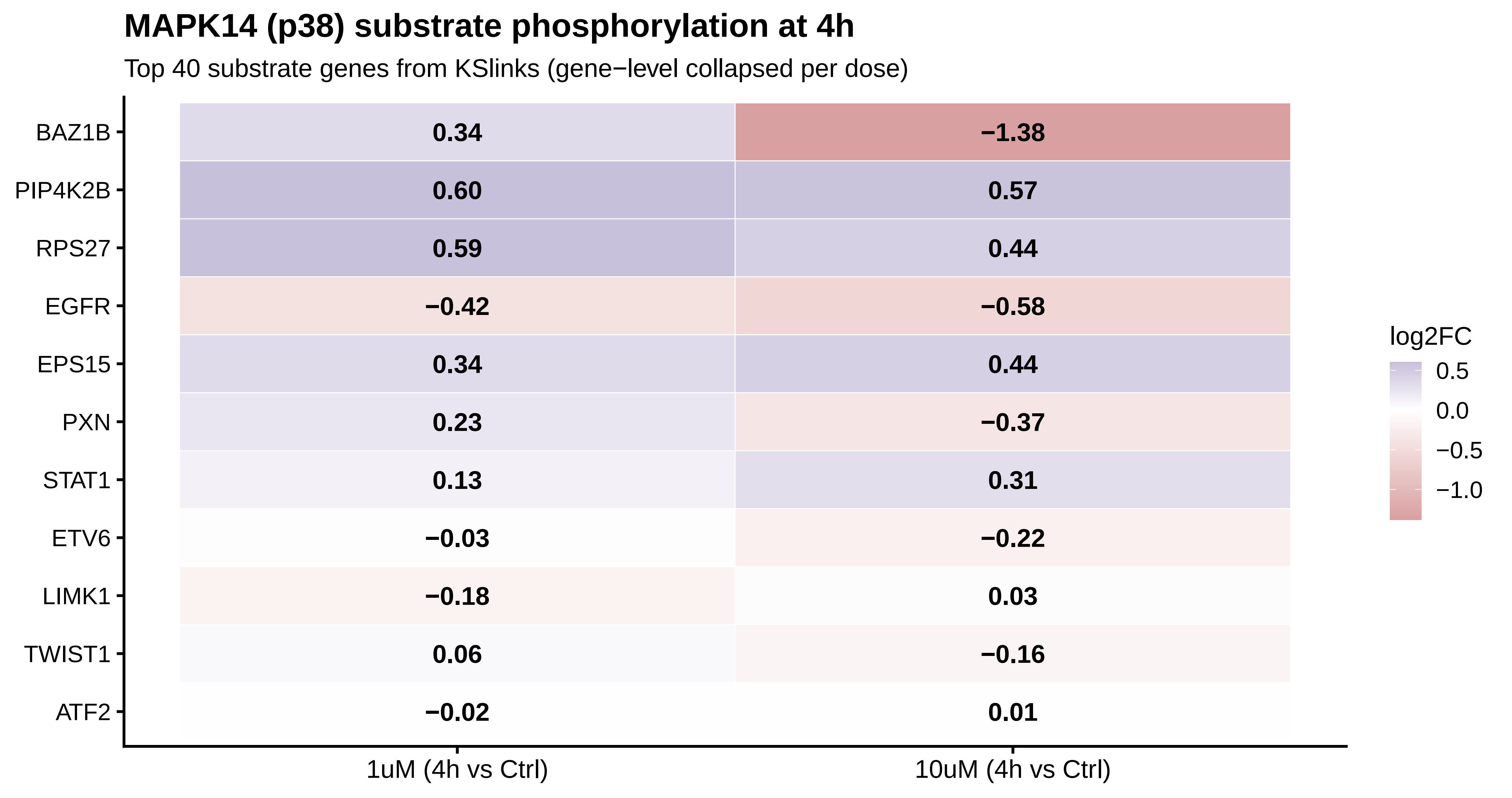

### Figure5D_MAPK14_Substrates_Heatmap_4h.png

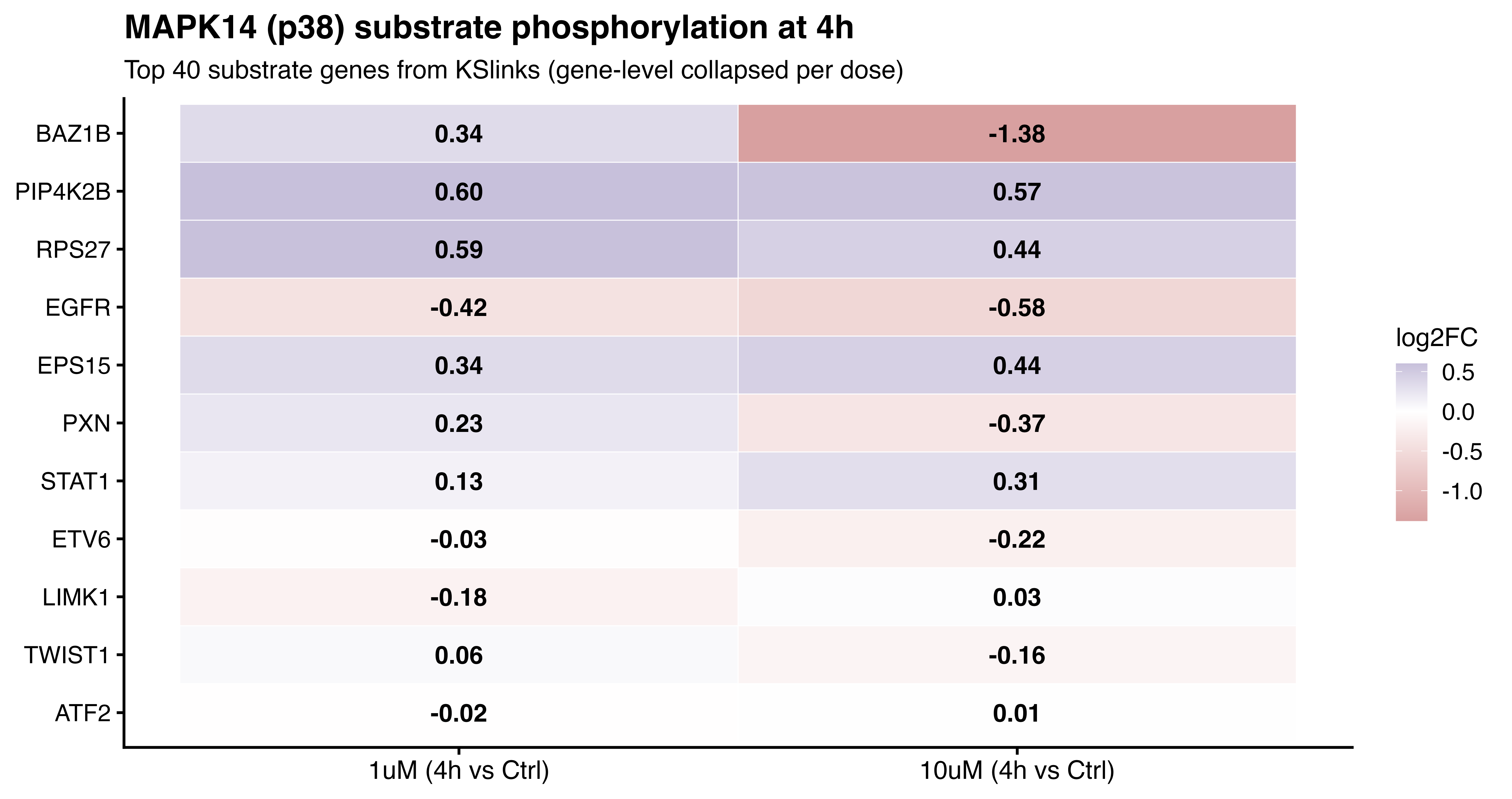

### KSEA_MAPK14_p38_Activity_4h.pdf

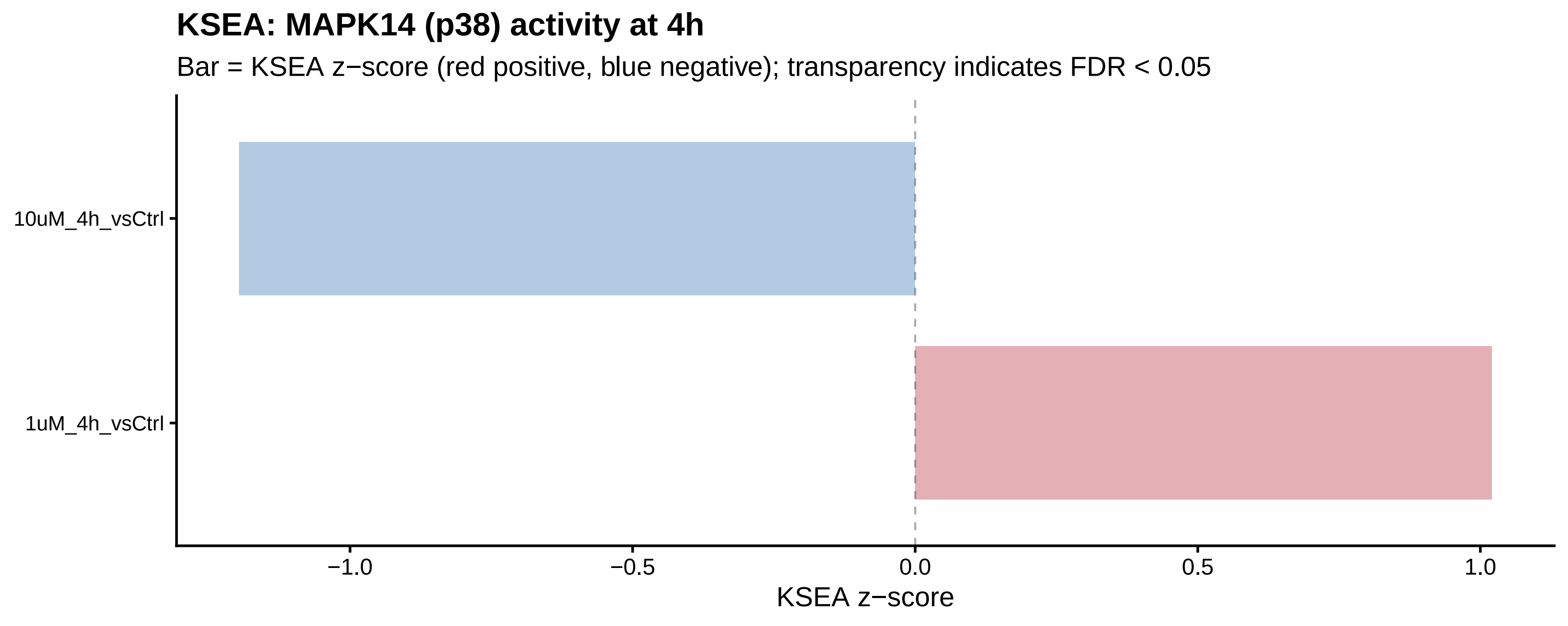

### KSEA_MAPK14_p38_Activity_4h.png

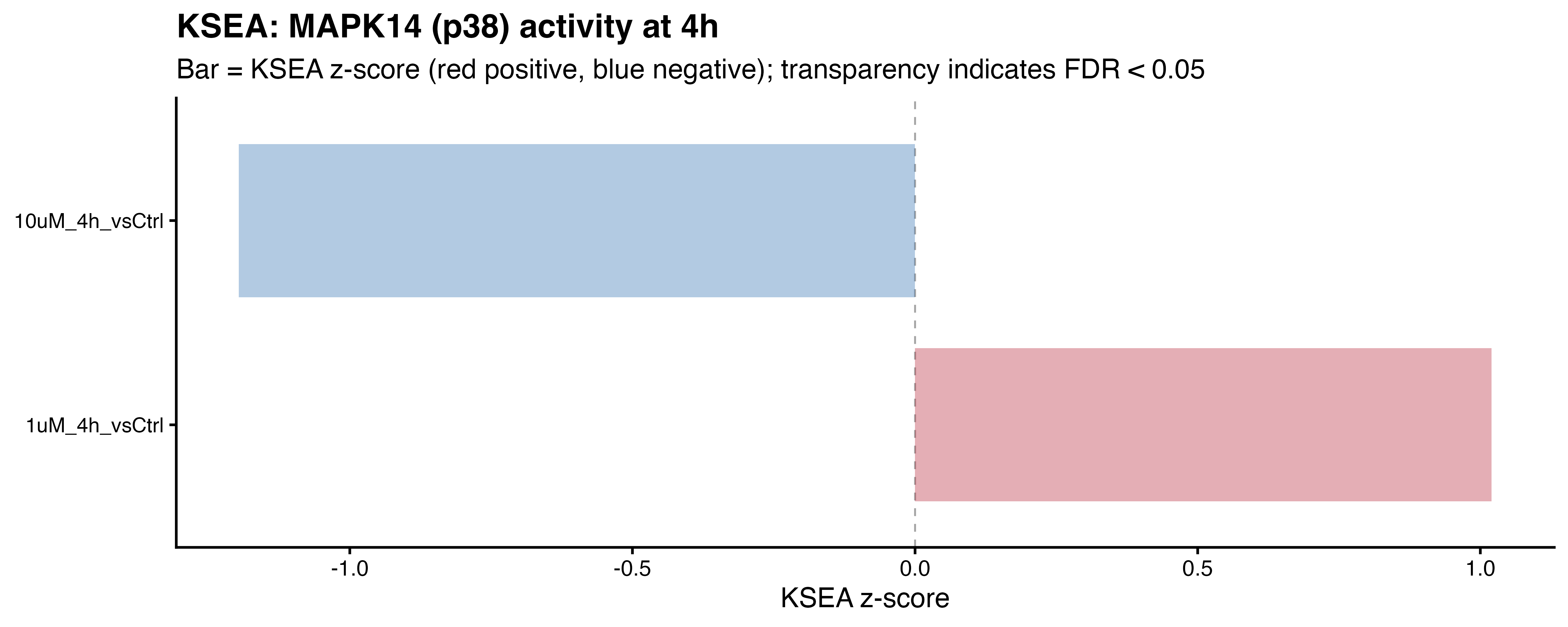

### MAPK14_p38_Sites_4h_Dumbbell.pdf

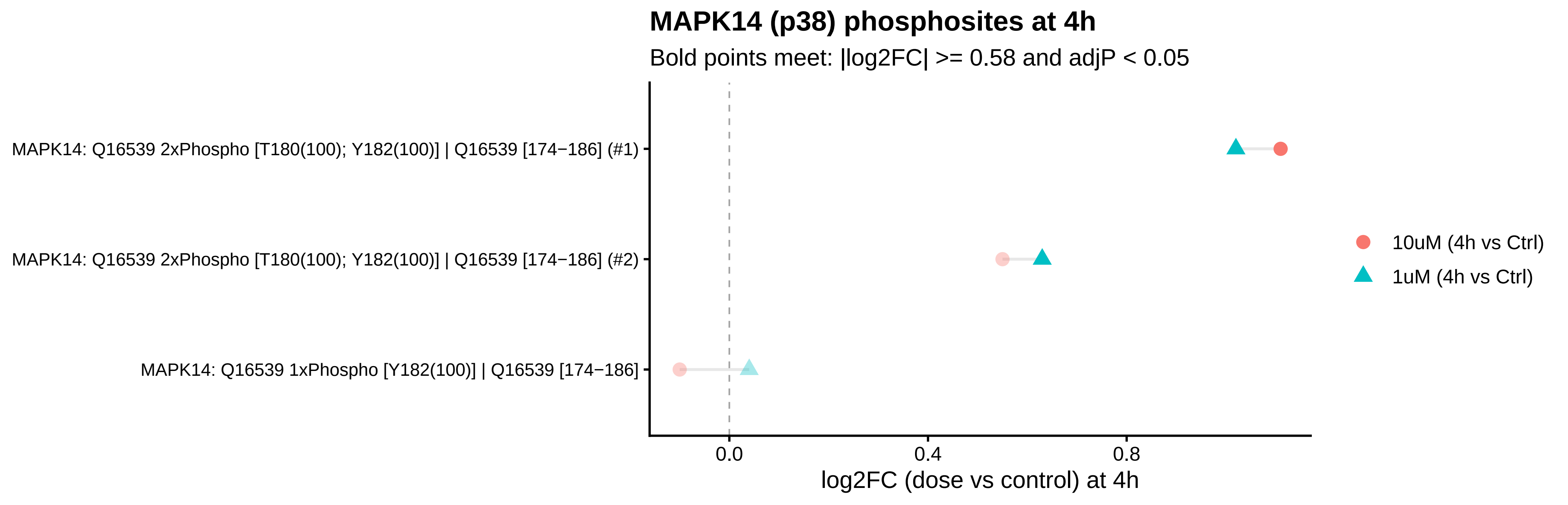

### MAPK14_p38_Sites_4h_Dumbbell.png

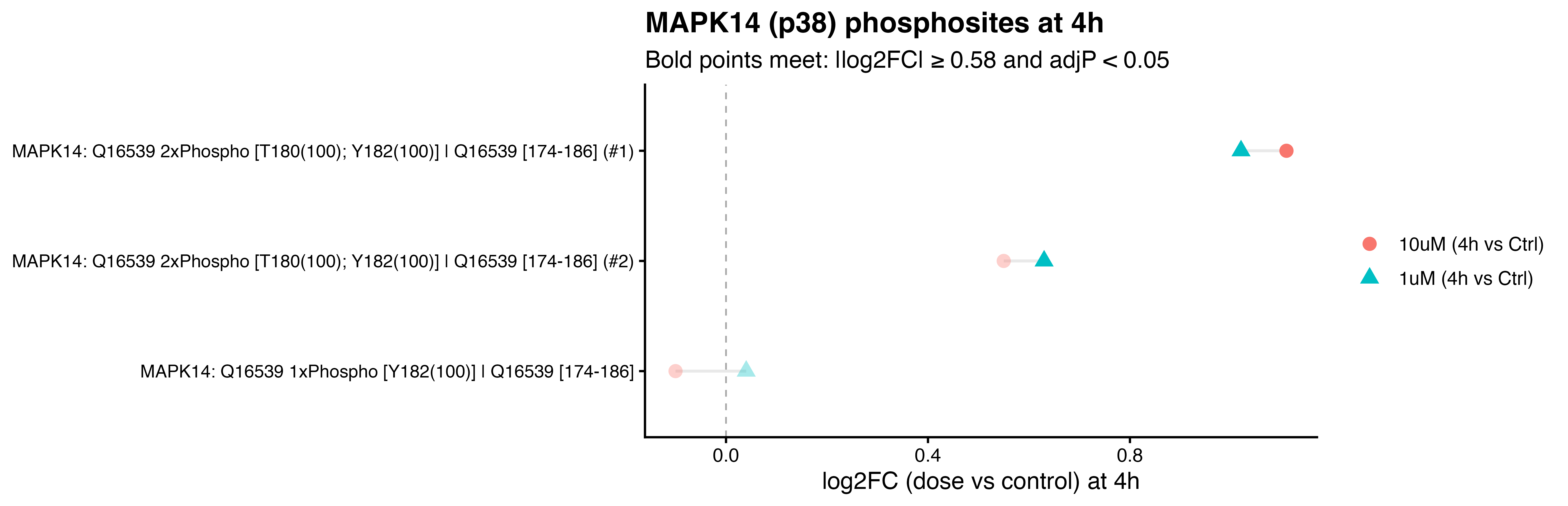

### Motility_Heatmap_log2FC_TopStable 3.55.15ΓÇ»PM.png

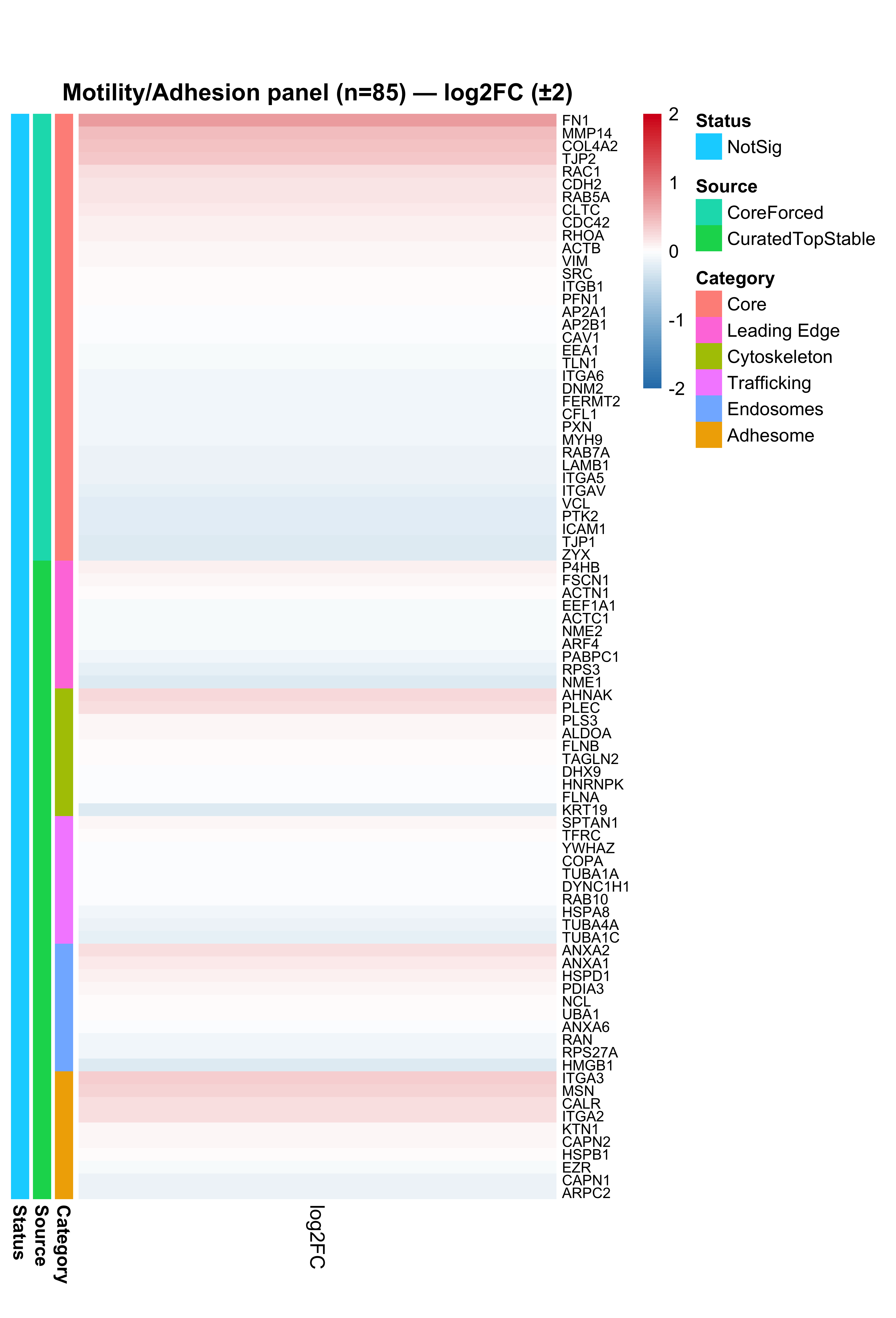

### Motility_Heatmap_log2FC_TopStable.pdf

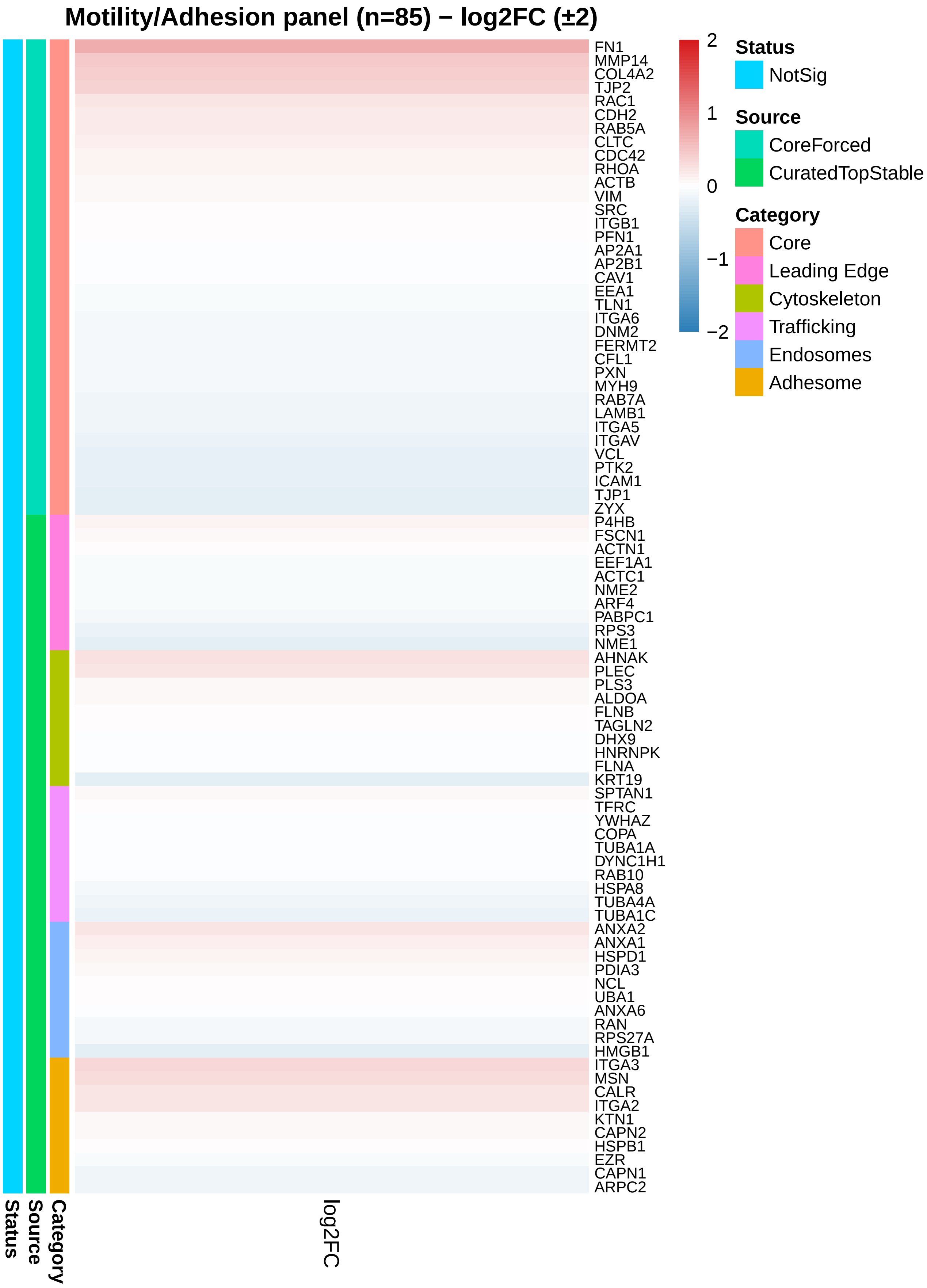

### PCA_PC1_vs_PC2.pdf

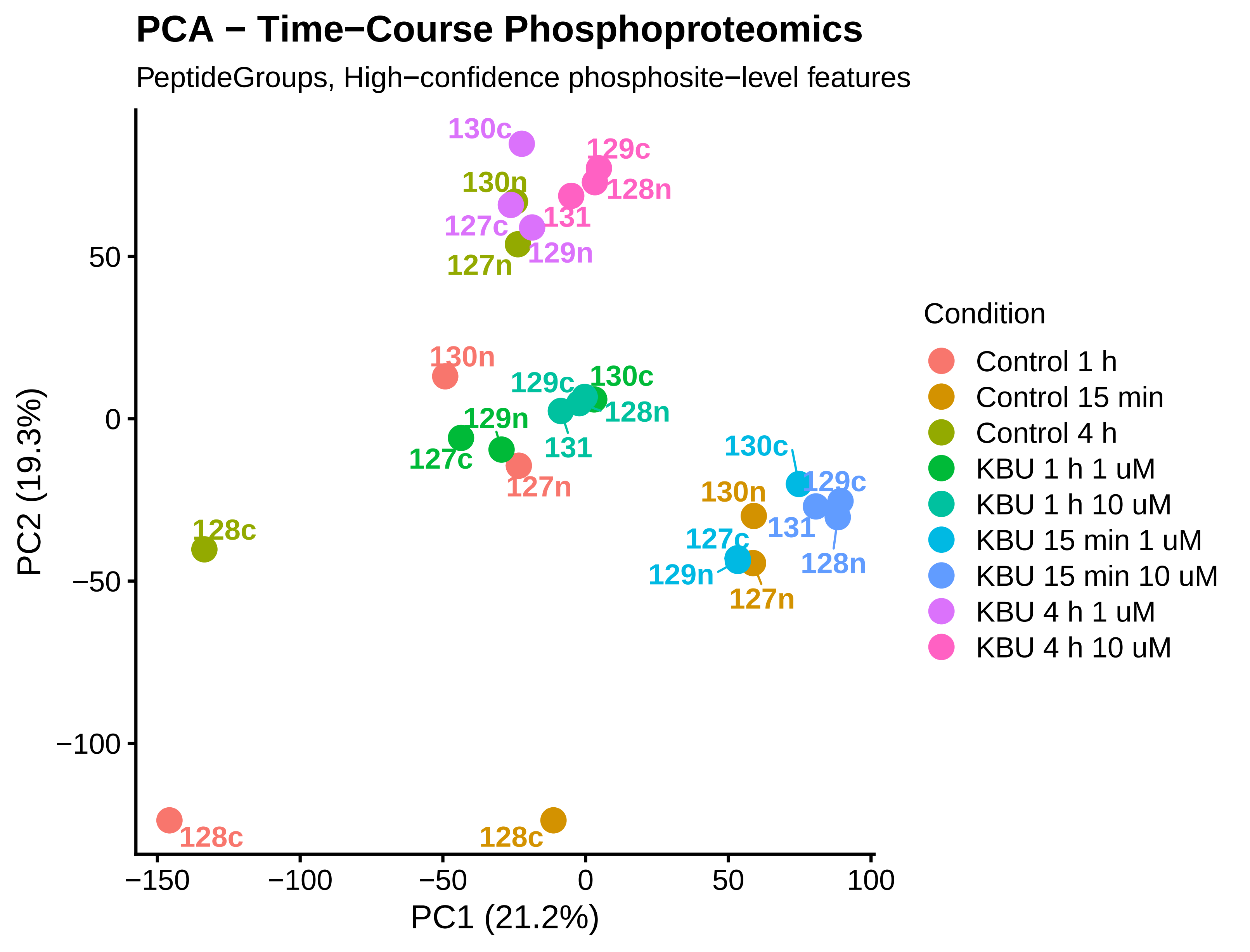

### PCA_PC1_vs_PC2.png

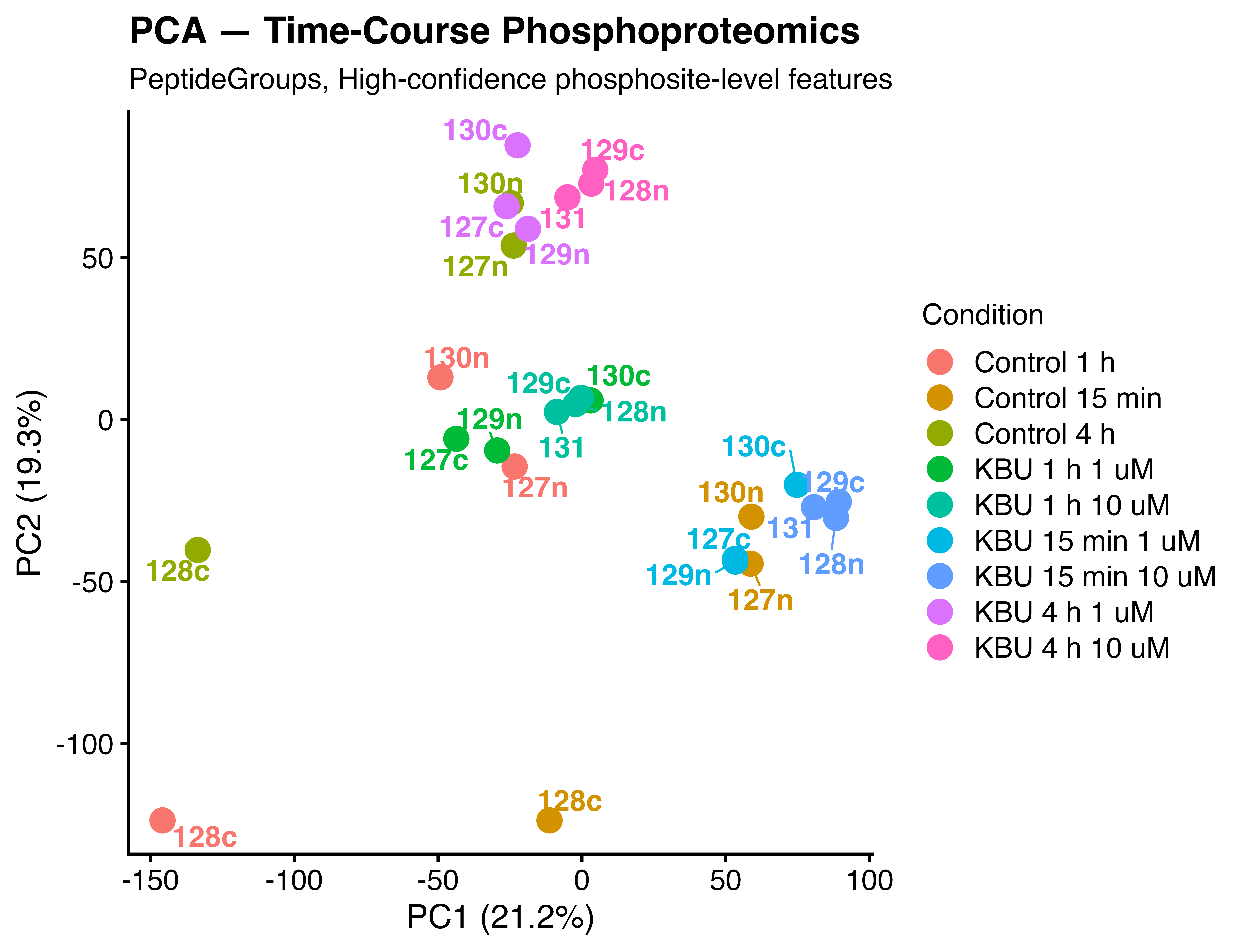

### PCA_PC1_vs_PC2_HQ.pdf

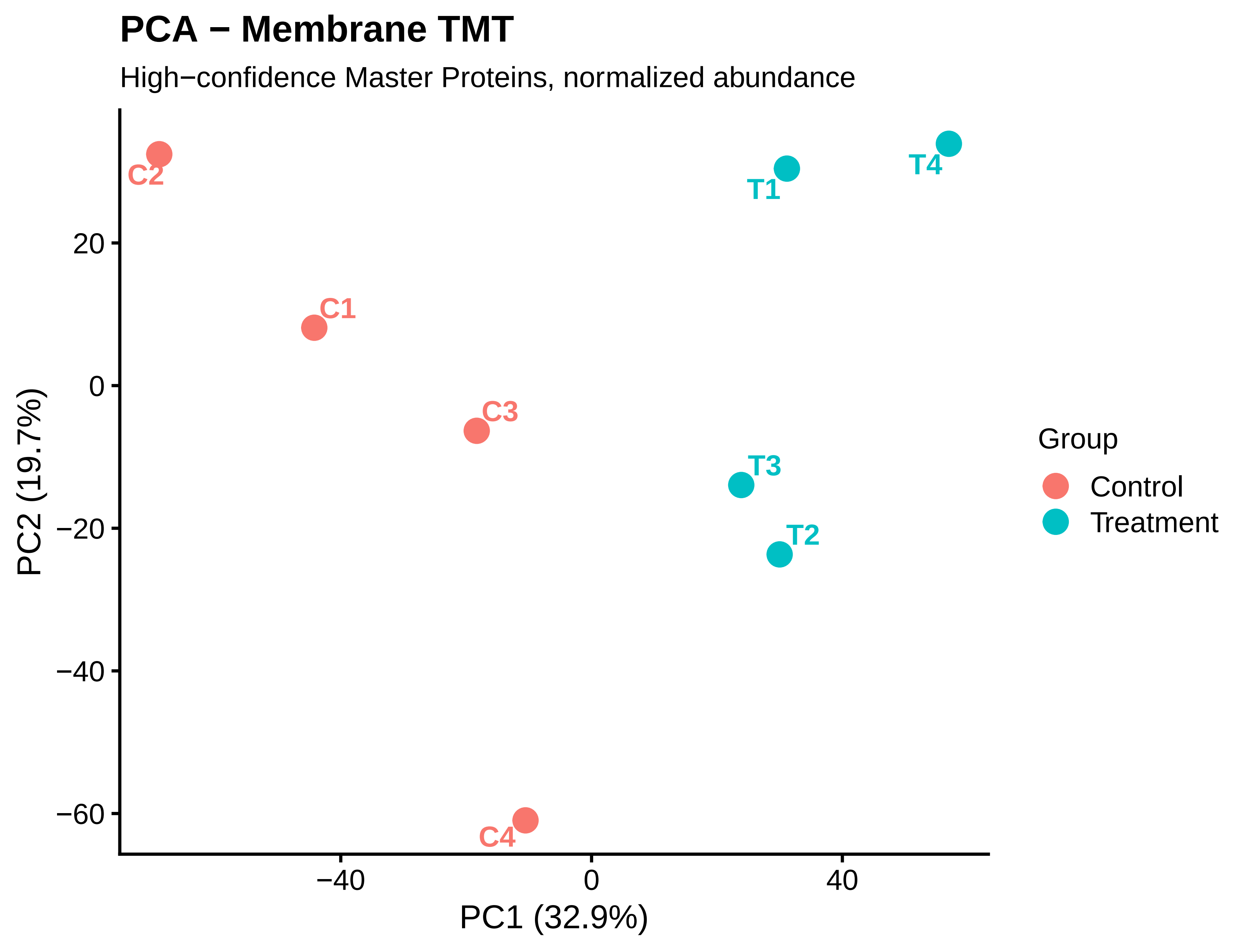

### PCA_PC1_vs_PC2_HQ.png

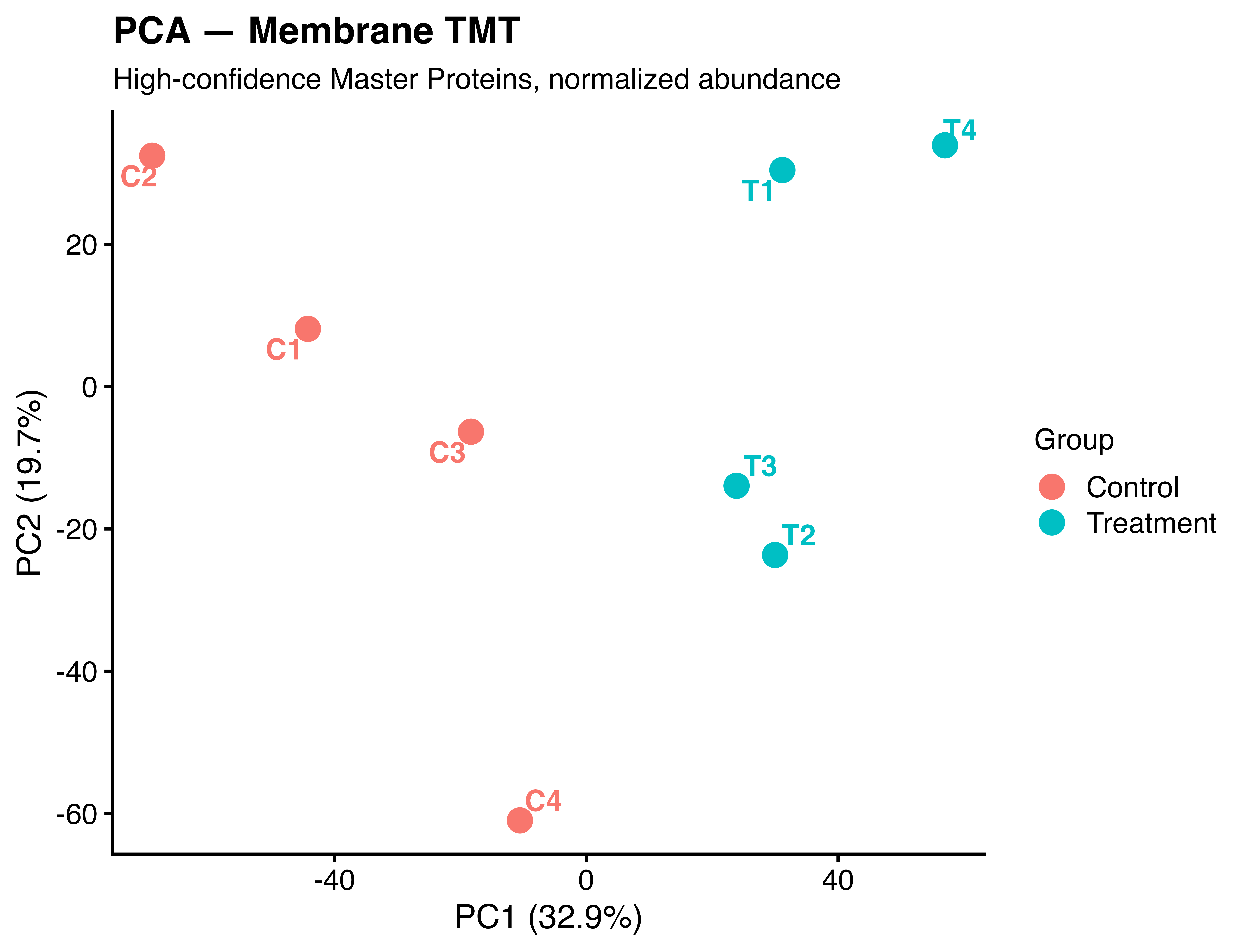

### QC_GO_CC_top100_dotplot_HQ.pdf

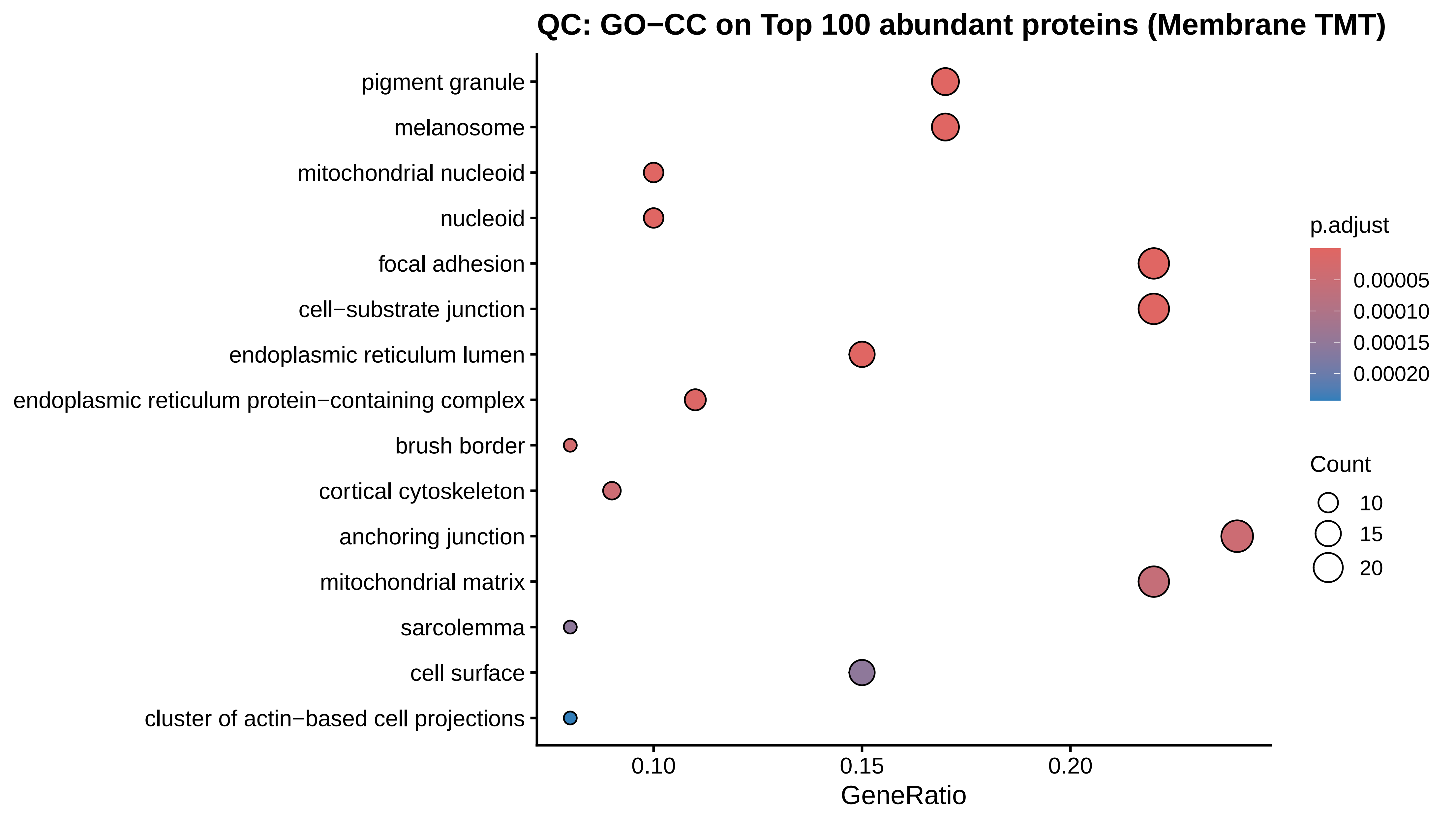

### QC_GO_CC_top100_dotplot_HQ.png

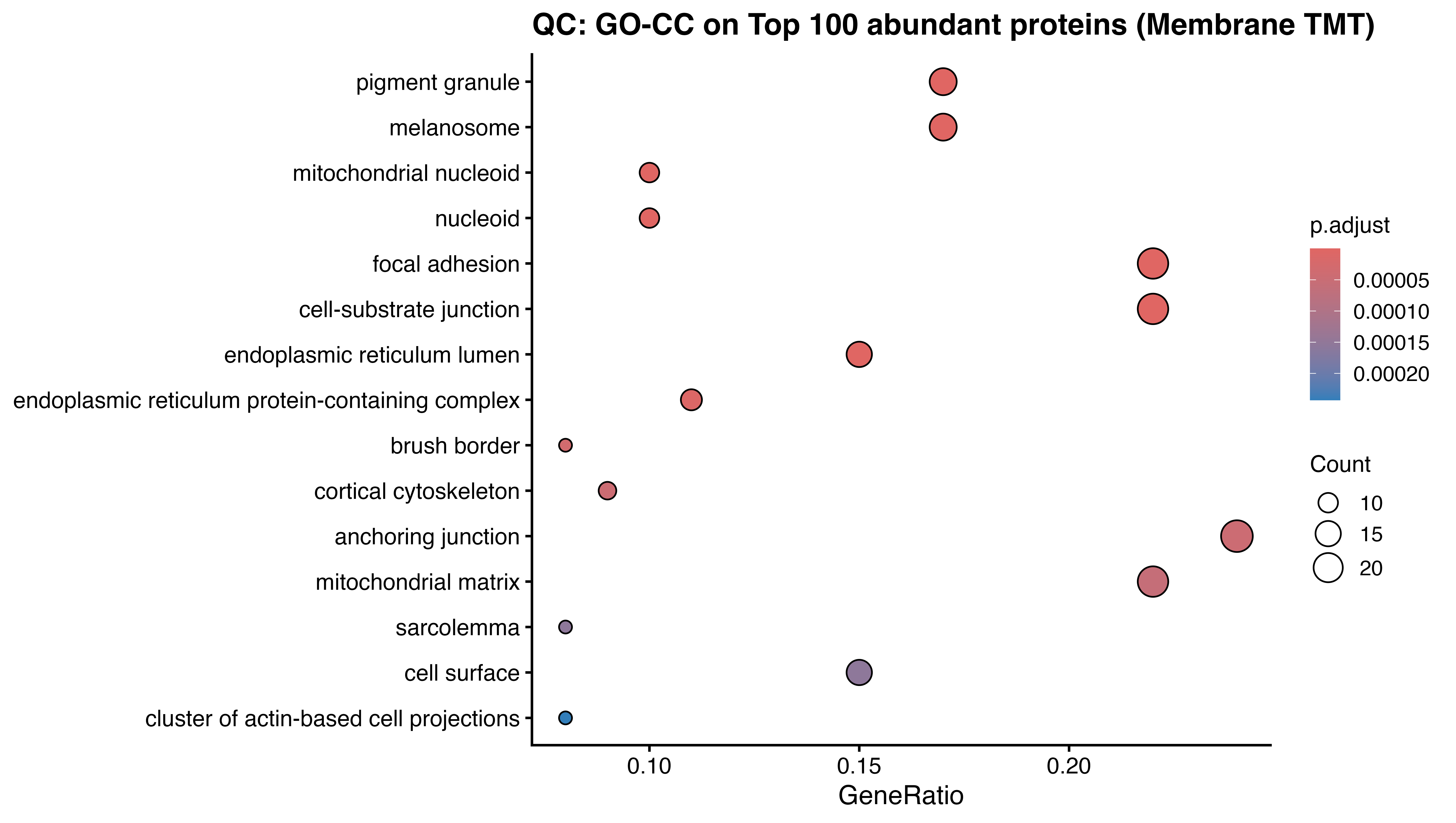

### QC_PCA_PC3_72h_HQ.pdf

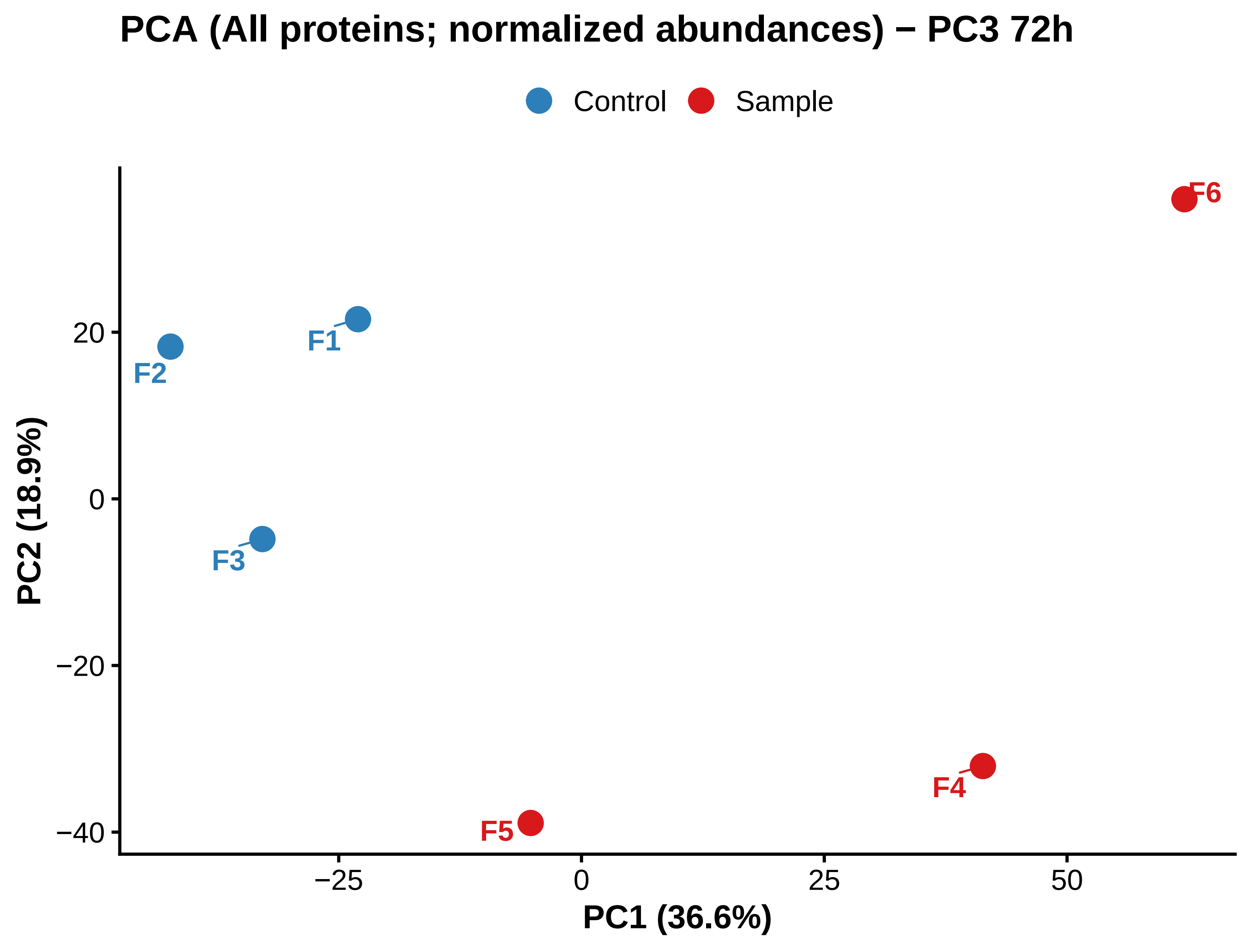

### QC_PCA_PC3_72h_HQ.png

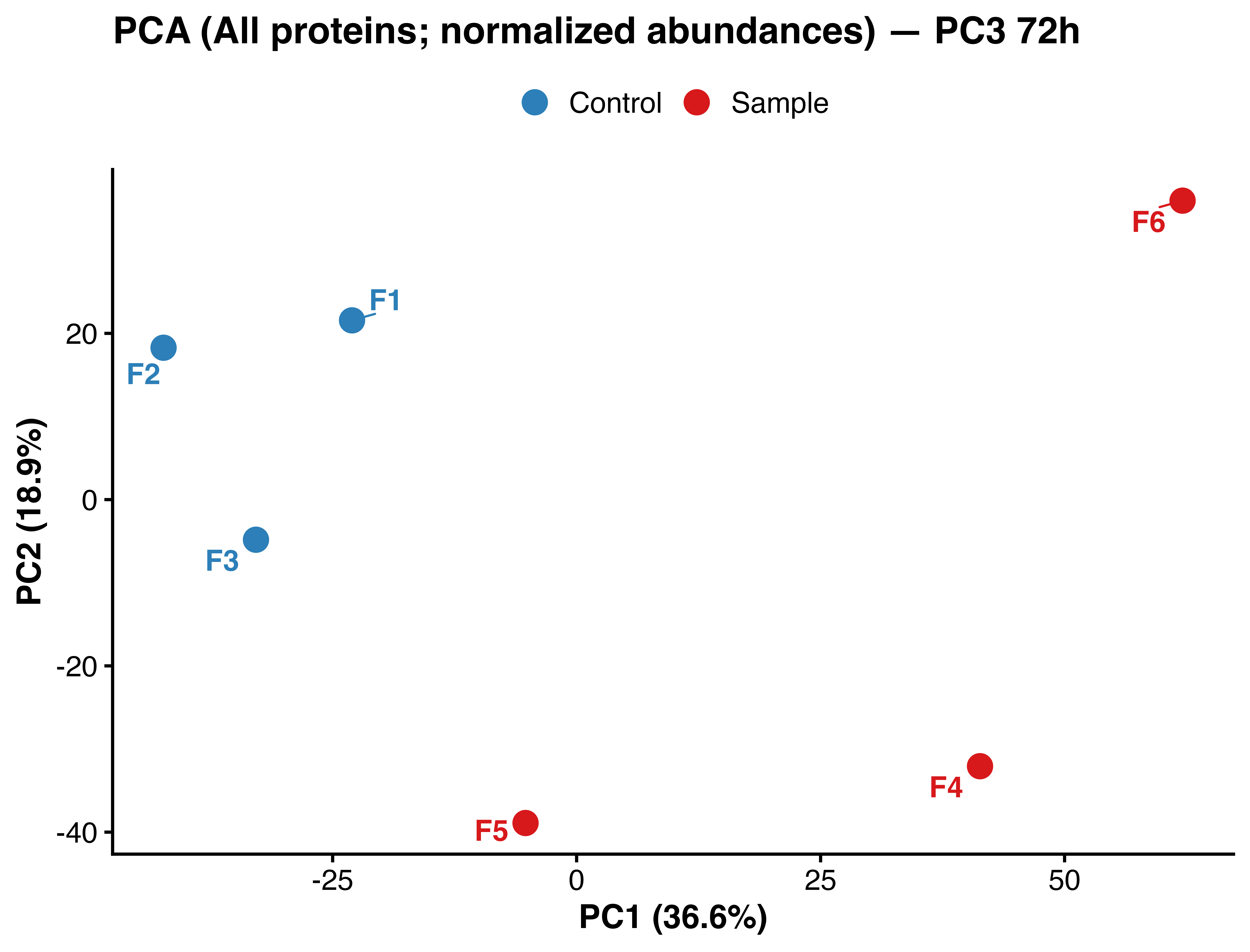

### Untitled8640x_1532C_DMSO_10UMKBU_2hrs_activated beta1(green)_actin(red)_DMSO_02_c1-4.tif

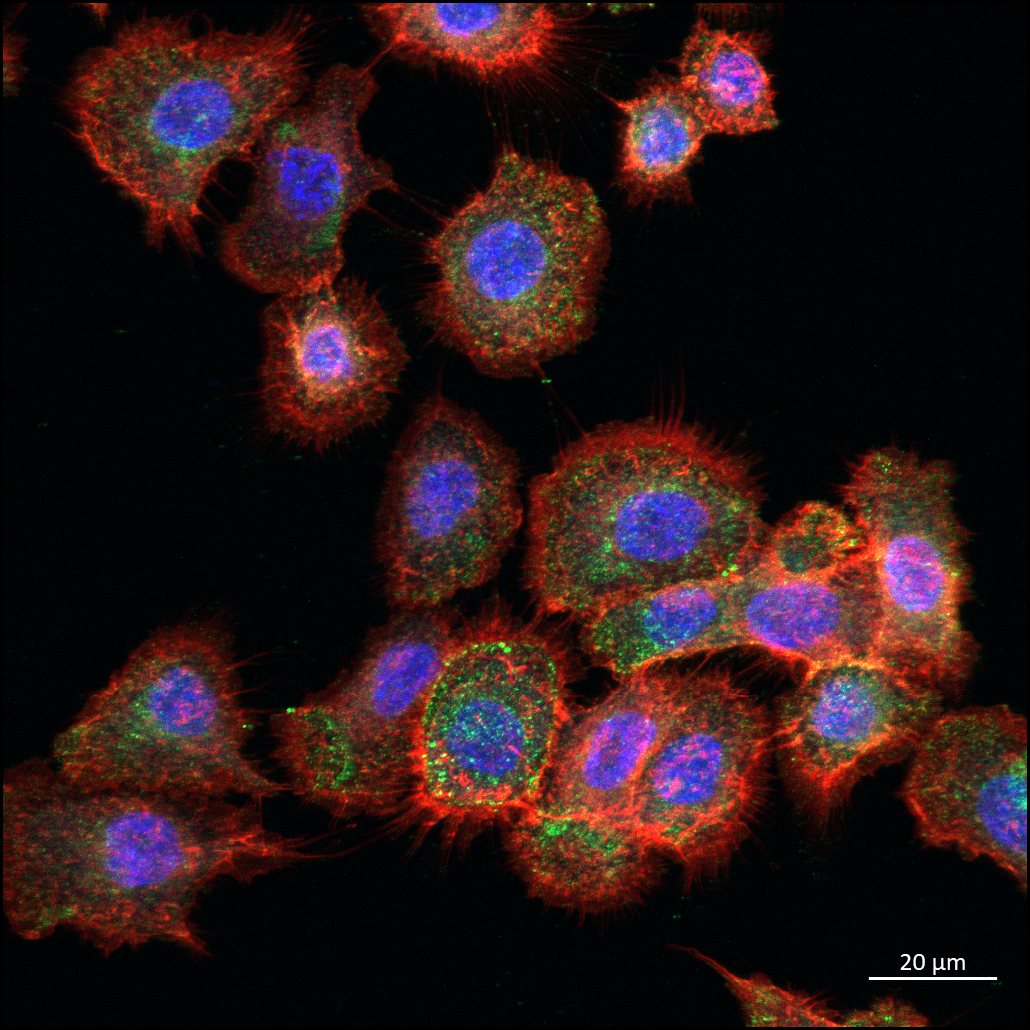

### Untitled8640x_1532C_DMSO_10UMKBU_2hrs_activated beta1(green)_actin(red)_DMSO_02_c1.tif

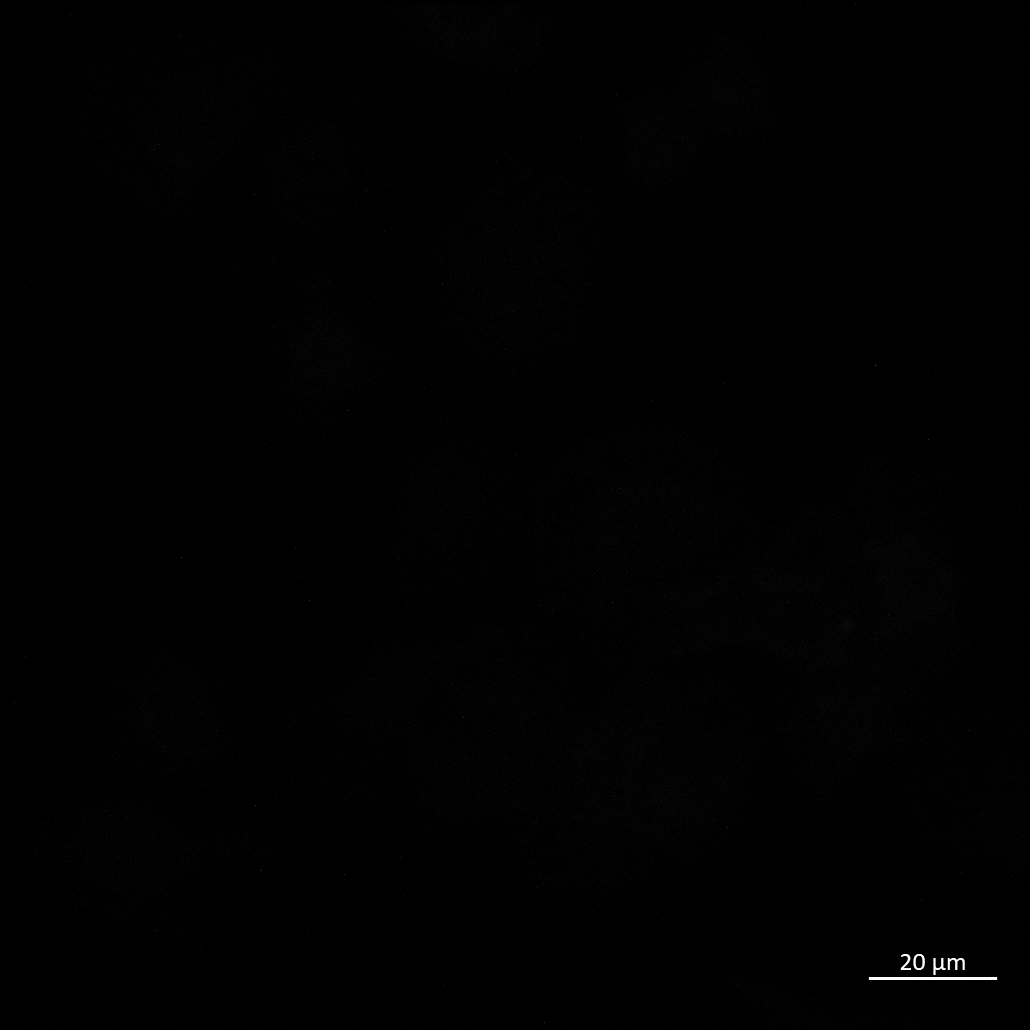

### Untitled8640x_1532C_DMSO_10UMKBU_2hrs_activated beta1(green)_actin(red)_DMSO_02_c1_ORG.tif

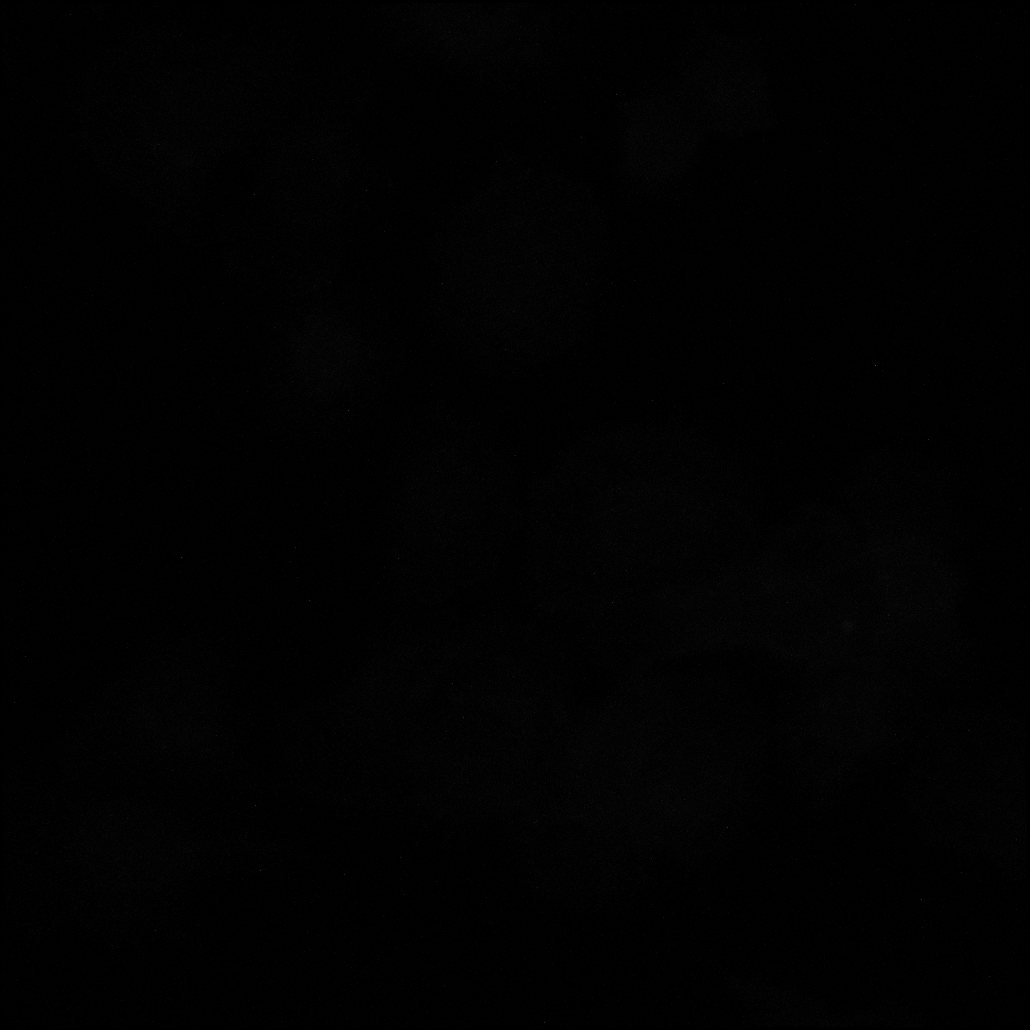

### Untitled8640x_1532C_DMSO_10UMKBU_2hrs_activated beta1(green)_actin(red)_DMSO_02_c2.tif

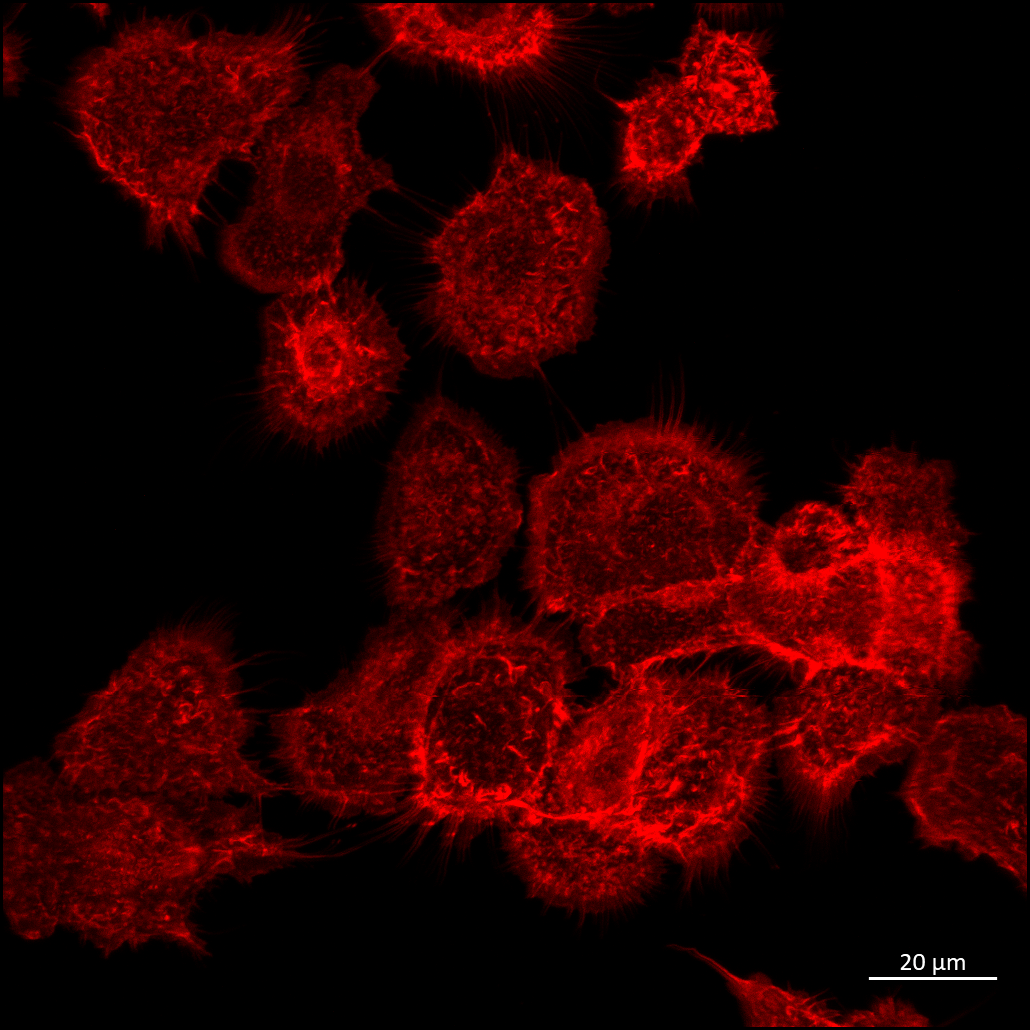

### Untitled8640x_1532C_DMSO_10UMKBU_2hrs_activated beta1(green)_actin(red)_DMSO_02_c2_ORG.tif

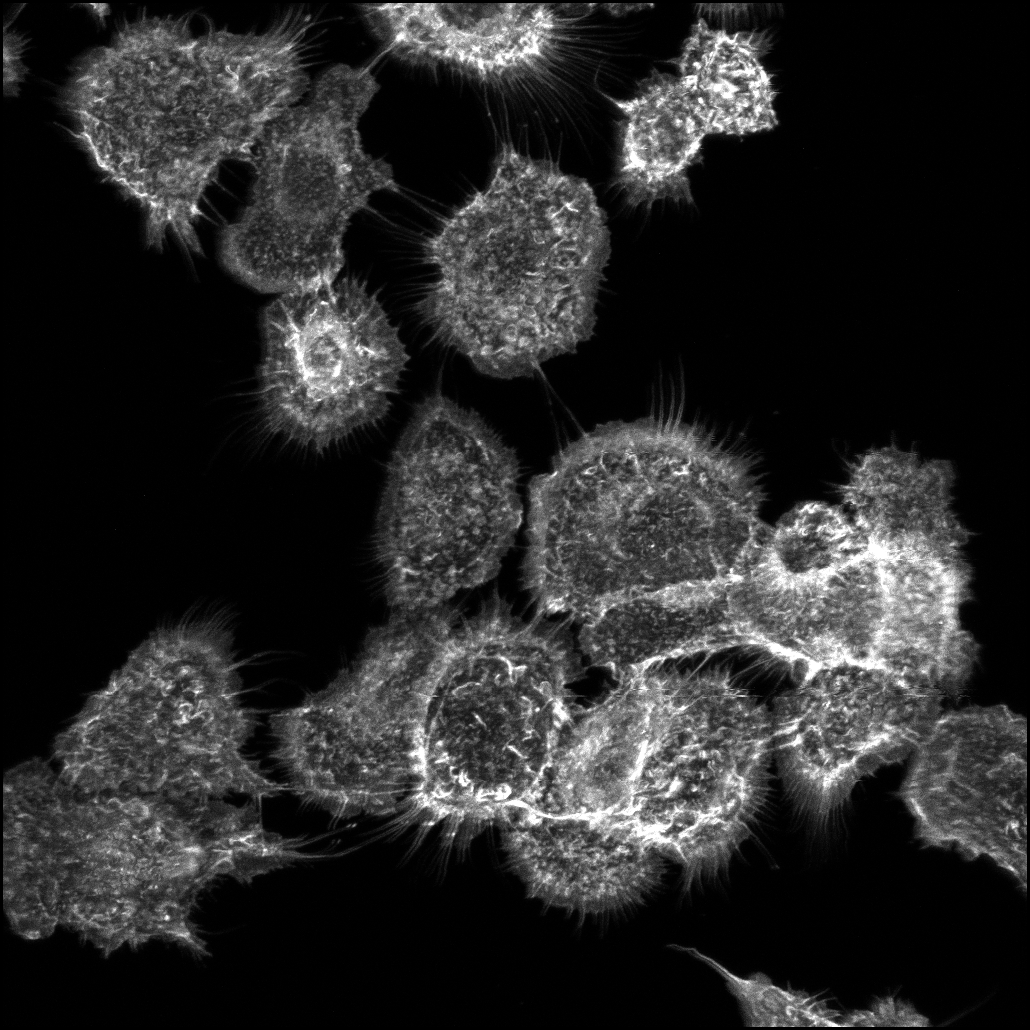

### Untitled8640x_1532C_DMSO_10UMKBU_2hrs_activated beta1(green)_actin(red)_DMSO_02_c3.tif

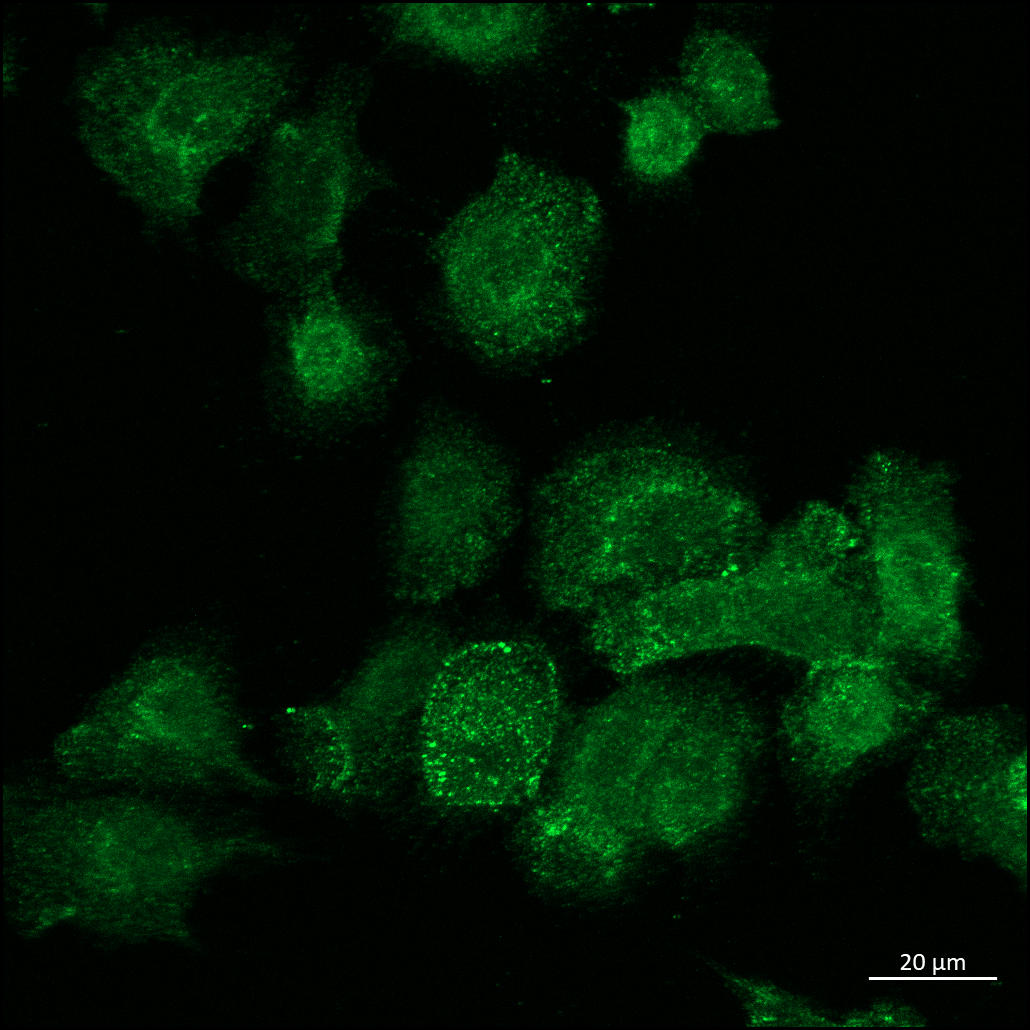

### Untitled8640x_1532C_DMSO_10UMKBU_2hrs_activated beta1(green)_actin(red)_DMSO_02_c3_ORG.tif

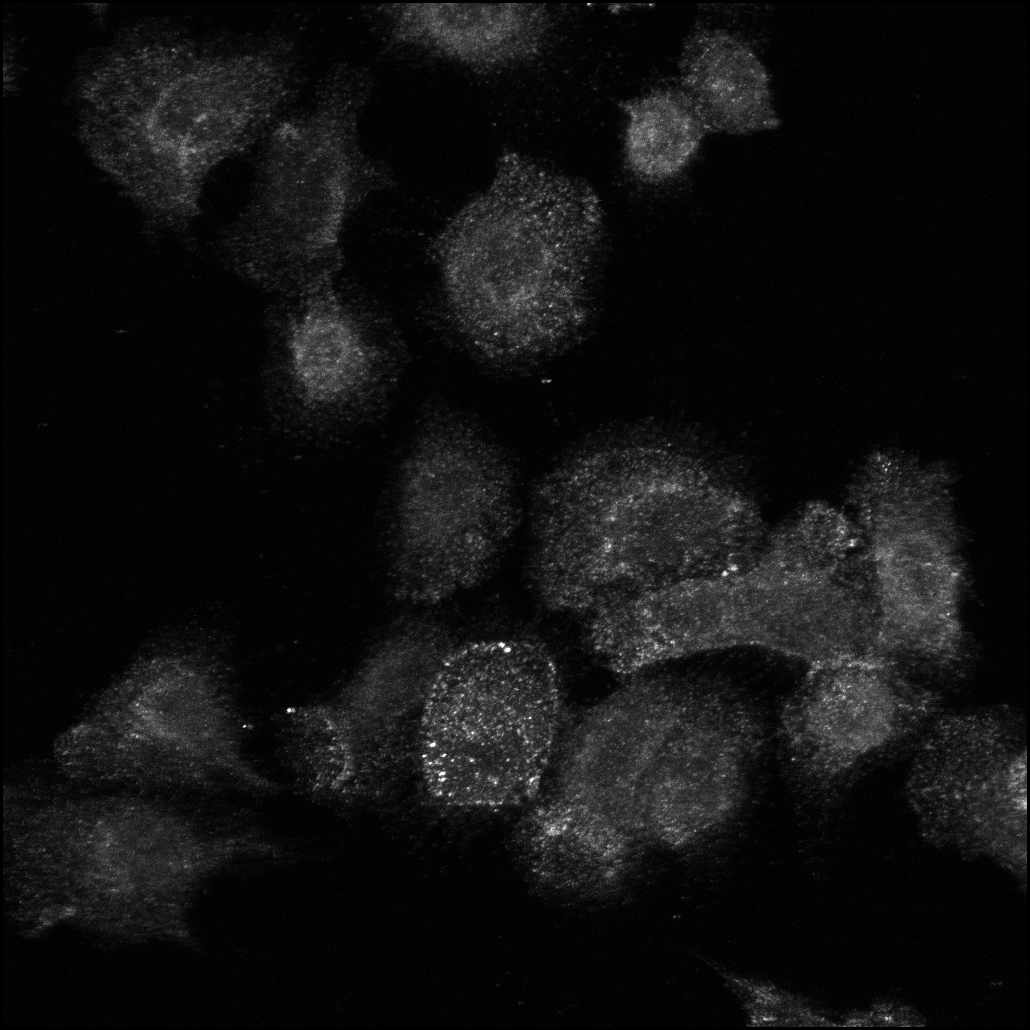

### Untitled8640x_1532C_DMSO_10UMKBU_2hrs_activated beta1(green)_actin(red)_DMSO_02_c4.tif

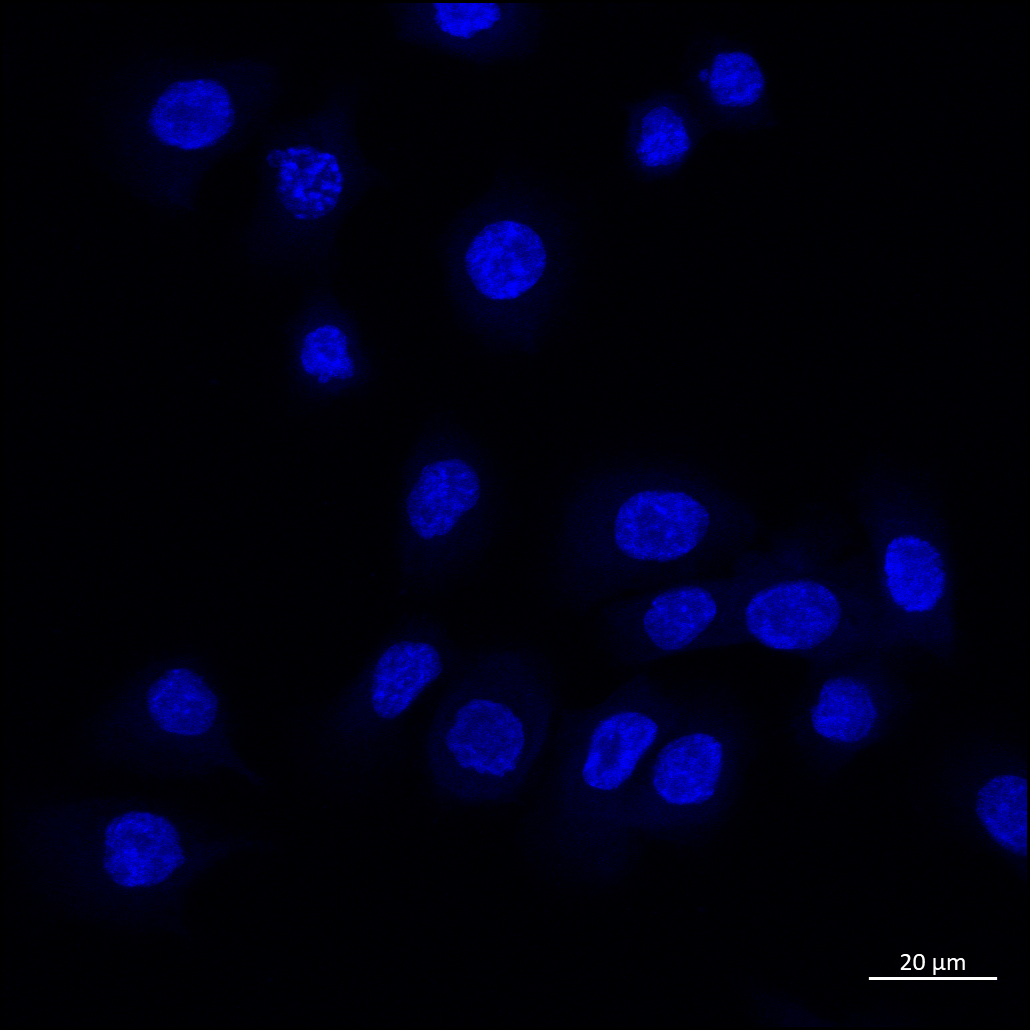

### Untitled8640x_1532C_DMSO_10UMKBU_2hrs_activated beta1(green)_actin(red)_DMSO_02_c4_ORG.tif

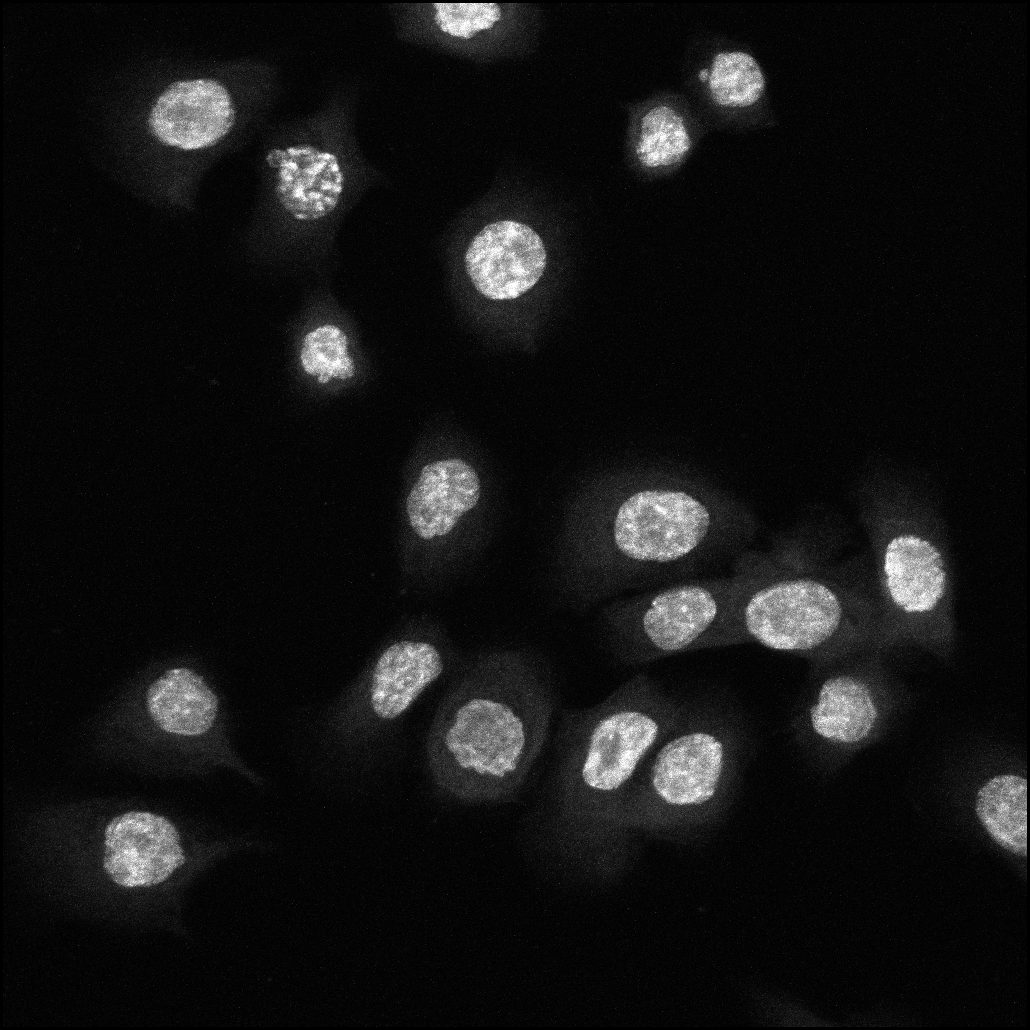

### upstream analysis_Cutoff1_4h_DiffCon_(visualize BH p-value)_expanded header.png

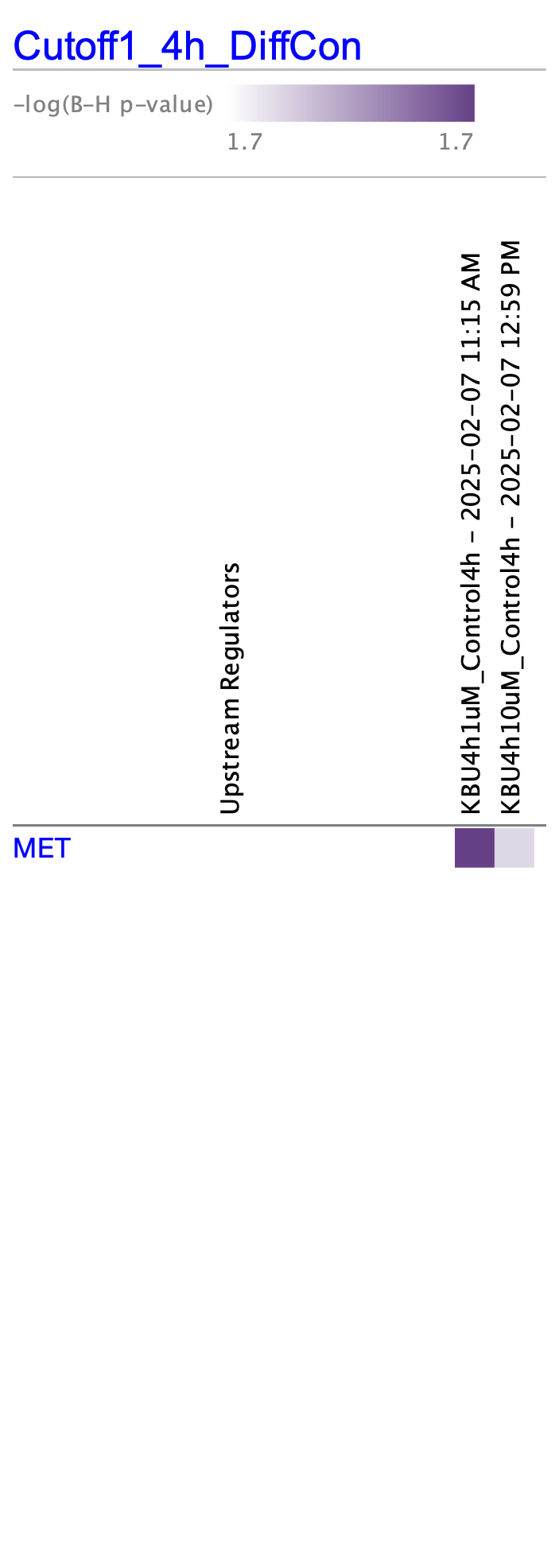

### upstream analysis_Cutoff1_4h_DiffCon_(visualize BH p-value)_not expanded header.png

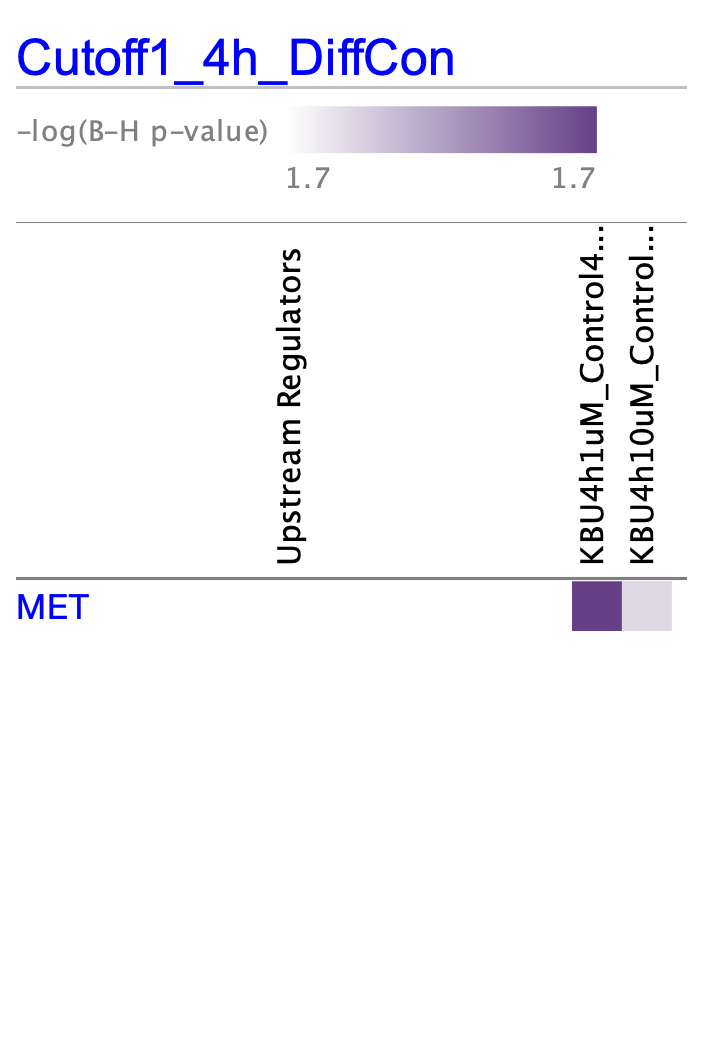

### upstream analysis_Cutoff1_4h_DiffCon_(visualize BH p-value)_not expanded header.png

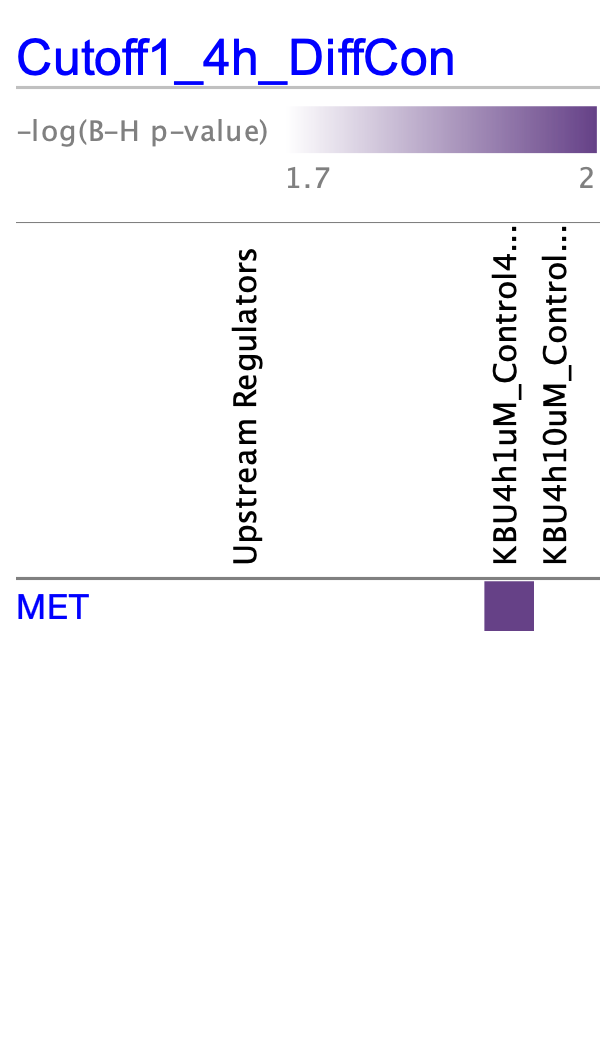

### Volcano_MembraneTMT_HighStringency.pdf

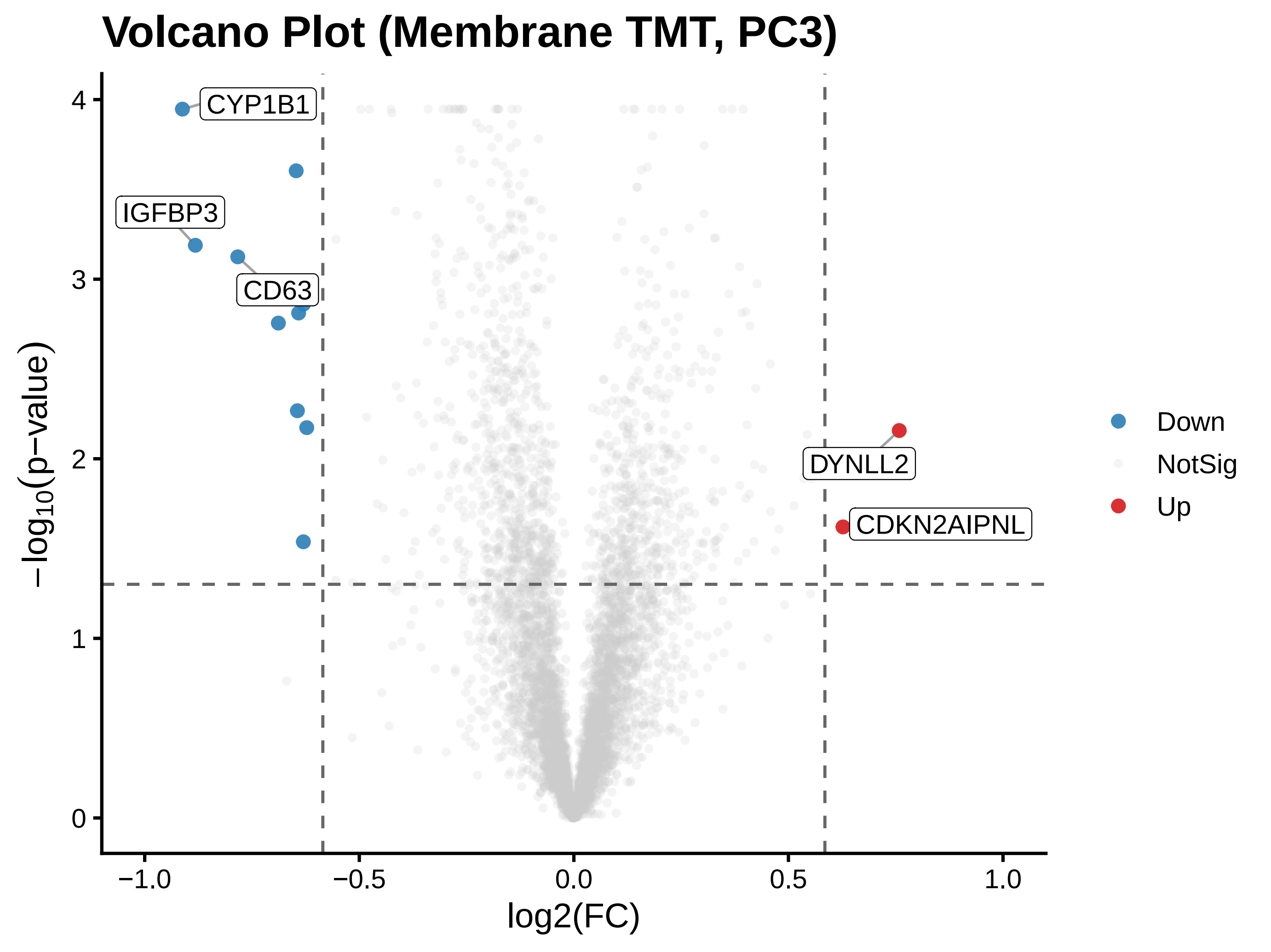

### Volcano_MembraneTMT_HighStringency.png

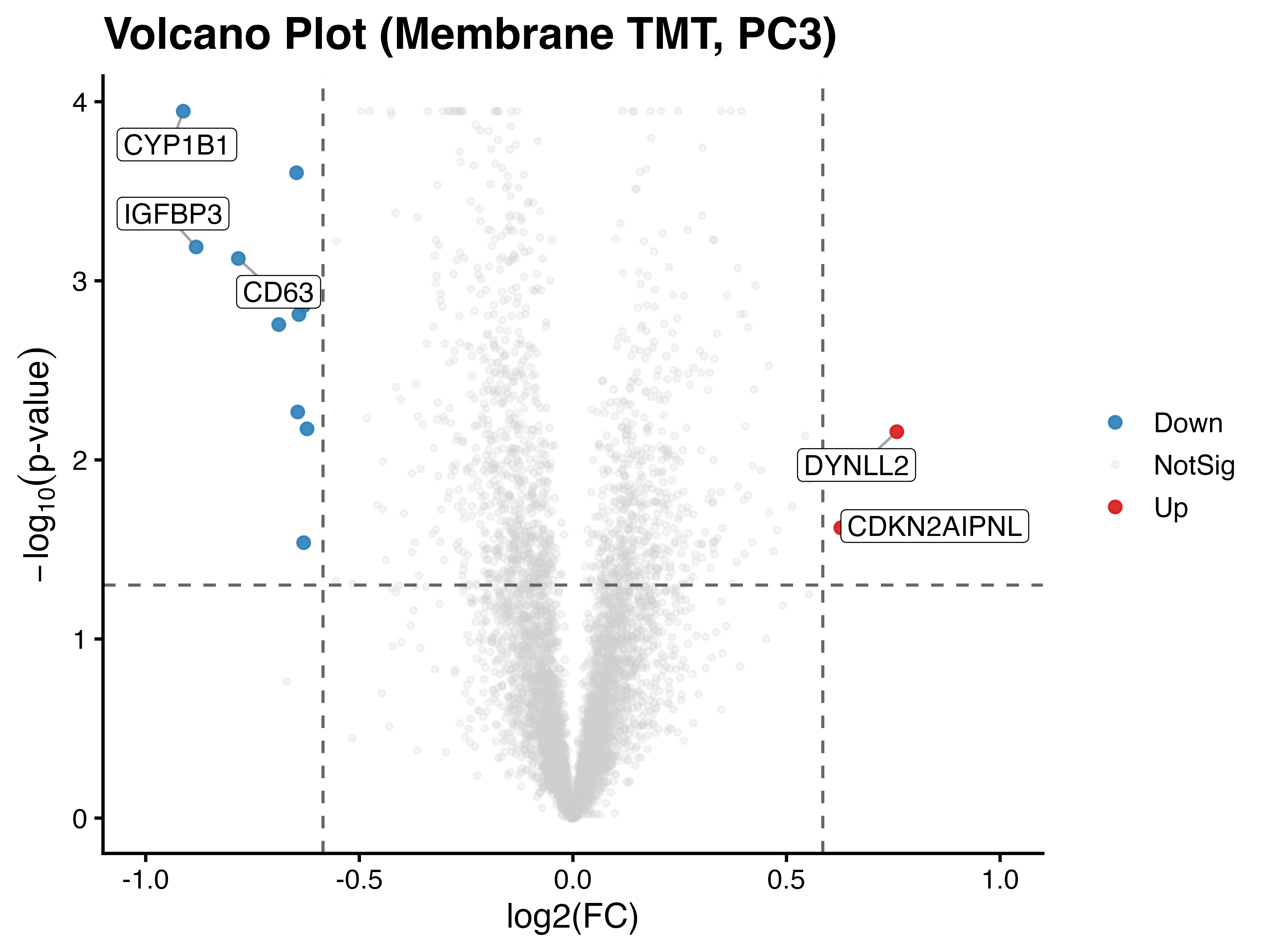

### Volcano_MembraneTMT_Standard.pdf

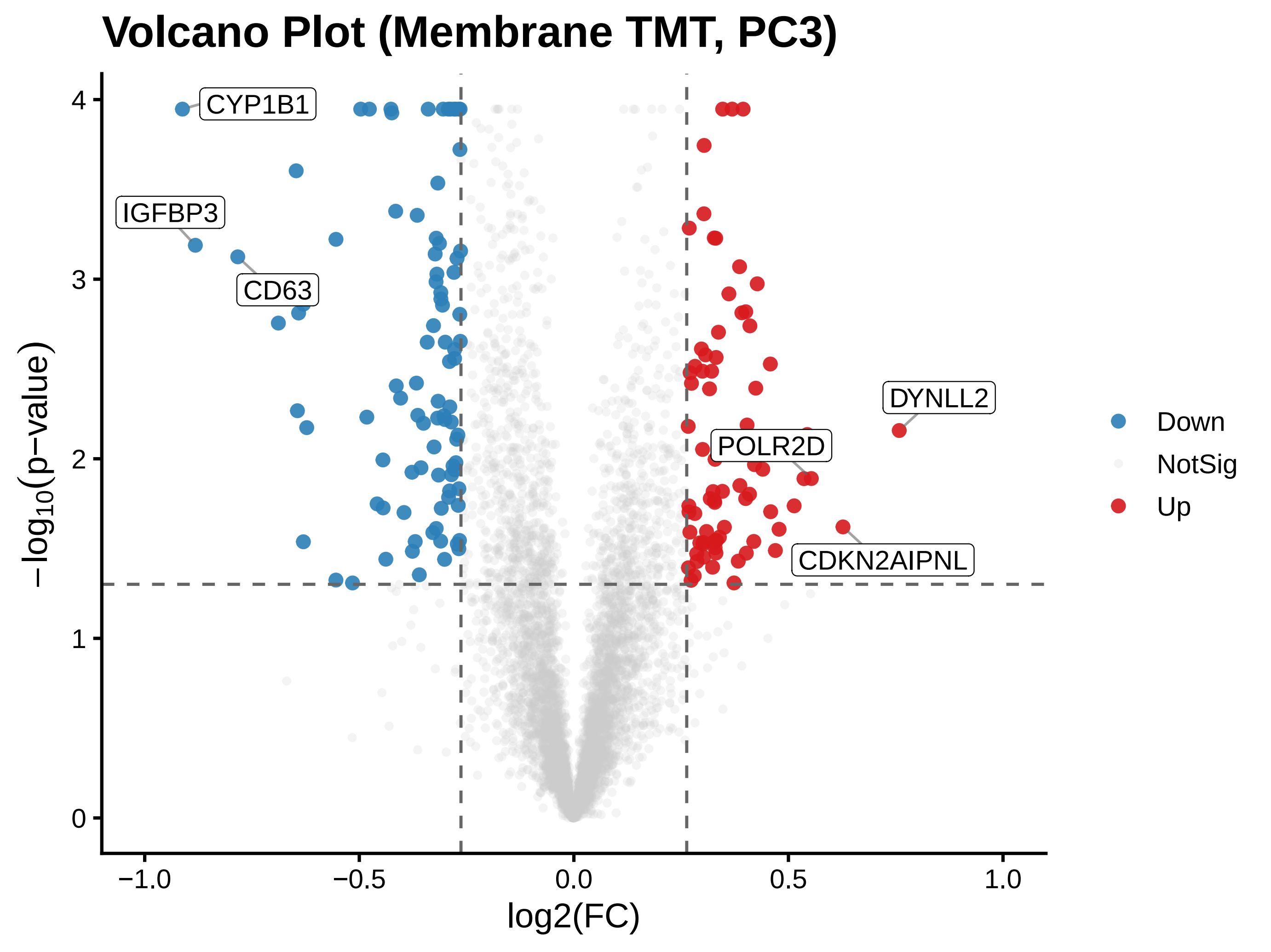

### Volcano_MembraneTMT_Standard.png

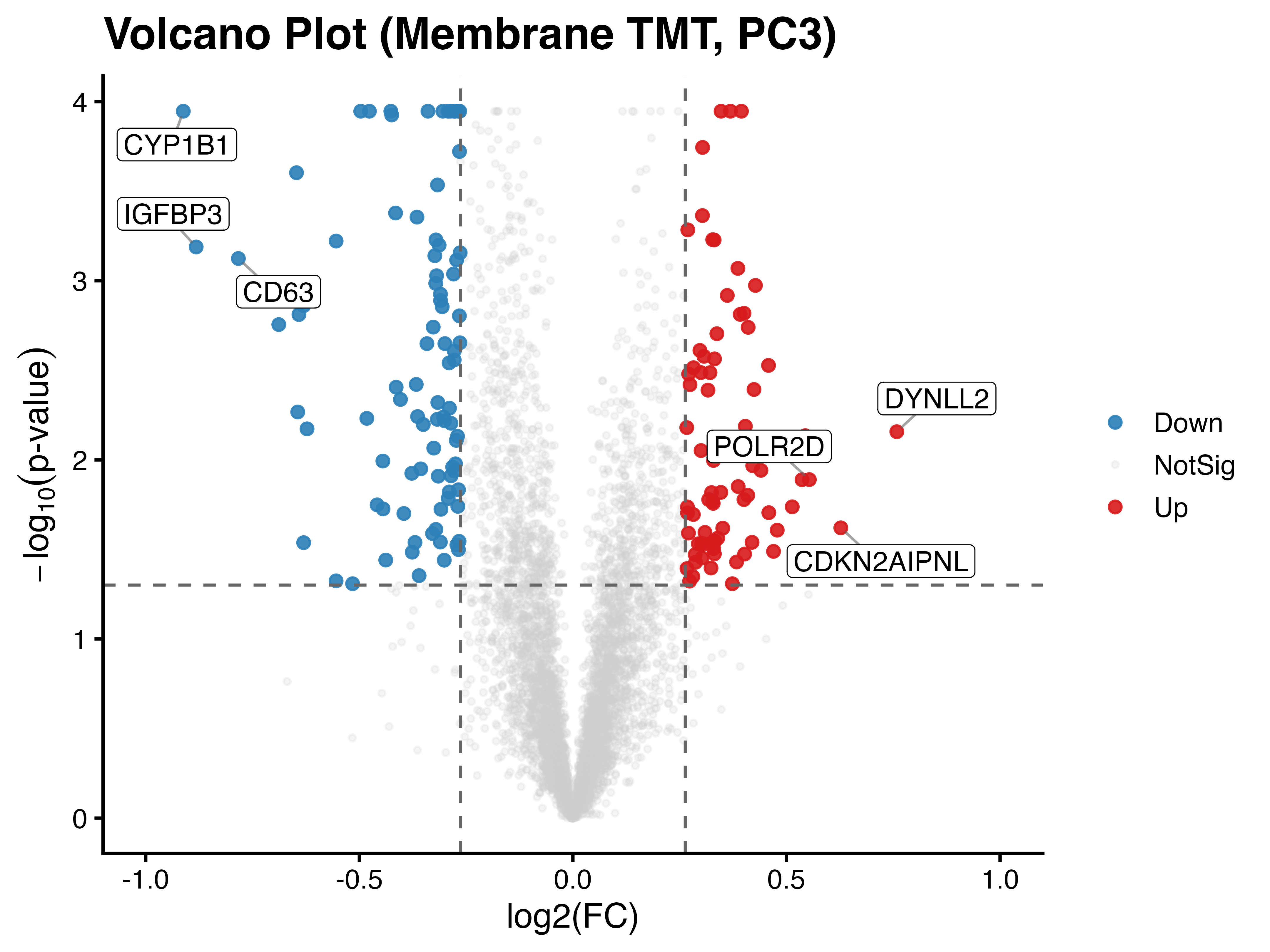

### Volcano_PC3_72h_publication_HQ.pdf

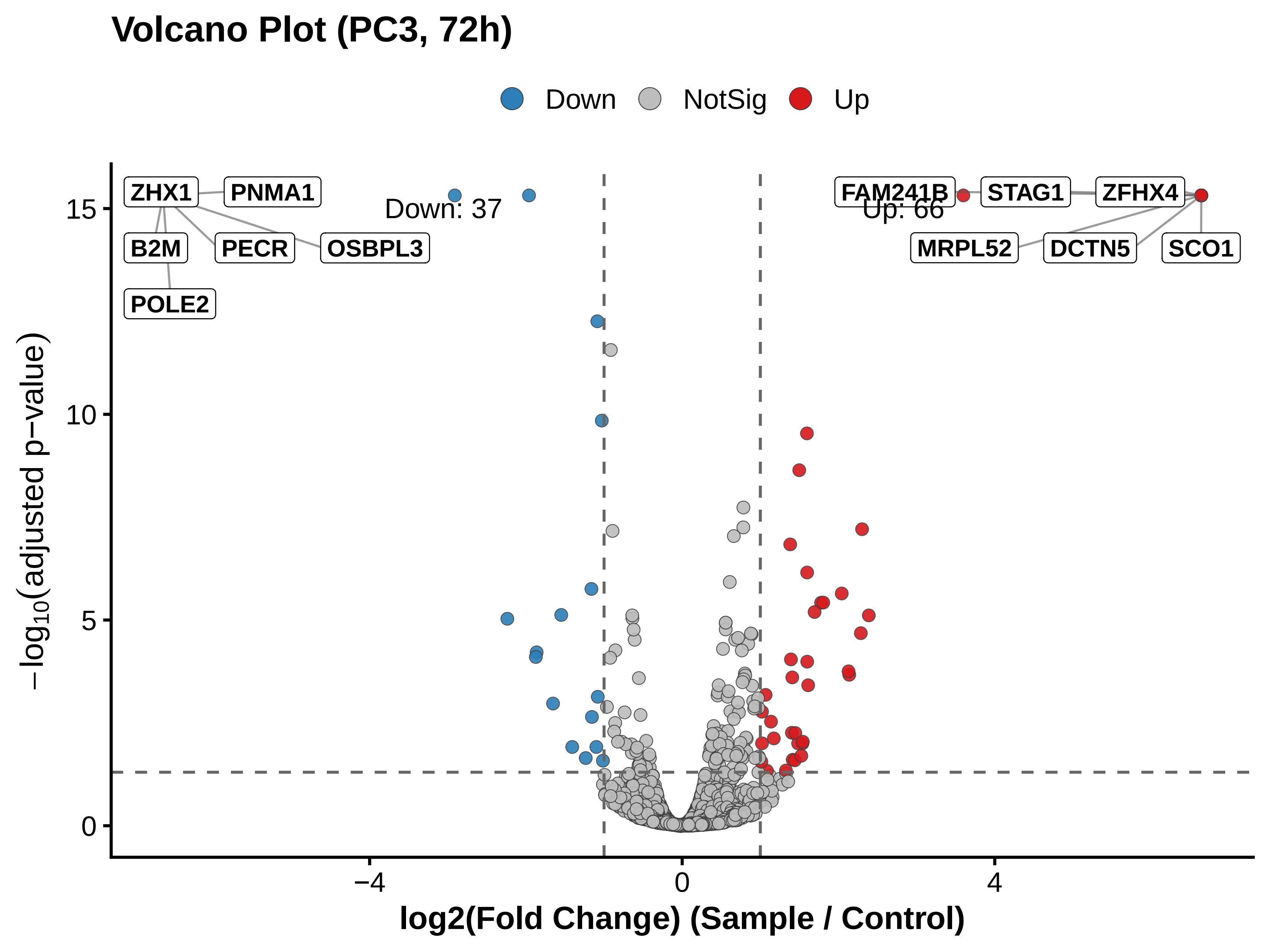

### Volcano_PC3_72h_publication_HQ.png

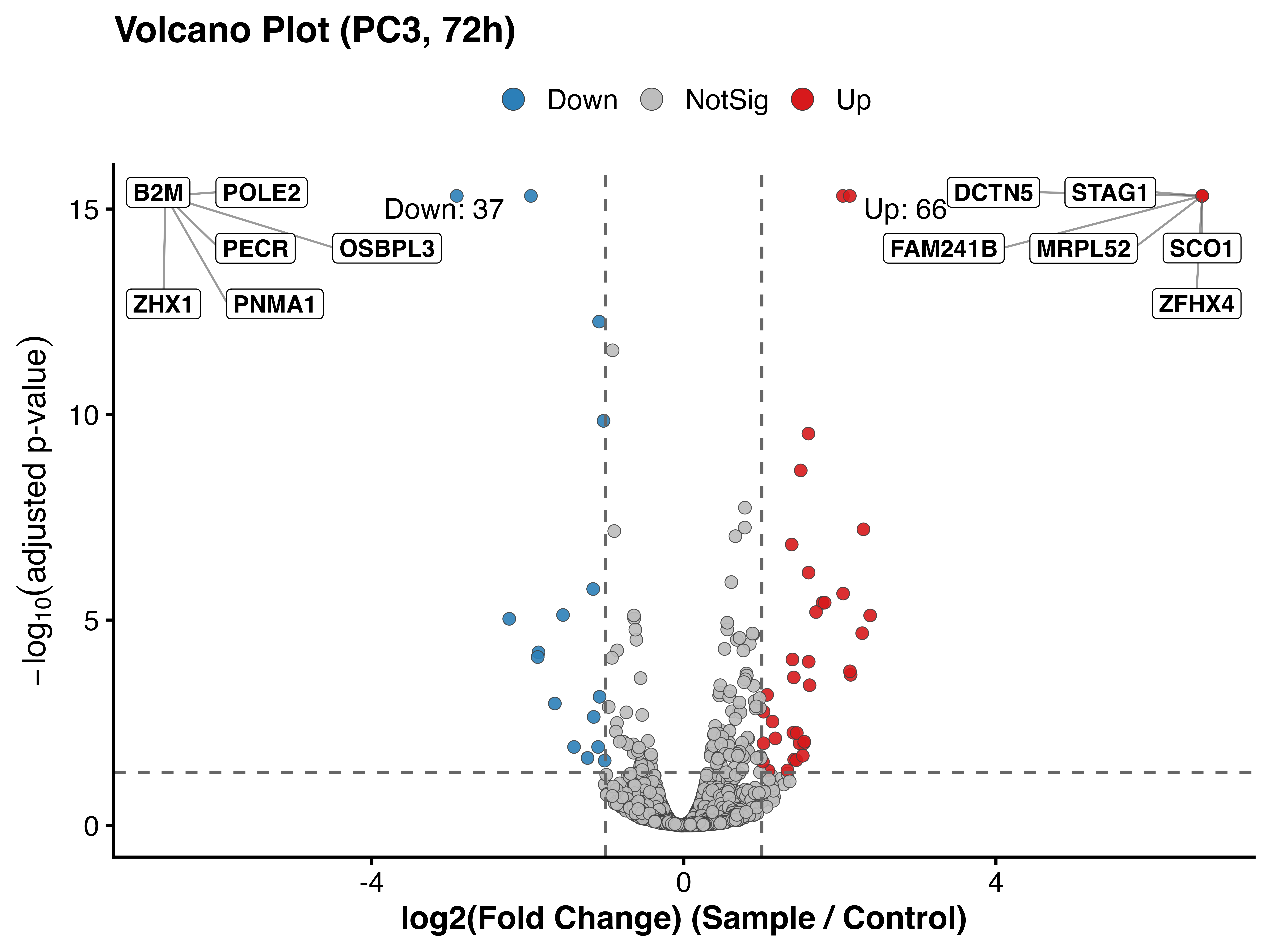
